# Genetic variants associated with cell-type-specific intra-individual gene expression variability reveal new mechanisms of genome regulation

**DOI:** 10.1101/2024.05.05.592598

**Authors:** Angli Xue, Seyhan Yazar, José Alquicira-Hernández, Anna S E Cuomo, Anne Senabouth, Gracie Gordon, Pooja Kathail, Chun Jimme Ye, Alex W. Hewitt, Joseph E. Powell

## Abstract

Gene expression levels can vary substantially across cells, even in a seemingly homogeneous cell population. Identifying the relationships between genetic variation and gene expression is critical for understanding the mechanisms of genome regulation. However, the genetic control of gene expression variability among the cells *within* individuals has yet to be extensively examined. This is primarily due to the statistical challenges, such as the need for sufficiently powered cohorts and adjusting mean-variance dependence. Here, we introduce MEOTIVE (Mapping genetic Effects On inTra-Individual Variability of gene Expression), a novel statistical framework to identify genetic effects on the gene expression variability (sc-veQTL) accounting for the mean-variance dependence. Using single-cell RNA-seq data of 1.2 million peripheral blood mononuclear cells from 980 human donors, we identified 14 – 3,488 genes with significant sc-veQTLs (study-wide *q*-value < 0.05) across different blood cell types, 2,103 of which were shared across more than one cell type. We further detected 55 SNP-gene pairs (in 34 unique genes) by directly linking genetic variations with gene expression dispersion (sc-deQTL) regardless of mean-variance dependence, and these genes were enriched in biological processes relevant to immune response and viral infection. An example is rs1131017 (*p*<9.08×10^−52^), a sc-veQTL in the 5’UTR of *RPS26*, which shows a ubiquitous dispersion effect across cell types, with higher dispersion levels associated with lower auto-immune disease risk, including rheumatoid arthritis and type 1 diabetes. Another example is *LYZ*, which is associated with antibacterial activity against bacterial species and was only detected with a monocyte-specific deQTL (rs1384) located at the 3’ UTR region (*p*=1.48×10^−11^) and replicated in an independent cohort. Our results demonstrate an efficient and robust statistical method to identify genetic effects on gene expression variability and how these associations and their involved pathways confer auto-immune disease risk. This analytical framework provides a new approach to unravelling the genetic regulation of gene expression at the single-cell resolution, advancing our understanding of complex biological processes.

## Introduction

Dissecting the genetic control of gene expression is necessary to understand the biological mechanisms of genome regulation. With the development of single-cell sequencing technology, many studies have sought to identify how genetic effects underlying gene expression act at the level of individual cells. These single-cell expression Quantitative Trait Loci (sc-eQTL) have revealed that the genetic effects underlying mean differences in gene expression between individuals are frequently cell-type specific^1–7^. These sc-eQTL analyses are based on the model assumption that allelic alternatives have an additive effect on the mean expression levels of RNA amongst genotype groups. They are typically tested using linear regression between an SNP’s genotype and the mean gene expression levels among a population. Variance QTL (vQTL) are different from eQTL in that genotypes are associated with the variation of the phenotype (Figure 1A). They have been studied for complex human traits such as body mass index (BMI)^8–12^, bone marrow density (BMD)^13^, vitamin D^14^, glycemic traits^15^, and serum cardiometabolic biomarkers^16^. However, only a few studies^17–20^ have investigated the characteristics of variance eQTL (veQTL), i.e., the eQTL that affects the gene expression variation in each genotype group. They have identified inter-individual veQTLs (genetic effects on variance across individual expression levels) and linked them to G x G effects, such as *cis*-*epistatic* interactions^18^ and interaction with a second gene’s expression level^19^. They can also be caused by gene-environment (G x E) interaction effects. Most of these analyses are based on a variance test, examining the phenotypic variability among genotype groups, such as Levene’s^21^ and Brown-Forsythe tests^22^.

**Figure 1.**
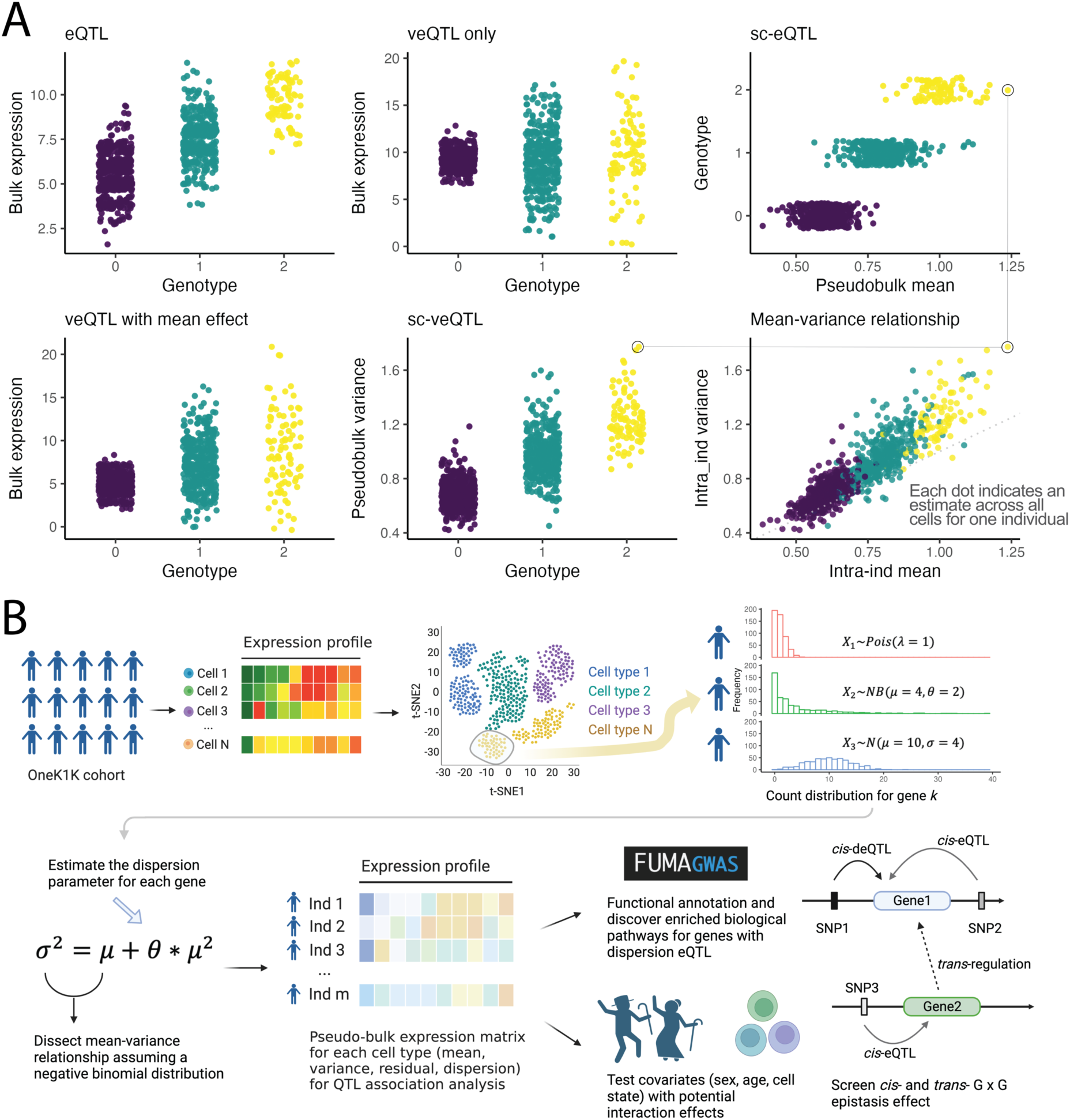
The illustration of variance eQTL and overview of the study design. **(A)** Illustrative plot to explain the bulk eQTL/veQTL and single-cell eQTL/veQTL. In bulk-level analysis, each dot indicates the gene expression count for one individual. In single-cell level analysis, each dot indicates the pseudobulk mean/variance expression for one individual in a specific cell type. **(B)** The analytical pipeline of MEOTIVE framework identifying genetic variants that affect intra-individual gene expression variability.

An under-investigated phenomenon is intra-individual veQTL – the relationship between genetic variants and gene expression variability within an individual. This type of eQTL is defined as the genetic variants that affect gene expression variability across cells within an individual between different genotype groups (**Figure 1A**). Such variability in a population of cells (or within a pre-defined cell type) contains the information of cell-to-cell heterogeneity. It may arise from genetic regulation, immune response, cell cycle, cell state, and/or the stochasticity of cellular gene expression. One of the few empirical studies to tackle this question tested for intra-veQTLs (veQTLs hereafter for brevity unless otherwise specified) in sc-RNA-seq data from 5,447 induced Pluripotent Stem Cells (iPSCs) collected across 53 individuals. They identified five study-wide significant sc-veQTL, but demonstrated that the differences in variance were induced just by statistical dependence rather than true biological effects^23^. As the authors identified, sample size and number of cells limited the discovery of sc-veQTL.

Intra-individual variation is particularly interesting because cell-to-cell heterogeneity in the gene expression (especially in immune cells) within an individual might result from important functional consequences, i.e., genetic regulatory mechanism on gene expression during development and homeostasis, as well as a potential response to therapies, and pathological development (such as tumour growth). For example, a recent study^24^, collecting peripheral blood mononuclear cells (PBMCs) from asthma patients, has linked sc-veQTL to the transcriptional response to immune stimuli and glucocorticoids. In addition, intra-individual variation could be a potential indicator for the G x E interaction with cellular traits such as the cell cycle and cell state at the single-cell level.

Several challenges are key to identifying sc-veQTLs and understanding their characteristics. First, we need to accurately estimate the intra-individual variance from the scRNA-seq data, where expression counts across cells are typically assumed to follow either Poisson, negative binomial (NB), or zero-inflated negative binomial (ZINB) distribution, especially when the number of cells per sample is low. Second, we need to account for the mean-variance relationship as they are mathematically correlated under those aforementioned models. For example, assuming the gene expression level of a gene across cells follows a negative binomial distribution, the variance (σ^2^) can be estimated such that σ^2^ = μ + μ^2^ ∗ θ, where μ is the mean and θ is the dispersion of the distribution. In this scenario, sc-veQTLs could be primarily induced by differences in the mean expression levels between individuals of different genotype groups. Third, the selection of appropriate statistical tests for detecting sc-veQTLs is uncertain. This may depend on the distributions of intra-individual variance/dispersion estimates across the cohort, and such distributions are likely to vary between genes and cell types.

Here, we introduce MEOTIVE (**M**apping genetic **E**ffects **O**n in**T**ra-**I**ndividual **V**ariability of gene **E**xpression), a novel statistical framework to identify the relationship between genetic variants and within-individual variability in the expression levels of a gene at single-cell resolution. Leveraging data from the OneK1K cohort^6^, comprising genotype and scRNA-seq data from 1.27 million PBMCs collected across 980 individuals, we test genetic variants’ role on the variance of intra-individual gene expression at the population level while accounting for the mean-variance relationship (**Figure 1B**). Considering the statistical challenges, we decide to use two methods in parallel to identify the sc-veQTL that are not induced by mean effect (ME): (1) a model-free strategy to detect those significant sc-veQTLs that do not show significant results in the sc-eQTL association analysis and (2) a model-based strategy to first estimate the intra-individual dispersion parameter (θ) of the expression and then perform the dispersion-eQTL (sc-deQTL) association analysis. We also investigate how sc-deQTLs show heterogeneity underlying different cell states and seek potential biological explanations for them. Overall, we provide a statistical framework to identify single-cell variance eQTL and reveal novel mechanisms of how genetic variants affect gene expression variability at the single-cell level.

## Results

To understand the relationship between genetic loci and intra-individual variation in single-cell gene expression, we began by exploring the estimation of data parameters and their relationship in sc-eQTL models. This is important due to: (1) previously identified relationships between estimates such as mean and variance^25–27^, and (2) the potential impact of differences in the distribution of a gene’s expression levels, with genes varying between zero-inflated negative binomial through to Gaussian distributions.

## The relationship between mean and intra-individual variance effects

Identifying genetic effects on variance is attractive as they can be interpreted easily. However, as previously described, the mean difference between individuals can be correlated with intra-individual variance effects^23^. Nevertheless, variance effects can exist independently, with the most likely explanation that they are ‘true’ genetic effects arising from genotype by genotype (GxG) or genotype by environment (GxE) interactions. Using TensorQTL^28^, we tested for intra-individual sc-veQTL per cell type, adjusting for sex, age, and latent variables (**Methods**). In total, we identified 4,642 significant (*q*-value < 0.05) vGenes across all 14 cell types (10,527 unique SNP-gene pairs), ranging from 14 to 3,488 for each cell type. As expected, the intra-individual mean estimates are highly correlated with variance estimates for all cell types (**Figure 2A**), and the correlation is highly dependent on the mean estimates and proportion of cells with zero expression (**Figure 2B-C**). Testing for the overlap with mean eQTL effects, we observe 94.5% of vGenes also have mean effects (eGenes), and conversely, 60.4% of eGenes also have a variance effect (**Table 1** and **Figure 2D**). The results suggest a small number of sc-veQTL may have variance-only effects and are worth further investigation. We evaluated the effect size, significant level, and TSS distance between vGenes with or without mean effects and the comparison showed the latter group is merely above the significant threshold (*q*-value = 0.05), suggesting this group of signals are more likely to be random noises due to arbitrary threshold (**Supplementary Figures 1-3** and **Methods**). The number of vGenes identified per cell type is a function of the sample size (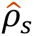 = 0.93) and average number of cells per donor (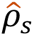 = 0.98) (**Figure 2E-F**), an observation also seen with eGenes (**Table 1**). Furthermore, sc-veQTL and sc-eQTL allelic effects are highly correlated (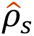 = 0.942∼0.991), confirming the relationship between mean and variance effects (**Figure 3** and **Supplementary Figure 4**). Finally, for the vGenes that are also eGenes, all the top veQTLs have the same direction of allelic effect, with *RPS15A* the only exception, but the mean effect of the top veQTL (rs7193785) is not significant (nominal *p*-value = 0.335). These results confirm the strong correlation between the intra-individual mean and variance (**Figure 2A** and **Supplementary** Figures 5), verifying that mean veQTLs can explain most effects.

**Figure 2.**
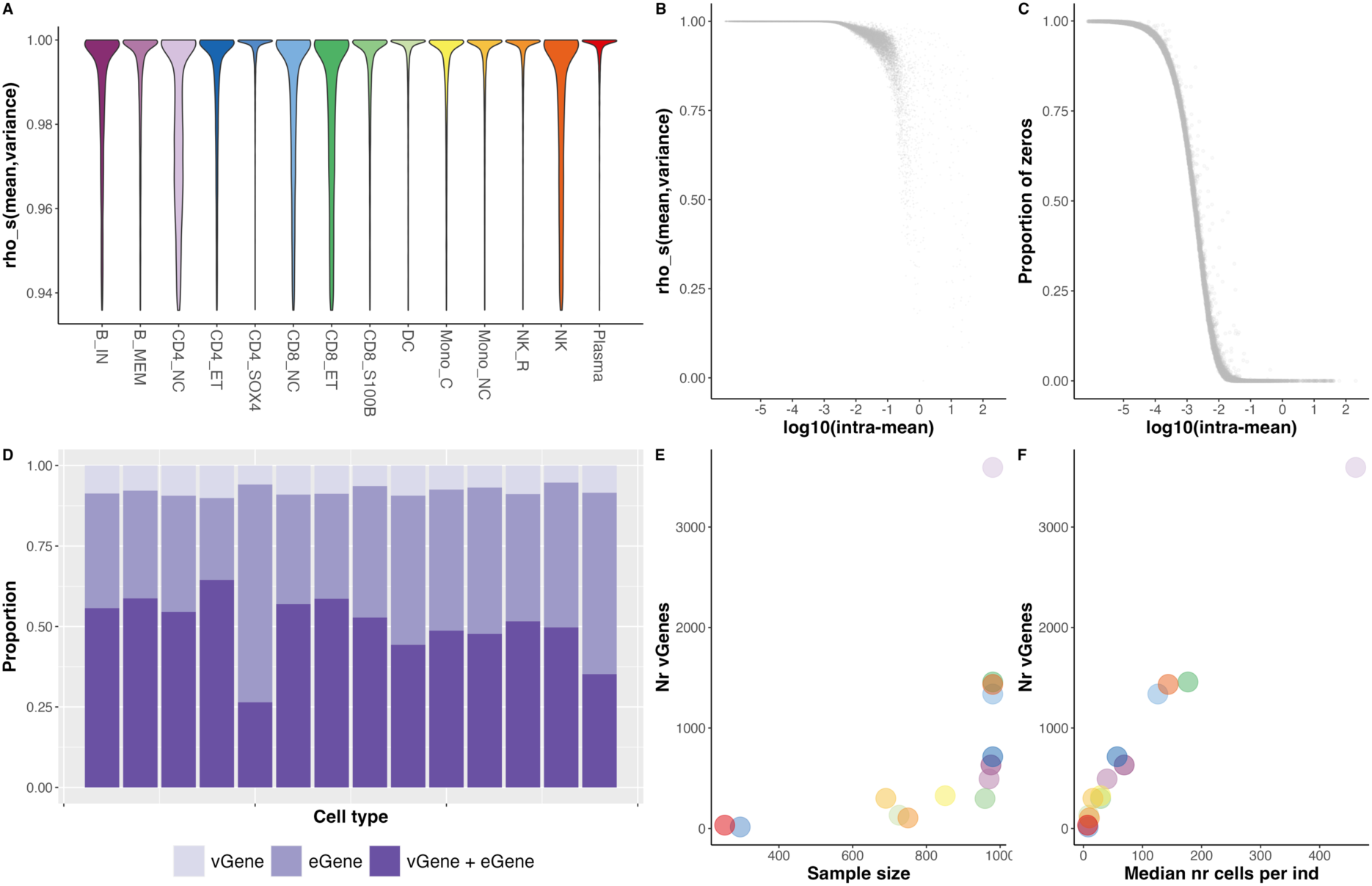
The intra-individual mean-variance relationship and overlap between eGenes and vGenes per cell type. **(A)** The Spearman’s correlation estimates between the intra-individual mean and variance of each gene. The colour of the violin plot denotes the corresponding cell type. The bottom 10% correlation estimates are omitted. All violins have the same maximum width. **(B)** The relationship between intra-individual mean and mean-variance correlation per gene in CD4 naïve cells. **(C)** The relationship between intra-individual mean and proportion of no-expression individuals per gene in CD4 naïve cells. **(D)** The percentage of eGene, vGene, and dGene in each cell type. **(E)** The relationship between sample size and number of vGenes. **(F)** The relationship between median number of cells per individual and number of vGenes. Each dot represents a cell type, and the colour of the dots corresponds to panel.

**Figure 3.**
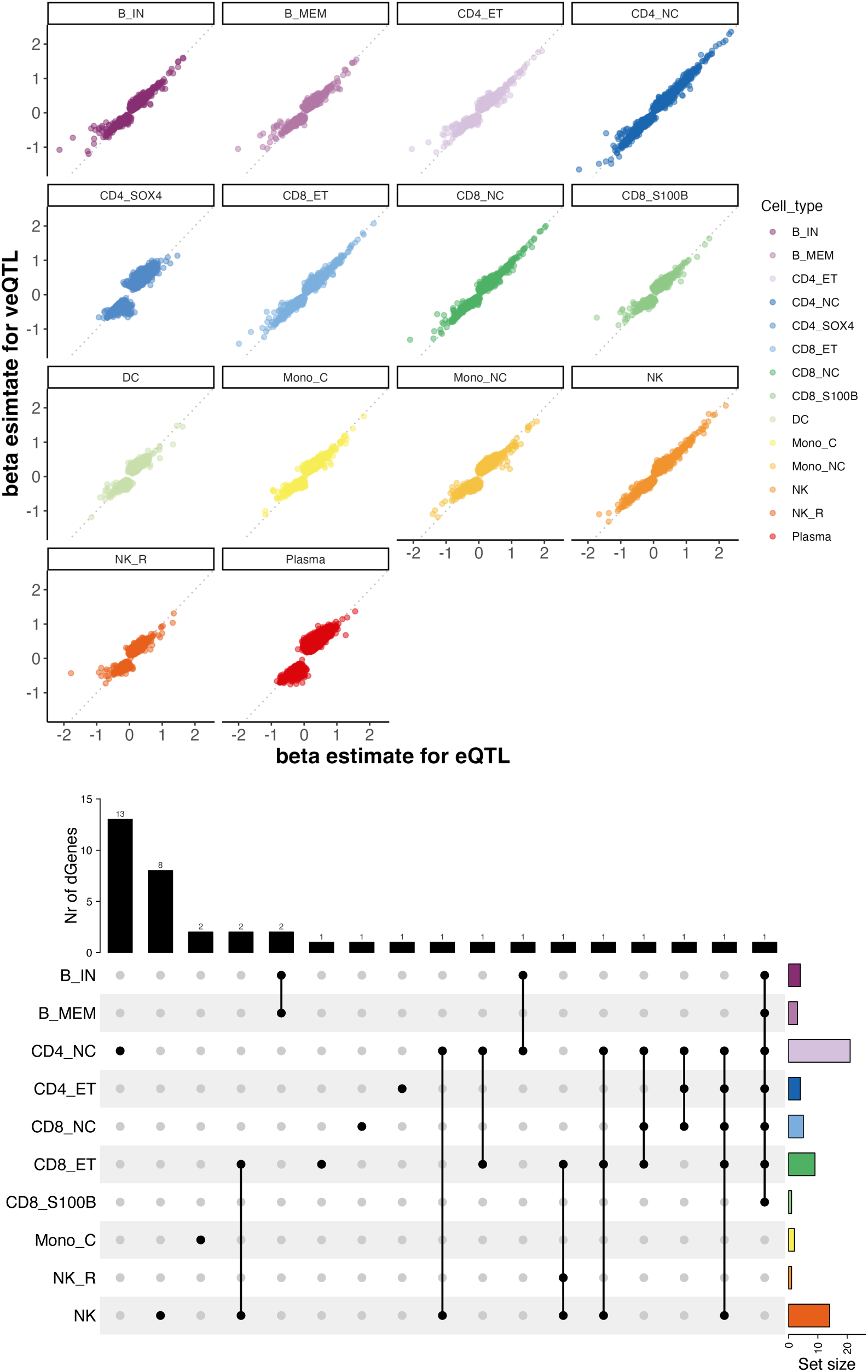
Effect size comparison among the mean, variance, and dispersion eQTLs. (**A**) The x-axis and y-axis denote the beta estimate for eQTL and veQTL, respectively. Each dot indicates an SNP-gene pair test, and the dot’s colour indicates the cell type. The grey dashed line denotes the diagonal line of the coordinate panel. (**B**) An Upset plot for the number of dGenes in each cell type and the dGenes shared across cell types. The number of each intersection was annotated as the number above the bar plot.

**Table 1.** Summary of eGene and vGene identified in 14 cell types.

| Cell type | N | Median<br>nr cells | Nr<br>genes | eGene | vGene | overlap | vGene<br>with<br>ME | eGene<br>with<br>VE |
| --- | --- | --- | --- | --- | --- | --- | --- | --- |
| B_IN | 975 | 69 | 12219 | 843 | 605 | 574 | 0.949 | 0.681 |
| B_MEM | 970 | 40 | 11663 | 672 | 480 | 454 | 0.946 | 0.676 |
| CD4_NC | 980 | 461 | 15433 | 4252 | 3488 | 3363 | 0.964 | 0.791 |
| CD4_ET | 980 | 57 | 12040 | 986 | 692 | 655 | 0.947 | 0.664 |
| CD4_SOX4 | 295 | 8 | 8234 | 37 | 14 | 13 | 0.929 | 0.351 |
| CD8_NC | 980 | 126 | 13508 | 1760 | 1263 | 1213 | 0.960 | 0.689 |
| CD8_ET | 980 | 177 | 13798 | 1884 | 1369 | 1314 | 0.960 | 0.697 |
| CD8_S100B | 959 | 29 | 11070 | 461 | 295 | 271 | 0.919 | 0.588 |
| DC | 726 | 9 | 10149 | 228 | 127 | 118 | 0.929 | 0.518 |
| Mono_C | 851 | 29 | 11620 | 517 | 319 | 305 | 0.956 | 0.590 |
| Mono_NC | 690 | 16 | 10544 | 497 | 297 | 284 | 0.956 | 0.571 |
| NK_R | 750 | 10 | 9224 | 179 | 105 | 98 | 0.933 | 0.547 |
| NK | 980 | 143.5 | 13577 | 2099 | 1440 | 1360 | 0.944 | 0.648 |
| Plasma | 253 | 7 | 8983 | 70 | 33 | 31 | 0.939 | 0.443 |

Notes: N, sample size; Median nr cells, the median number of cells per individual; Nr genes, number of genes tested in the QTL analysis; eGene, number of significant eGene; vGene, number of significant vGene; overlap, the number of genes that are both eGene and vGene; vGene with ME, the proportion of vGene that are also eGene; eGene with VE, the proportion of eGene that are also vGene.

One explanation is that variance effects occur in addition to mean effects, i.e., the variance effects are greater than those expected under the mean-variance relationship. To investigate this, we tested for genetic effects on the residuals from the regression of the variance on the mean per gene (**Methods**). We identified 307 genes (*q*-value < 0.05) with at least one significant residual-eQTL (**Supplementary Table 1**). However, only seven genes overlap with the vGenes without mean effects. This lack of overlap validates that testing or such residual-eQTL is not valid for identifying true veQTL. Previous studies have attempted other parameterisations to test for genetic effects on intra-individual gene expression variance, including variance-to-mean ratio (VMR, also known as Fano factor) and coefficient of variation (CV)^23^. However, these two metrics are highly dependent on the intra-individual mean differences (**Supplementary** Figures 6-7). Another way to adjust the mean-variance relationship is to fit the mean as a covariate when detecting the veQTL per gene^29^. However, this strategy is expected to be valid only when the mean and variance follow a linear relationship; otherwise, the residuals may have a spurious quadratic relationship with the mean. A recent study^30^ proposed using a single polynomial regression to remove mean-variance dependence, but it might nonetheless ignore the genuine biological relationship between mean and dispersion since it forces the residuals to be independent of the mean. Morgan et al.^31^ used local polynomial fit between the squared CV (i.e., CV^2^) and mean across individuals and tried to identify variability protein-QTLs. Another relevant study^27^ estimates a latent gene-specific residual dispersion parameter by fitting a global trend between mean and over-dispersion estimates for all genes, but this was for differential variability testing between two groups of cells and the method was based on the strong assumption that genes with similar expression level are more likely to have similar dispersion level.

## MEOTIVE: direct estimation of dispersion identifies independent genetic effects in intra-individual single-cell expression variability

To solve the problems associated with the mean-variance relationship, we developed MEOTIVE, a framework that tests for allelic effects on intra-individual expression dispersion levels and demonstrates that effects are independent from mean effects. MEOTIVE starts with directly estimating the dispersion parameter (θ) for each gene’s intra-individual expression distribution using a Cox-Reid adjusted Maximum Likelihood Estimation (CR-MLE) method^32^ (**Methods** and **Supplementary Figure 8**). We subsequently test the genotypes of *cis*-SNPs (< 1Mb) for association with the intra-individual dispersion for each gene per cell type, accounting for the accuracy of parameter estimation (**Methods**). In total, we identified 55 significant (*q*-value < 0.05) deQTL-dGene pairs across 34 unique dGenes (**Table 2**, **Figure 3** and **Supplementary Figure 9**). Most (78%) of the deQTLs have an opposite direction of allelic effect to their corresponding veQTL (**Supplementary Table 2**). This is because the distribution type of gene expression count per genotype group varies. For example, when the expression level is low for a specific genotype group, the distribution of the count data is more left-skewed driving the dispersion to be higher. For heterozygous individuals, the distribution approaches Gaussian, thus the dispersion becomes much smaller, and so on for individuals with alternative homozygous alleles (see example of *RPS26* below, **Figure 4**). This feature results in a negative relationship between intra-individual variance and intra-individual dispersion for many genes with significant veQTLs and deQTLs.

**Figure 4.**
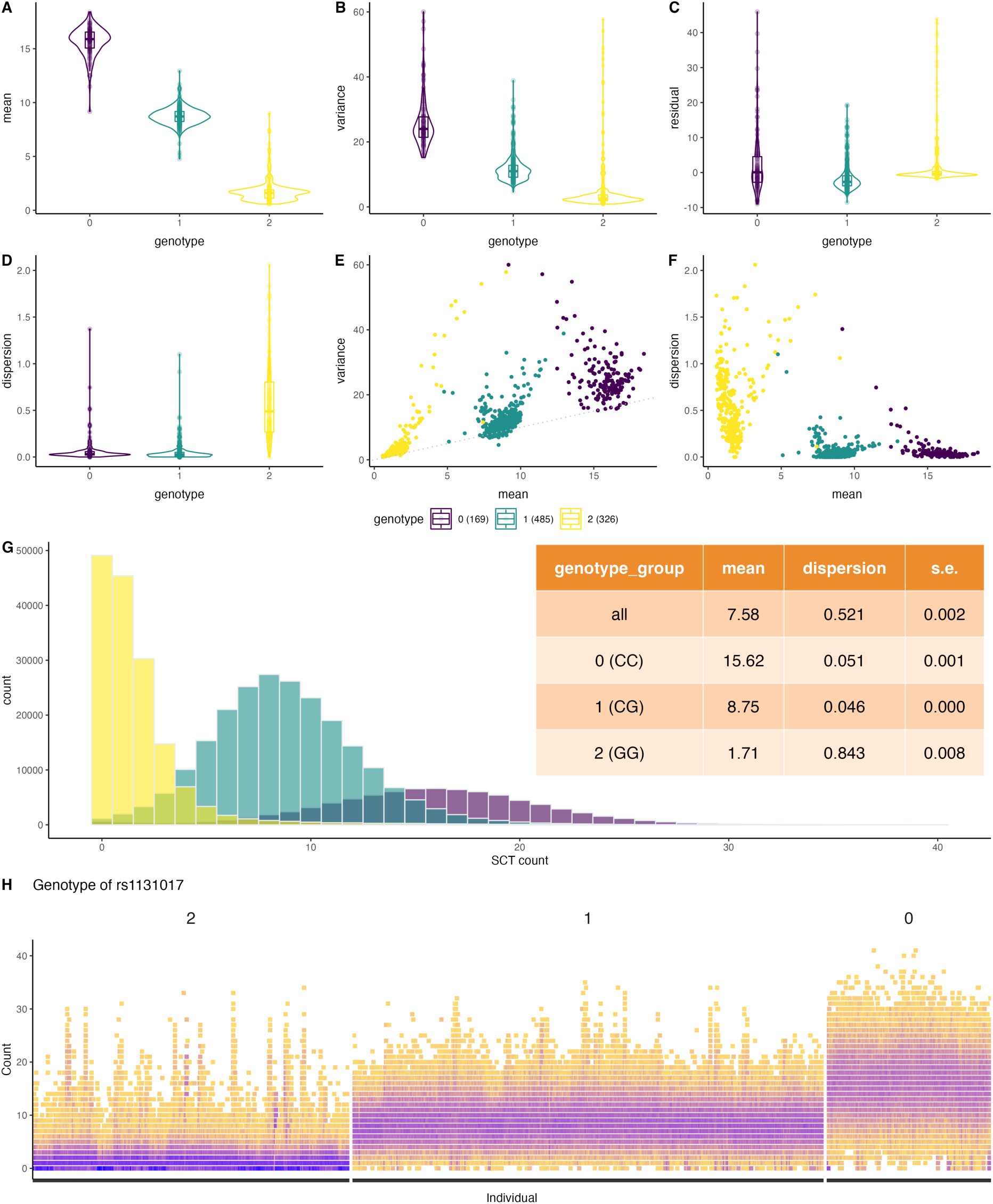
The association between rs1131017 and *RPS26* expression change in CD4 naïve cells. **(A-D)** The violin plots of individual genotypes of SNP rs1131017 correspond to the intra-individual mean, variance, residual, and dispersion of *RPS26* expression in CD4 naïve cells. The x-axis indicates the genotype (coded as 0, 1, 2 indicating the number of G alleles carried). **(E)** Scatter plot of intra-individual mean against intra-individual variance of expression. **(F)** Scatter plot of intra-individual mean against intra-individual dispersion of expression. The dispersion was estimated by CR-MLE method. **(G)** The distribution of SCT transformed count expression of *RPS26* per cell. There are 463,496 cells, and each bar indicates the number of cells with the corresponding count expression. The table included the mean and dispersion estimates for the whole cohort and within each genotype group. The genotype group of CC alleles have much higher intra-individual dispersion of *RPS26* expression than the other two groups. (**H**) The distribution of SCT transformed count expression for all 980 individuals in CD4_NC cells separated by three genotype groups. The colour of each square denotes the density of a certain count within a corresponding individual, and darker purple denotes higher density.

**Table 2.**
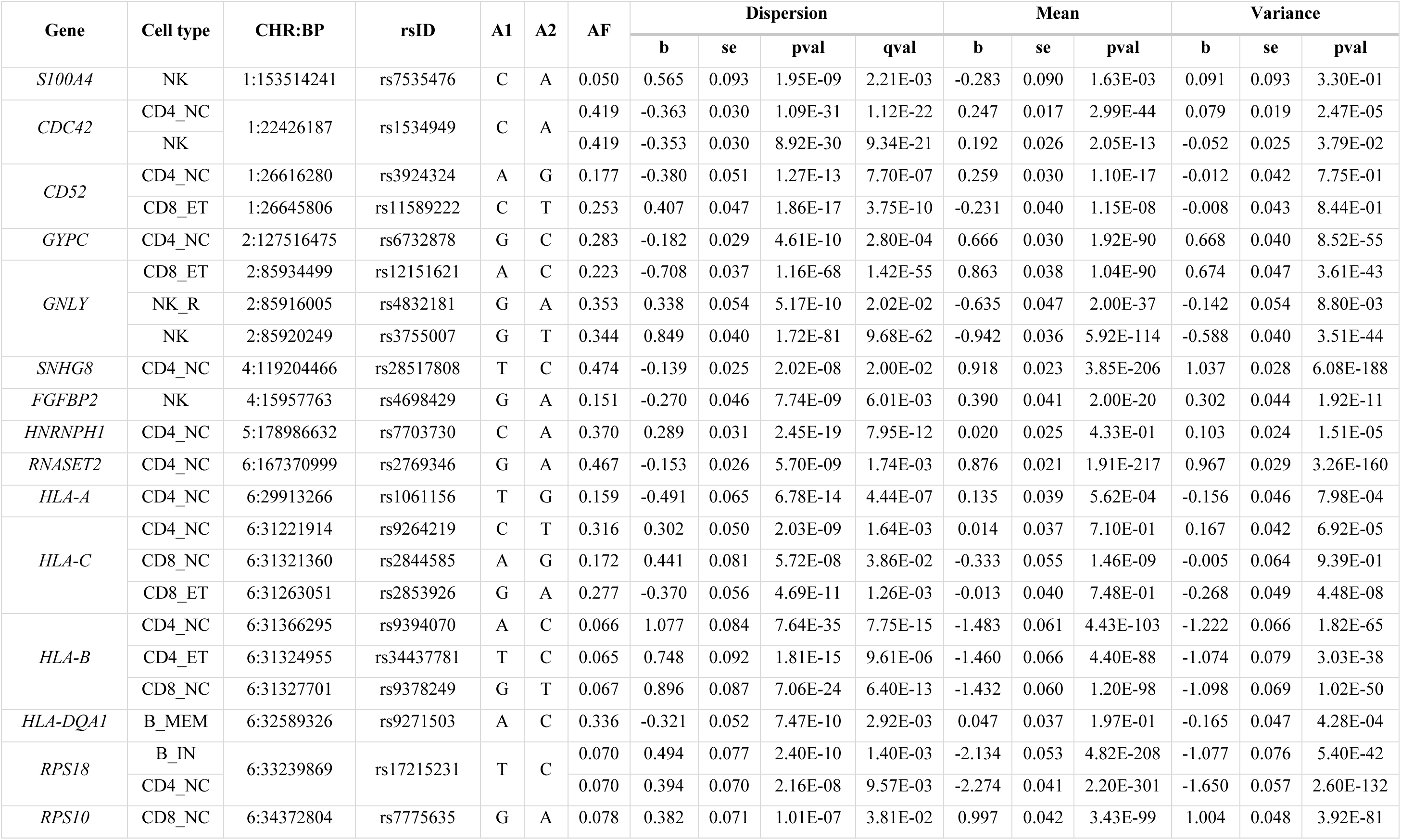
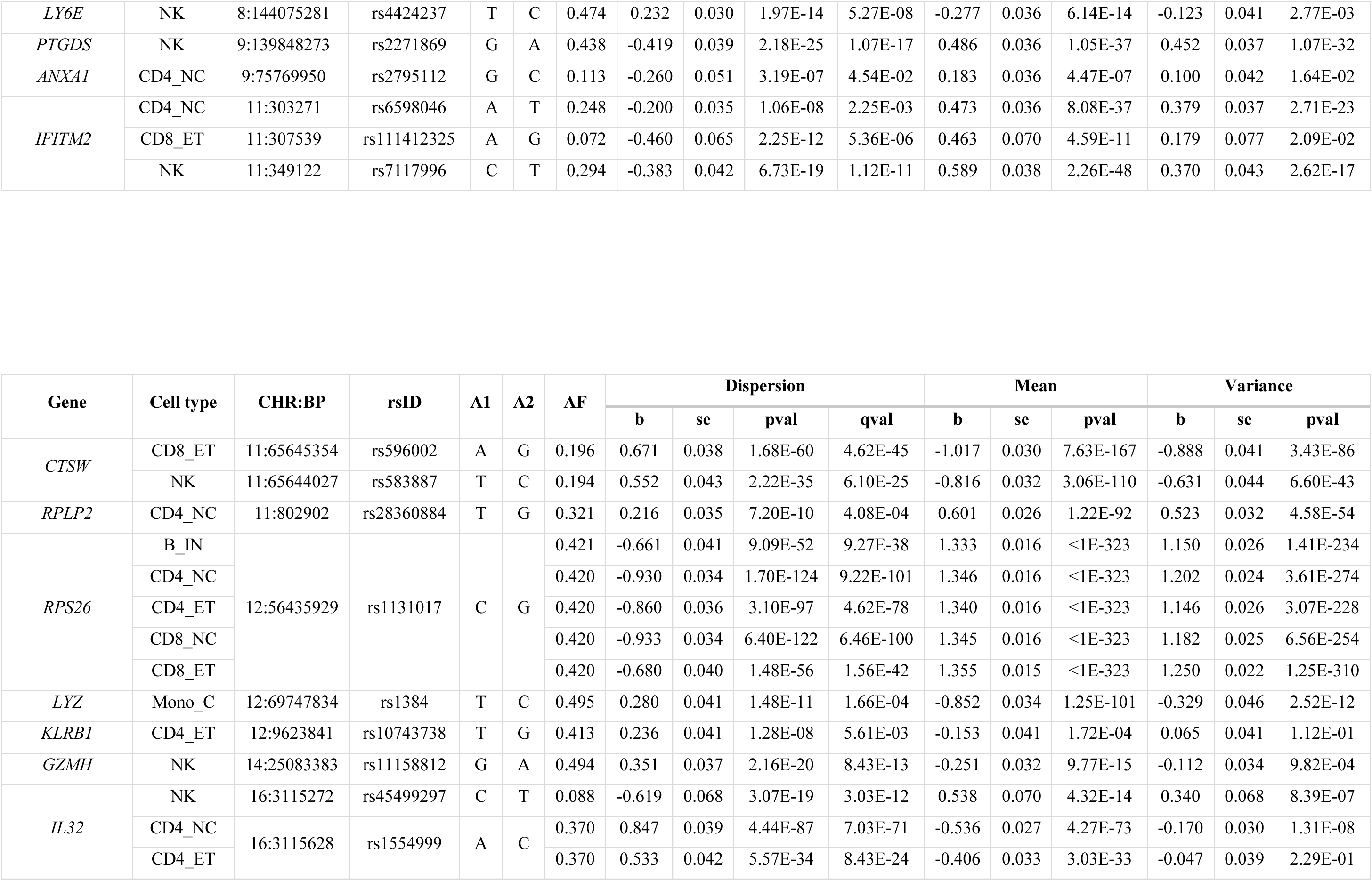
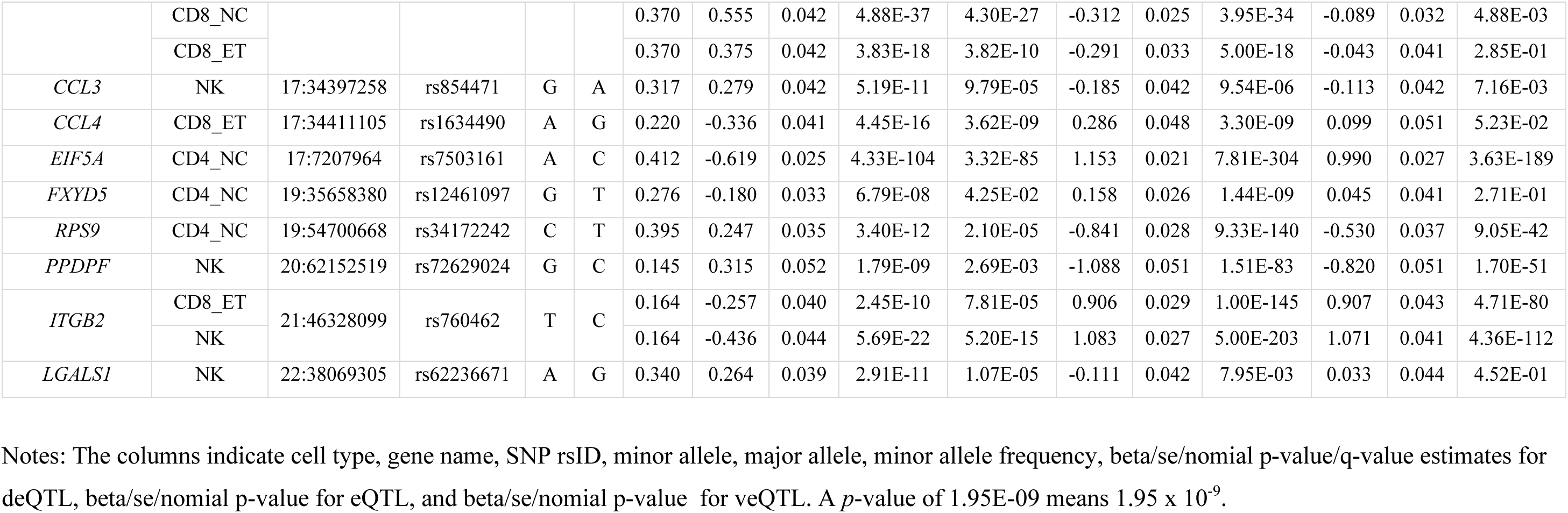
Summary of the 39 dGenes identified in 14 cell types.

We subsequently evaluated the degree of cell-type specificity of the 34 dGenes. Of these, 67.6% (23) have evidence of an allelic effect in just one cell type. Of the remaining 11 dGenes, there are five genes with sc-deQTL in two cell types, four genes with sc-deQTL in three cell types, and two genes with sc-deQTL in five cell types respectively (**Figure 3** and **Table 2**). Notably, two genes (*IL32* and *RPS26*) had significant deQTLs in five cell types (**Table 2**). For example, in both naïve and effective CD4 and CD8 cells, we identified rs1554999 as the top deQTL for *IL32*. The A allele shows an increasing effect on the dispersion level of this gene. On average, individuals carrying each copy of the A allele of rs1554999 have an increase of 0.25 in the dispersion level of mRNA transcript molecules across cells. The top deQTL in the NK cell is a different SNP, rs45499297, but in LD with rs1554999 (*R*^2^ = 0.056, D’ = 1). The rs1554999 is located in the 5’ UTR region of *IL32*, and was previously reported to be strongly associated with methylation level at three CpG sites in CD4+ T cells^33^. While most dGenes were also vGenes, there was minimal overlap with residual genes (**Supplementary Table 3**), providing further supporting evidence that the intra-individual residuals capture different sources of variation from the intra-individual dispersion.

### Functional annotations of deQTLs and dGenes

To understand the functional characteristics of deQTLs and dGenes, we first used ANNOVAR^34^ to perform variant annotation for the top deQTLs and subsequently tested for dGenes overlap with GWAS loci (**Methods**). Most deQTLs are in the intergenic or intronic regions, implying they affect intra-individual variation via regulatory mechanisms. This observation aligns with those made for single-cell mean-effect eQTLs. The majority of dGenes (31 out of 34) are reported to be associated with either inflammation, immune responses, or viral infections. An example is *GNLY*, which has significant sc-deQTL in NK, NK_R_, and CD8_ET_ cells (**Table 2**). *GNLY* encodes a protein called granulysin, present in natural killer cells and cytotoxic T lymphocytes, and has been shown to have antimicrobial activity against tuberculosis through changes in cell membrane integrity and reducing the viability of the bacillus^35,36^. All of the sc-deQTLs for *GNLY* are located in intergenic regions. For example, the sc-deQTL for CD8_ET_ (rs12151621[A/C]) is located in a *CTCF* binding site (ENSR00000291834), ∼8.1kb downstream of *GNLY*. The A allele of this SNP has a decreasing effect on the dispersion of *GNLY* expression but an increasing effect on the mean expression, which means individuals with the CA or CC genotype will have lower mean expression but higher dispersion than the individuals with the AA genotype (**Supplementary Figure 9**). We hypothesise that the C allele leads to a higher binding affinity of CTCF, and the binding will repress the expression of *GNLY*. The A allele has been reported to have an increasing effect on the protein levels^37^ and toxoplasma antibody IgG levels^38^ (*p*-value = 9.1 x 10^-4^, not genome-wide significant given limited sample size = 557). This may suggest that individuals with an A allele tend to have higher levels of IgG antibodies against the toxoplasma parasite. The body will produce more *GNLY* proteins in the NK cells but less variability across cells, ensuring enough functional proteins to kill the microbes. A second example is seen in *GZMH,* which encodes a serine protease (granzyme H) and is constitutively expressed in CD8_ET_ and NK cells. We identified a deQTL (rs11158812[G/A]) which only has a significant effect in NK cells (*p*-value = 2.16 x 10^-20^, *q*-value = 8.43 x 10^-13^). The top deQTL is located at the intergenic region between *GZMH* and *GZMB* (no association between rs11158812 and *GZMB* in any cell type), a region that has only been reported to be associated with vitiligo^39^. The top GWAS hit (rs8192917) is a missense variant (Arg55Gln) of *GZMB* but not in LD with rs11158812 (*R*^2^ = 1.2 x 10^-3^). Another example is *IFITM2, which* encodes interferon-induced transmembrane protein 2, an interferon-induced antiviral protein family member. This locus is not associated with human diseases but is highly associated with human blood cell traits^40^ such as granulocyte, neutrophil, and eosinophil counts. The top deQTL of *IFTIM2* differs in three cell types, but for naïve CD4 (CD4_NC_) and CD8_ET_ cells, the top deQTL is the same as the top eQTL and in linkage disequilibrium (*R*^2^ = 0.11 and *D*’ = 0.69). For NK cells, the top deQTL (rs7117996) is located in the intergenic region and independent from the other two top deQTLs (*R*^2^ < 0.01), but its top eQTL (rs1059091, independent from rs7117996) is a missense variant for *IFITM2*.

There are also dGenes whose top deQTL is located outside the intergenic region. For example, *ITGB2* (integrin subunit beta 2) is a significant dGene in CD8_ET_ (*q*-value = 7.81 x 10^-5^) and NK cells (*q*-value = 5.20 x 10^-15^). In a previous study, the top deQTL (rs760462[A1/A2]) located in intron 3 of *ITGB2* was annotated as a splice acceptor variant^41^. *LYZ* encodes lysozyme, which has antibacterial activity against several bacterial species and is highly expressed in monocytes (Mono) and dendritic cells (DC). The top deQTL rs1384 is a 3’ UTR variant, and a recent study^42^ has suggested a monocyte-specific *trans-* action mediated by *LYZ* in this site. Fairfax et al^43^ have reported *LYZ* as a monocyte-specific master regulator and its monocyte-specific *cis*-eQTL (rs10784774, in complete LD with rs1384) is also a *trans*-eQTL to 62 genes. The G allele of rs10784774, as well as the T allele of rs1384, is associated with lower expression but higher dispersion level of *LYZ* in classical monocytes (**Table 2**), and also reported to be the increasing allele for neutrophil percentage of white cells^40^.

Using the FUMA platform, we performed pathway enrichment analyses on the dGenes (**Methods**), and identified significant enrichment in three Hallmark gene sets: “Allograft Rejection”, “Interferon Gamma Response”, and “Interferon Alpha Response”. The KEGG pathway enrichment identified 12 significant pathways, with the top three “KEGG Viral Myocarditis”, “KEGG Ribosome”, and “KEGG Allograft Rejection”. However, after removing the five MHC genes, only “Allograft Rejection” remained significant in Hallmark gene sets and “KEGG Ribosome” in the KEGG pathway (**Supplementary Table 4**). We also performed an enrichment analysis of dGenes for GO biological process. The most enriched process is “Interspecies Interaction Between Organisms” and “Cytokine Mediated Signaling Pathway”, and “Viral Gene Expression”. Our results suggest that the dispersion effects on the gene expression across cells within individuals are enriched in the biological process of immune response and viral infection. This is consistent with the prior knowledge that the G x E effect could induce phenotypic variability, and thus the potential environment is worth further investigation.

### Trans-regulation of dGenes partially explained the dispersion difference

Next, we sought to explore potential genomic factors underlying deQTLs. Previous vQTL GWAS, testing for genetic effects on high-order trait variability between individuals, have often explored whether variance effects are caused by G x G or G x E effects. Intuitively, we hypothesise if the dispersion difference in gene expression could be explained by (1) G x G *cis*-epistasis, where multiple independent *cis*-SNPs affect the expression variability of the target gene; (2) G x G *trans*-epistasis, where *trans*-QTLs are regarded as an interaction effect for the target gene and influence the variability level via the trans-regulation.

We compared the number of independent cis-QTLs for the 34 candidate dGenes to test if multiple cis-regulations could drive the dispersion difference. On average, each dGene has 2.36 independent *cis*-eQTLs and 1.71 independent *cis*-veQTLs. The comparison also showed that no dGene has more independent *cis*-veQTLs than *cis*-eQTLs except for *HNRNPH1* (but it is only 0 vs 1, so it is not a large difference). However, 14 dGenes have significant *trans*-deQTLs, among which there are also five dGenes (*RPS18*, *SNHG7*, *GNLY*, *CCL3*, and *LGALS1*) that do not have any trans-eQTL or veQTL (**Supplementary Table 5**). For example, rs78089025 [A1/A2] (9:73039725) showed a genome-wide significant (*p*-value = 4.04 x 10^-8^) association with the dispersion levels of *GNLY* in NK cells but not with the mean or variance levels. This SNP is an intron variant for *KLF9-DT*, a divergent transcript of transcription factor *KLF9*. When fitting *GNLY*’s the *trans*-deQTL (rs78089025) and top *cis*-deQTL (rs3755007) in the same association model, the interaction term showed significant effects on the dispersion level (*P*_interaction_ = 4.93 x 10^-12^), and a significant change in the main effect. Specifically, when only top *cis*-deQTL was fitted, the beta = –0.171, *s.e.* = 0.009, and when *trans*-deQTL was fitted in the interaction model, the beta = –0.484, *s.e.* = 0.046. These results imply that the deQTLs are not induced by the G X G effect from independent *cis*-SNPs but can be partially explained by the *trans*-SNP effects on the dGenes.

### Genetic control of variance heterogeneity underlying different contexts

Since we show that *trans*-regulation could be a putative driving factor for deQTLs, we further asked if the G x E interaction between genetic effects and cellular state also affects intra-individual dispersion. To test this, we inferred the cell state landscape for B cells and fitted the average cell state per individual as an interaction term in the deQTL association model (**Methods**). Only one significant interaction between genotype and cell state was identified for dGene *RPS18* in B_MEM_ cells (adjusted *p*-value = 4.66 x 10^-3^). One plausible explanation is that the pseudotime was calculated based on highly variable genes, so the gene expression variability of the dGenes might not be well captured by this approach.

We also tested if deQTL effects are associated with the interaction between genotype and sex or age. For genotype by age interaction, only *HLA-B* in naïve CD8 (CD8_NC_) cells and *RP11-1143G9.4* in classical monocyte (Mono_C_) cells showed significant associations (**Supplementary Table 6**). For genotype by sex interaction, only *HLA-B* showed significant interaction in effective CD4 (CD4_ET_) (rs34437781) and CD8_NC_ cells (rs9394070). The rs34437781 was a significant deQTL in CD4_ET_ cells (nominal *p*-value = 1.81 x 10^-15^, *q*-value = 9.61 x 10^-6^). rs9394070, which is in strong linkage disequilibrium (LD) with rs34437781 (R^2^

= 0.886, D’ = 0.949), is a significant deQTL for *HLA-B* in CD8 naïve cells and top deQTL in CD4 naïve cells. Still, when fitting the genotype by sex interaction term in the model, the genotype itself became insignificant (genotype *p*-value = 5.74 x 10^-3^, interaction *p*-value = 5.43 x 10^-8^). The interaction is mainly induced by only females with the TT genotype (4/980), where their intra-individual mean estimates are the lowest among 980 individuals but the dispersion estimates are relatively higher (**Supplementary Figure 9**). After removing the four individuals with the TT genotype, the interaction term was no longer significant (interaction p-value = 0.029). Similarly, the beta of the deQTL when performing association test in separate sex group are not significantly different (*p*-value = 0.165). These results indicate that neither sex nor age is the main driving factor of the genetic effects on the dispersion level across cells.

### Association between dispersion eQTL and immune phenotypes

To understand the relationship between sc-deQTL and disease risk, we tested for the overlap between sc-deQTL loci and public GWAS associations in GWAScatalog (**Methods**). The most frequent traits include blood protein levels, asthma, eosinophil counts, type 1 diabetes, Crohn’s disease, height, and rheumatoid arthritis. Combined with the FUMA enrichment results above, it further suggests that the intra-individual dispersion effects are enriched in the genetic association with auto-immune and infectious diseases.

Highlighting *RPS26* as an example, carrying copies of the G allele for rs1131017[G/C] has an increasing effect on the intra-individual dispersion (**Figure 4**). This SNP was tested against our association analysis’s dispersion level of 53 *cis*-genes but was only significant for *RPS26*, with shared allelic effects across five cell types (innate B cell, naïve/effective CD4, and naïve/effective CD8). This locus has previously been reported to be strongly associated with auto-immune diseases, including type 1 diabetes^44^, asthma^45^, vitiligo^46^ and rheumatoid arthritis^47^. The top SNPs are not the same one but all are in strong LD with each other (*R*^2^ > 0.8). Furthermore, the C allele of rs1131017 was consistently shown to have an increasing effect on these auto-immune disease risks, which suggests that the lower dispersion of intra-individual expression of *RPS26* is associated with higher auto-immune disease risk. When we directly estimate the dispersion of the unique molecular identifier (UMI) count distribution (after SCTranformation) across all cells in each genotype group (using CD4_NC_ cells as the example), the CC genotype group shows a much larger over-dispersion (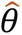 = 0.843) than the CG or GG group (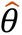 = 0.046 and 0.051) (**Figure 4G**). So, we further dissect the count distribution for each individual. We observe that the UMI count distribution for individuals in the CC genotype group mostly follows a negative binomial distribution, while the UMI count distributions for CG and GG individuals follow a Poisson distribution. This suggests that even for the same gene, genetic effects could impact the distribution type across the cells within an individual.

The top SNP for *RPS26*’s deQTL is located in the gene’s 5’UTR region, the binding site for six transcription factors (*RBM39*, *TCF7*, *LEF1*, *KLF6*, *CD74* and *MAF*)^48^. We hypothesized that if these transcription factors (TFs) regulate the expression level of *RPS26* via binding to this site, rs1131017 should be detected as a co-expression eQTL between *RPS26* and these TFs in our data. To evaluate this, we calculated the co-expression between each TF and *RPS26* within each cell type and ran a co-eQTL association analysis (**Methods**). For all 84 (6 x 14 cell types) tests, we identified 40 significant co-eQTLs (FDR < 0.05) (**Figure 5**). The most frequent (11/14) co-deQTL is between *RPS26* and *CD74*. Interestingly, the allelic effect in T cells is in the opposite direction to those in B cells and monocytes (**Figure 5A**). These results indicate that the potential regulation of *CD74* on *RPS26* via promoter binding is cell-type specific. In CD4_NC_ cells, all six transcription factors have significant co-eQTL estimates, and *TCF7* and *LEF1* showed the opposite direction of the co-expression to the other four TFs (**Figure 5B**). Interestingly, the effect size of the co-eQTL is generally larger in the naïve CD4 and CD8 cells compared to the effective cells, and it is not driven by the difference number of cells between naïve and effective cell types. For CD4 cells, all six transcription factor genes showed significantly (*p*-adjust < 0.05) larger co-deQTL effect size in the naïve cell type. This relationship is also observed for *KLF6*, *CD74*, and *MAF* in CD8 T cells. These results indicate that the effect of rs1131017 on the co-expression level between *RPS26* and the transcription factors is impacted by an immune response.

**Figure 5.**
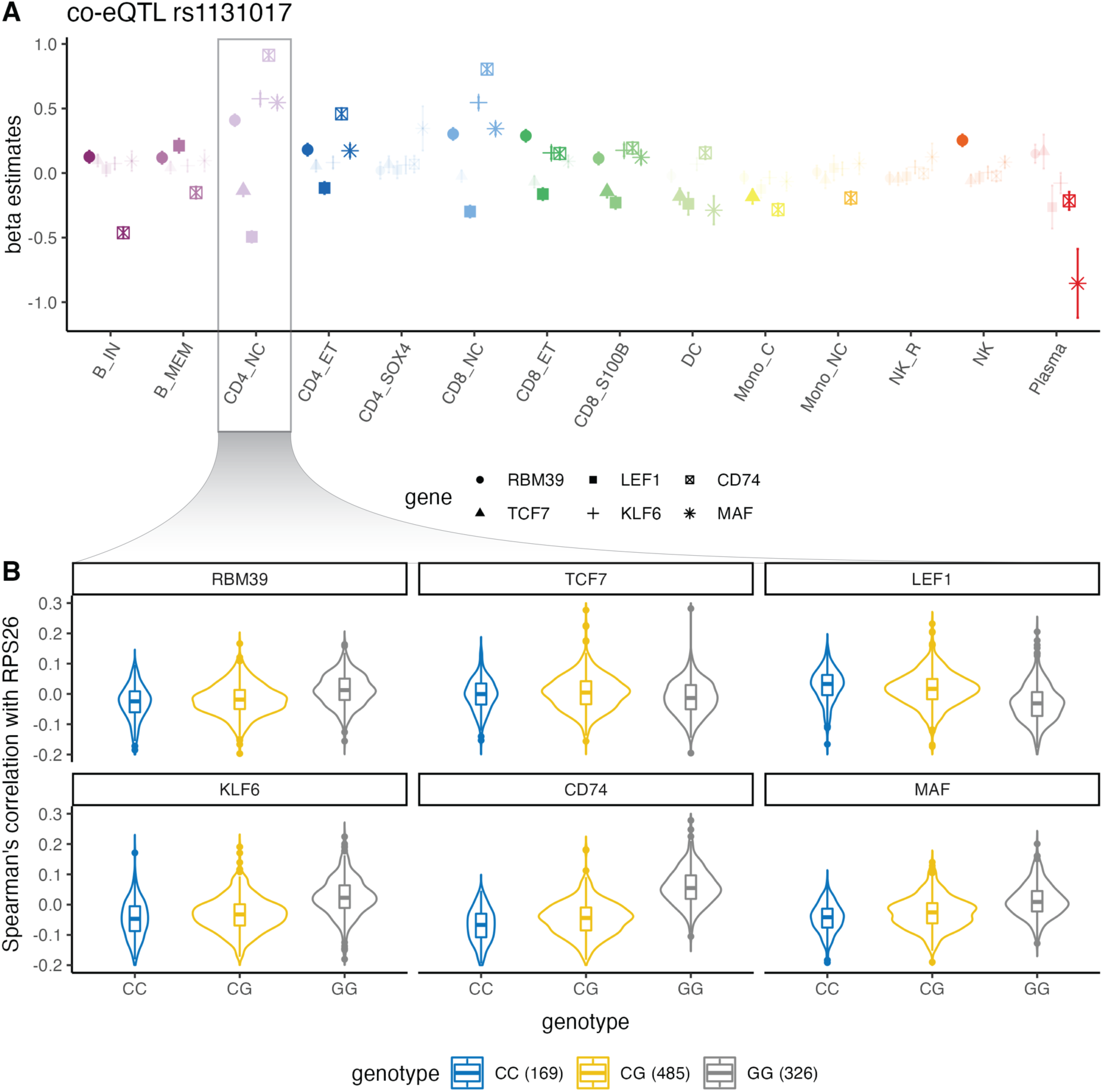
The putative regulatory mechanism of dispersion eQTL of *RPS26*. (**A**) The beta effect estimates for co-eQTL between *RPS26* and six transcription factors (TFs) in 14 cell types. Each point is an estimate coloured and grouped based on cell types, and the error bar denotes the standard error of the estimate. Six TFs are annotated with different shapes. The significant estimates (BH corrected FDR < 0.05) are strengthened by transparency of the dot. (**B**) The violin plot for co-expression per genotype group in CD4_NC_ cells. The y-axis indicates the standardised the co-expression (measured by Spearman’s correlation across cells per individual) between *RPS26* and the TF gene. Four outlier dots were omitted for illustrative purpose.

## Replication of sc-deQTL in independent cohort

We attempted to replicate our identified sc-deQTL in an independent cohort from Perez et al.^49^ Out of 34 dGenes we identified in OneK1K, we replicated 6 genes (*LYZ*, *HLA-DQA1*, *RPS26*, *CD52*, *GNLY*, and *CTSW*) with significant deQTL (FDR < 0.05). Given the different SNP panels and cell type annotations between the two cohorts, only two genes (rs596002-*CTSW* in CD8 with FDR = 0.026; and *LYZ*-rs1384 in monocyte with FDR = 3.23E-10) have the exactly the same deQTL in the same cell type. For the most significant example of rs1131017-*RPS26*, we only detected significant deQTL association in classical monocyte (*p*_nominal_ = 4.97E-07, FDR = 07.62E-3). The association did not pass the multiple correction (*p*_nominal_ = 2.43E-06, FDR = 0.436), but the association patterns were very consistent between OneK1K and Perez et al (**Supplementary Figure 10**) and the correlation of the test statistics –log10(*p*_nominal_) between two datasets is very high (Pearson’s cor = 0.894). Thus, we speculated that the insignificant replication is mainly due to the limited power given the tiny sample size of the replication cohort.

## Discussion

In this study, we present MEOTIVE, a robust framework to identify genetic variations associated with intra-individual variability in gene expression single-cell level. MEOTIVE addressed the issue associated with the mean-variance relationship, exacerbated by the non-gaussian distributions of scRNA-seq data. By applying MEOTIVE to data from the OneK1K cohort, it is the first study to identify intra-individual deQTLs at the population scale successfully. While most previous studies focused on the mean effects of the genetic variants on gene expression, the genetic effects on the variability and dispersion across cells within individuals are poorly understood and single-cell RNA-seq data provides a solution for dissecting the high-dimensional effect on genome regulation. In total, we identified 34 dGenes accounting for the mean-variance dependency and they were enriched in the biological pathways relevant to interferon response, interspecies interaction, allograft rejection and viral gene expression. These results suggest that the transcriptional variability at the single-cell level could arise due to immune and/or external stimulus^50^ and that variability is under genetic control. Although we only identified 55 deQTL-dGene pairs in the current study, given a larger sample size and number of cells per individual, we would expect to discover thousands of genetic variants that affect the dispersion level of intra-individual gene expression. New analyses with larger datasets are set to uncover a fundamentally new avenue of genetic effects on human genome regulation.

Herein, we also propose a new explanation for the “zero-inflated” model in single-cell RNA-seq data. In the example of rs1131017-*RPS26*, it was noted that an observed zero-inflated negative binomial (ZINB) distribution across a population of cells results from a mixture of three genotype groups (CC, CG, GG) each showing a negative binomial distribution (**Figure 4G**). In such cases, the so-called “structural” or “excessive” zeros are not generated from a separate biological process but from the genotype group with low abundance expression. This finding challenges the conventional understanding of the underlying model of scRNA-seq count data and necessitates the need to re-evaluate the previous zero-inflated negative binomial model.

This study has several limitations. First, although it is the largest single-cell eQTL cohort, with data from 980 individuals, we are limited in statistical power to testing only SNPs with MAF > 5%. Sarkar et al.^23^ predicted that it needs 4,000 individuals to achieve 80% to detect the deQTLs. The number of cells per individual per cell type is also an important limitation because several cell types only have 100 to 200 effective samples. Second, the accuracy of dispersion estimation is mainly affected by the mean and number of cells per individual. From our simulations, we observe that given 500 cells, we need a mean > 0.3 to have an accurate estimate for dispersion. Should we have 5,000 cells per individual for a certain cell type, the filtering threshold for the mean expression can go down to 0.1, which can rescue more genes for association testing (from 3% to 10%). Assuming the largest group is CD4_NC_ cells, the minimum requirement for the number of cells of single-cell RNA-seq data would be ∼10,000 per individual. Even in such cases, the rare cell types such as plasma or dendritic cells would still only have 70 to 100 cells per individual. Third, the high sparsity in the 10X data is one of the reasons preventing us from better understanding the underlying model of scRNA-seq data. A recent study^51^ demonstrated that lower sequencing depth would make the observed data more similar to Poisson distribution even if the true model is over-dispersed. Since ∼50% intra-individual mean is 0 for our data obtained using 10X v2 kit, processing samples with a higher capture rate will benefit the estimation of the true underlying distribution of scRNA-seq data.

In summary, we identified genetic effects on the within-individual gene expression variability while accounting for the mean-variance dependence in the scRNA-seq data of human PBMCs. The MEOTIVE statistical framework we present here can be implemented on any single-cell RNA-seq dataset with genotype information to identify genetic variations that influence intra-individual variability of gene expression. As cohort sample sizes increase (e.g., TenK10K), ongoing analyses will continue to reveal novel genetic mechanisms underlying inter-individual variability and cell-to-cell heterogeneity.

## Online Methods

### The OneK1K cohort

The OneK1K cohort is a collection of genotype and single-cell gene expression data for 982 individuals of Northern European ancestry. Each individual was genotyped and imputed with 759,993 SNPs against the HRC panel^52^. There are 1,267,758 peripheral blood mononuclear cells (PBMCs) with gene expression data after demultiplexing and doublets removal. Identical to the cell type classification in Yazar *et al*.^6^, we predicted the OneK1K cohort into 14 cell types based on the *scPred* method^53^ (**Table 1**). During a sensitivity test of latent variables^54^, we identified two outlier samples (one due to a low number of cells and the other due to extremely imbalanced cell composition). We excluded them from all the analyses in this study. Thus, the final sample size we retained in this study is 980.

### Strategy of estimating genetic effects on intra-individual expression variability

To accurately estimate the mean and variance of intra-individual gene expression, we applied several steps to exclude potential confounding factors in the single-cell RNA-seq data. First, the count matrix was pre-processed by Seurat using *sctransform* algorithm^55^ to remove the technical confounders such as sequencing depth and batch effects. Second, all cells were classified into 14 different cell types by a semi-supervised method (scPred^53^), and individuals with less than five cells in each cell type were excluded to avoid biased estimation for intra-individual mean and variance driven by outliers. Third, genes expressed in less than 10% of individuals or the intra-individual mean across the cohort less than 0.001 were also excluded (**Methods**). After the quality control, the median number of cells per individual ranges from 7 (plasma cells) to 461 (CD4_NC_ cells), and the number of individuals and genes also varies across different cell types (**Table 1**).

We first used the moment estimate of the intra-individual mean and variance per gene and generated two *M* * *N* matrices for each cell type (**Methods**), where *M* is the number of genes and *N* is the number of individuals for each cell type.

The moment estimates of mean and variance for gene *k* of individual *i* across cell *j*, are denoted as the following:

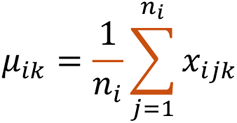

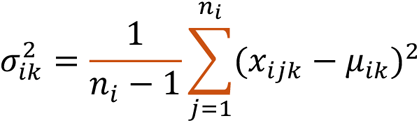

where *n_i_* indicates the number of cells for individual *i*. We also calculated the proportion of zero expression within an individual (π_0_) and detected a strong negative relationship between the π_0_and μ (**Supplementary Figure 11**).

For single-cell RNA-seq data, the intra-individual mean and variance are correlated since a large proportion of them follow non-normal distributions such as Poisson, NB or ZINB distributions^23,51^. In our OneK1K data set, Spearman’s correlation coefficients between intra-individual mean and variance across individuals per gene were extremely high. For example, ∼94.6% of genes showed 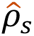 > 0.8 and ∼75.2% showed 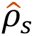 > 0.99 (**Figure 2A** and **Supplementary Note 1**). On the other hand, the correlation estimates were strongly dependent on the mean expression level in a negative trend (**Figure 2B** and **Supplementary Figure 2**), and so did the proportion of non-expression individuals per gene (**Figure 2C** and **Supplementary Figure 11**). Given such a strong mean-variance dependency, we predict that the significant veQTLs could be primarily explained by the effects of the mean difference. To better understand the characteristics of vGenes and its relationship with eGenes, we compared the vGenes with or without the ME on several aspects. First, the vGenes without ME showed significantly larger *q*-values than those with ME in all cell types, but most of the *q*-values were just clustered around 0.05 (**Supplementary Figure 3**). Second, the effect sizes of the veQTLs without ME were much smaller than those with ME in five cell types (CD4_NC_, CD4_ET_, CD8_NC_, CD8_ET_, and NK), with 1.54∼3.94 fold smaller for the median effect size (**Supplementary Figure 4**). Third, we tested if the veQTLs without ME were closer to the TSS location than those with ME. The results showed that the TSS distance of eQTLs is not significantly different from veQTLs (**Supplementary Figure 5**, Welch Two Sample t-test, *p*-value = 0.91) and veQTLs without ME are uniformly distributed based on the TSS distance. Thus, alternative ways to rule out the mean effects are needed when estimating the genetic effects on the variability of intra-individual gene expression. We also estimated the relationship between the mean and other metrics, including the variance-mean-ratio (VMR) and coefficient of variation (CV). These two metrics are also highly correlated with the intra-individual mean (**Supplementary** Figures 6-7). Previous study has used them as the dispersion indicator in the eQTL data set^23^. However, since most of the mean of gene expression in single-cell RNA-seq data is very close to 0, the VMR or CV could approach large numbers and are sensitive to even tiny changes when the mean expression is low. Thus, VMR and CV are not suitable for the dispersion indicator in the single-cell data sets.

### The TensorQTL for SNP-gene association analysis

To understand how genetic variations between individuals affect the variance of intra-individual gene expression, we performed association analysis using TensorQTL^28^ for each cell type. We first filtered out those genes with expression in less than 10% of individuals or extremely low inter-individual abundance (μ < 0.001). The intra-individual mean, variance, and dispersion were log(x+1) transformed and then z-score normalized per gene to avoid extreme outliers. The residual expression matrix was just z-score normalized per gene. The sex, age, first 6 principal components (PCs), and first 10 PEER factors^56^ were fitted as covariates in the model. The PCs are calculated by PLINK^57^ based on the genotype information. The PEER factors are derived based on the intra-individual mean of gene expression matrix for each cell type to capture the latent variables. We chose 10 PEER factors to be fitted in the association model by a sensitivity analysis and a local greedy method to balance the discovery power and overfitting^54^. The number of remaining individuals, genes, and median number of cells per individual for each cell type are presented in **Table 1**. In the *cis-*QTL analysis, we only retained ∼4.2 million SNPs located within ± 1Mb *cis*-region from the centre of the gene body and with a minor allele frequency (MAF) larger than 0.05. After obtaining the nominal *p*-values for every SNP-gene pair, a beta-approximation permutation was applied to correct the *p*-values and 10,000 times of permutations were conducted for each gene. The most significant SNP for each gene (top *cis*-eQTL or veQTL) was further corrected and the permuted *p*-value was converted to a *q*-value to control the false positive per chromosome^58^. An SNP-gene association with *q*-value < 0.05 was deemed significant.

### The estimation of dispersion in gene expression distribution

We adopted two methods to generate the estimates for dispersion for the intra-individual gene expression.

First, we used a straightforward method and regressed out the intra-mean from the intra-variance and used the residuals as the dispersion indicator;

Second, we assume all the intra-individual gene expression follows a negative binomial (NB) distribution. Let:

- *x_ijk_* be the number of molecules for individual *i*, cell *j*, gene *k* after accounting for confounders and size factor
- μ*_ik_* be the mean of expression of gene *k* in individual *i*
- θ*_ik_* be the dispersion of expression of gene *k* in individual *i*

Then we assume,

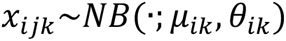

The likelihood function for the intra-individual distribution of each gene is

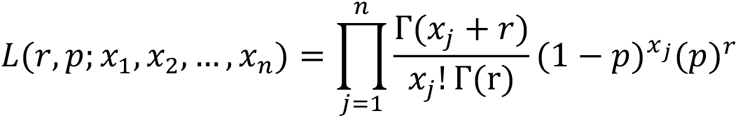

We need to estimate the *r* and *p*, where 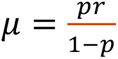 and 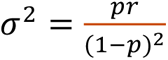. Alternative parameterization is to use theta (θ = 1/r) as the dispersion parameter. We used the “*glmGamPoi*” R package which implements the Cox-Reid adjusted MLE^59^ method to estimate the dispersion parameter based on the SCTranformed count data. Then we generated an intra-individual dispersion matrix for each cell type.

When checking the preliminary results (326 deQTLs with *q*-value < 0.05), however, we found that for many genes, the CR-MLE estimates were highly inflated, especially for those with low abundance or individuals with a small number of cells (**Supplementary Figure 12**). This is because when the mean of an NB distribution is low, the likelihood curve will be very flat, thus making it extremely difficult for the optimisation algorithm to search for the maxima. This scenario became even worse in the single-cell RNA-seq data since less than 10% of the genes have an intra-individual mean higher than 0.1, and around half of the intra-individual expression is 0 (**Supplementary Figure 2**). Although the CR-MLE method partially mitigates the problem using a penalised log-likelihood^60^, we still saw inflated dispersion estimates in many genes from the real data (**Supplementary Figure 12**). To avoid false discovery of deQTLs and the spurious relationship between dispersion and mean, we simulated CR-MLE dispersion estimates given different mean, dispersion, and sample sizes (**Supplementary Note 2** and **Supplementary** Figures 13-14). We considered these parameters and adopted a data-driven threshold of intra-individual mean expression to select significant signals for each cell type (**Supplementary Table 7**). For example, for CD4_NC_ cells, we need genes to have mean expression > 0.3 so that > 90% times the estimates will fall within ±5% of the true dispersion parameter. Based on this filtering, we retained 64 deQTL-dGene pairs but still found that there is still moderate inflation in the dispersion estimates of some genes in certain genotype groups (see examples in **Supplementary Figure 15**). So, we further removed the genes if any of the genotype groups has a mean expression smaller than the threshold and ended up with 55 significant (*q*-value < 0.05) deQTL-dGene pairs in 34 unique genes.

We also tried to estimate the dispersion for gene *k* of individual *i* based on the methods of moments, such that

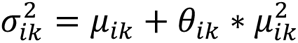

so, the dispersion can be estimated as,

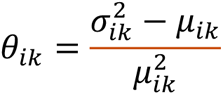

From the equation, it is obvious that (i) when the variance and mean are very close to each other and the mean is not so small, the moment estimator will be close to 0; (ii) when the mean is 0, the moment estimator does not exist but for such case, we manually assign the dispersion level as 0; (iii) in real data, the moment estimator could be a negative number, but the scale would not be large (CR-MLE estimate will always be non-negative). More details of different dispersion indices and their special cases, when the mean is small, are discussed in **Supplementary Note 3** and **Supplementary Table 8**.

### Functional annotation and gene sets enrichment analysis

We used FUMA^61^ (v1.5.3) to perform functional annotation for the deQTLs and gene enrichment for the dGenes. The SNP functional annotation is done by the built-in ANNOVAR^34^ in the platform. To test if the list of dGenes is overrepresented in certain biological functions, they are tested against the gene sets obtained from The Molecular Signatures Database (MSigDB). The Hallmark gene sets are from the MSigDB h collection (50 gene sets), the KEGG are from MSigDB c2 collection (186 gene sets) and GO biological processes from MSigDB c5 collection (7751 gene sets). The significant enrichment is defined at adjusted *p*-value ≤ 0.05. We also utilised FUMA platform to overlap the sc-deQTL with public GWAS associations in GWAScatalog (database update by 27/4/2023). There are 553 matched associations in 233 studies.

### The G x G epistasis analysis to identify driving factors for deQTL

We performed two complementary analyses to investigate whether G x G and G x E effects could be the potential driving factors for deQTLs. To identify multiple independent cis-signals for the same gene, we sought to map conditionally independent *cis*-eQTLs and veQTLs using a stepwise regression procedure^28^. For the *trans*-QTL analysis (both for intra-mean and variance), we tested all the SNPs located > 1Mb away from the gene body centre and matched the results with 64 candidate deQTL-dGenes pairs in each cell type. Significant *trans*-QTL is defined at nominal *p*-value < 5 x 10^-8^. We further fit the genotype of those *trans*-veQTLs in the *cis-*veQTL or *cis-*deQTL association model to see if the estimates will be significantly changed.

### Pseudotime trajectory of intra-individual variance and interaction tests

To understand the context-dependent effect of the prioritised deQTLs on the cell state, we estimated the pseudotime of each cell in inferred B cells (B_IN_ + B_MEM_). We used SCTransform^55^ to calculate the scaled expression Pearson residuals using the top 500 highly variable genes and fitted the percentage of mitochondrial expression and experimental pools as covariates. After transformation, we calculate the principal components (PCs) of the expression matrix and constructed the UMAP using the first 30 PCs by RunUMAP() function built in Seurat. We then used PHATE^62^ to estimate the quantitative indicator of cell state (i.e., pseudotime) in a two-dimensional space for each cell. For each individual, we computed the mean pseudotime across all cells and created a mean pseudotime trait. Then the mean pseudotime is tested as an interaction term (G x E) in the QTL association model (for mean, variance, and dispersion separately) in TensorQTL software. The sex or age was also tested for the interaction effect. The nominal *p*-value is first corrected by multiple testing based on the effective number of independent variants in the cis-window, and then converted to Benjamini-Hochberg adjusted *p*-value for each chromosome.

### Identifying co-eQTLs for *RPS26* and its transcription factors

We subset the SCTransformed count matrix for *RPS26* and six transcription factors per cell type, and calculate the Spearman’s correlation between *RPS26* and each of the TF gene across cells within each individual. For each TF gene, we have a co-expression estimate for every individual as the phenotype and run a linear regression of the co-expression phenotype on the genotype of rs1131017. The nominal *p*-values are then converted to Benjamini Hochberg FDR and the test with FDR < 0.05 will be deemed as significant co-eQTL.

### Replication in an independent cohort of non-European ancestry

To replicate our findings of sc-deQTLs in OneK1K of European (EUR) ancestry, we utilised another single-cell cohort from Perez et al.^49^. We conducted sc-deQTL mapping for the individuals of East Asian (EAS) ancestry (97 individuals including 75 healthy controls and 22 lupus patients). The single-cell gene expression was processed in the protocols we used for OneK1K. Intra-individual dispersion of gene expression was also estimated per gene per cell type. For the sc-deQTL mapping, covariates were adjusted in the association model including sex, age, batch, first 6 PCs, first 2 PEER factors, and lupus disease status. Given the difference in SNP panels between two datasets, we only investigate the 34 genes with significant sc-deQTLs in OneK1K cohort in the replication cohort.

## Declarations

### Data and materials availability

All data analysed in the study is available on the Gene Expression Omnibus (GSE196830) and also the HCA data science platform (https://cellxgene.cziscience.com/collections/dde06e0f-ab3b-46be-96a2-a8082383c4a1). The Seurat object of the raw and normalized single-cell gene expression matrix are deposited in the Dropbox (download link available at the Github page below).

### Code availability

The analysis code will be available prior to acceptance on https://github.com/powellgenomicslab/sc-veQTL

### Author contribution

JEP conceived the idea of the project. AX performed the computational analysis with the assistance from SY, JAH, AC, and AS. A.W.H. contributes to the data collection and provides constructive comments. AX and JEP wrote the manuscript with the participation of all authors. All authors read and approved the final manuscript.

