## Supplementary Material for "Genetic variants associated with cell-type-specific intra-individual gene expression variability reveal new mechanisms of genome regulation"

### Contents

Supplementary Notes 1-4
Supplementary Figures 1-15

**Supplementary Note 1**

**Dissecting the mean-variance relationship in the single-cell gene expression data**

We tried to quantify the relationship by visualize the Spearman's correlation coefficient ($\rho_S$) between within-individual mean and variance for all the genes. The mean and variance were calculated based on the SCTransformed counts per individual per gene for each cell type. We observed that the median $\rho_S$ ranged from 0.938 ~ 0.984 across 14 cell types (**Figure 2A**). Notably, there exist an obvious drop-off in the mean-correlation curve at the mean ranging from 0.01 to 0.1 (**Supplementary Figure 2**). Based on this observation, we predict that many of the identified veQTLs might also be significant in the eQTL association test. Seyhan et al. (2021) performed the eQTL analysis and identified 25,606 eQTLs that showed mean effects on the gene expression within individuals in a cell-type-specific manner in the OneK1K cohort. Since this number is based on Spearman's rank correlation analysis, for fair comparison, we reran the analysis using the Matrix eQTL and compared the results with veQTL signals (**Methods**).

**Supplementary Note 2**

**Comparison of different dispersion indices**

There are multiple ways to calculate the dispersion of the within-individual gene expression across cells. The most common non-parametric indices are Fano factor, coefficient of variation (CV), and $CV^2$.

$$Fano\ factor = \frac{\sigma^2}{\mu}$$

$$CV = \frac{\sigma}{\mu}$$

$$CV^2 = \frac{\sigma^2}{\mu^2}$$

From the equation we know that these indices still preserve a negative relationship with the mean. Given a negative binomial (NB) distribution where $\sigma^2 = \mu + \mu^2 \times \theta$, we can further derive these indices as:

$$Fano\ factor = 1 + \mu \times \theta$$

$$CV = \sqrt{\frac{1}{\mu} + \theta}$$

$$CV^2 = \frac{1}{\mu} + \theta$$

From the derived equation, Fano factor is linearly correlated with the mean given fixed dispersion. The CV and $CV^2$ would become extremely large when the mean is small, which is very common in single-cell data. Based on these properties, none of these indices are expected to be good indicators for dispersion.

Another index to represent the dispersion is using the residuals from linear regression of the variance on the mean of each individual. Several studies have adopted this strategy before.

However, there are three main limitations of this strategy. First, the strategy assumes a linear relationship between mean and variance but from the property of NB distribution the assumption is obviously violated. Second, if there is a true biological relationship between the mean and dispersion, this residual strategy will artificially remove it. Third, when the mean is very small give a NB distribution, the variance will be very close to the mean (at the meantime many intra-individual means will be zero under this scenario); thus, the linear regression will suffer a collinearity problem and the residuals will be heavily skewed.

**Supplementary Note 3**

**The special case of estimating dispersion when mean is small**

We have shown the moment estimators for mean and variance for NB distribution. It is easy to get moment estimate for dispersion $\theta = \frac{\sigma^2-\mu}{\mu^2}$. The moment estimator for dispersion will become biased when mean is very small and/or mean and variance are very close. As we know that when $\theta \to 0$, the NB distribution will be approximately equivalent to Poisson distribution. On the other hand, when mean is tiny, the difference between mean and variance will be also small even the dispersion is relatively large. This scenario will be frequently observed when mean < 1 in empirical analysis.

In OneK1K scRNA-seq data, we observe many genes with equal or very similar mean and variance. After looking into the actual distribution, most of the cases are from low mean genes. For example, out of 24,903 genes in $CD4_{NC}$ cells, there are 3,609 genes with equal mean and variance for all 980 individuals (including 539 individuals with all zeros). The non-zero mean estimates of these genes range from [2.94E-6, 1.62E-4], and the proportion of zero expression among individual range from [0.9286, 0.9990]. For those 3,609 genes, the distribution of gene expression within an individual is a special case. Their counts are either all equal to 0, or all 0s but with one 1. These two cases will lead to equal mean and variance. However, this does not necessarily mean that the original distribution is Poisson. Here is a very simple simulation to prove it.

We simulate 10,000 individuals each with 1,000 cells, and all the within-individual distributions are drawn from $NB \sim (\mu = 0.001, \theta = 1)$. The distribution patterns of these individuals are displayed in details (**Supplementary Table 8**). We can see the first four patterns account for ~98.2% distribution pattern. Also, when there is only 0 and 1, the variance is also no larger than mean and so the moment estimate for theta is always zero or negative. This can be easily proven as bellow:

Let $Fano\ factor\ = \frac{\sigma^2}{\mu} = \frac{m}{m+k-1}$, where *m* is number of 0s and *k* is an integer>1. This index is always smaller than 1 and larger than 0, so the variance is always smaller than the mean. Since the expected $\mu$ is small, empirically $m \gg k$, thus the Fano factor will be very close to 1. Also,

Let $\hat{\theta}_{MOE} = \frac{\sigma^2-\mu}{\mu^2} = \frac{1-k}{k} \cdot \frac{m+k}{m+k-1}$. This index is always negative because of the first component. Also, since $m \gg k$, the second component is close to 1. The final estimate will be only slightly larger than $\frac{1-k}{k}$, and the lower boundary is $-\frac{1}{2}$.

**Supplementary Note 4**

**The simulation for dispersion estimation using Cox-Reid adjusted MLE**

To assess the performance of Cox-Reid adjusted MLE for dispersion parameter, we conducted a simulation with different number of cells ($M$), mean ($\mu$), and dispersion ($\theta$). The simulation parameters are as following:

M ∈ (10, 25, 50, 100, 500, 1000, 5000)
μ ∈ (0.001, 0.01, 0.05, 0.1, 0.2, 0.3, 0.4, 0.5, 1, 1.5, 2, 2.5, 3, 5, 10, 30)
θ∈ (0.1, 0.5, 1)

We simulated an expression matrix with 1,000 genes x m cells and each gene follows the same Negative Binomial distribution. Each row is drawn from $X \sim NB(\mu, \theta)$, and the R code is *rnbinom(n = n, size = 1/theta, mu = mu)*. We then used *glmGamPoi::glm_gp()* to estimate the dispersion parameter for all rows simultaneously and obtained the mean of the dispersion estimates across 1,000 rows. The whole process is replicated for 100 times. For each true dispersion parameter, we visualized the simulation results by a boxplot of different means against the dispersion estimates across 100 replicates.

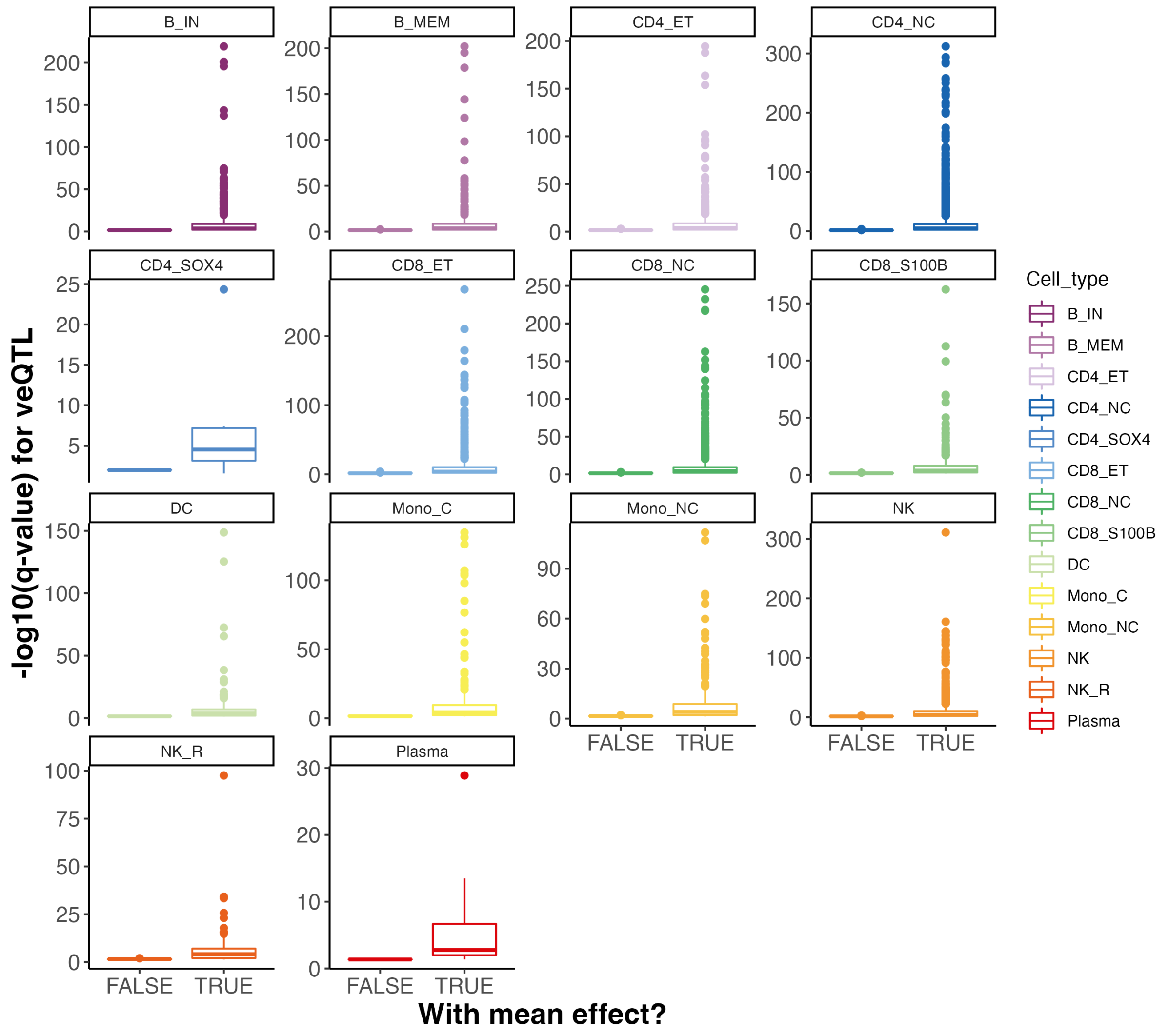


**Supplementary Figure 1. Association significant level of top veQTL with or without mean effects.**

The boxplot shows the distribution of significant level of the top veQTL for each vGene, grouped by whether this vGene is also an eGene. TRUE (FALSE) group means the corresponding vGene is (not) also an eGene. The beta effect size is the slope estimate for the top SNP of the vGene in the veQTL association tests.

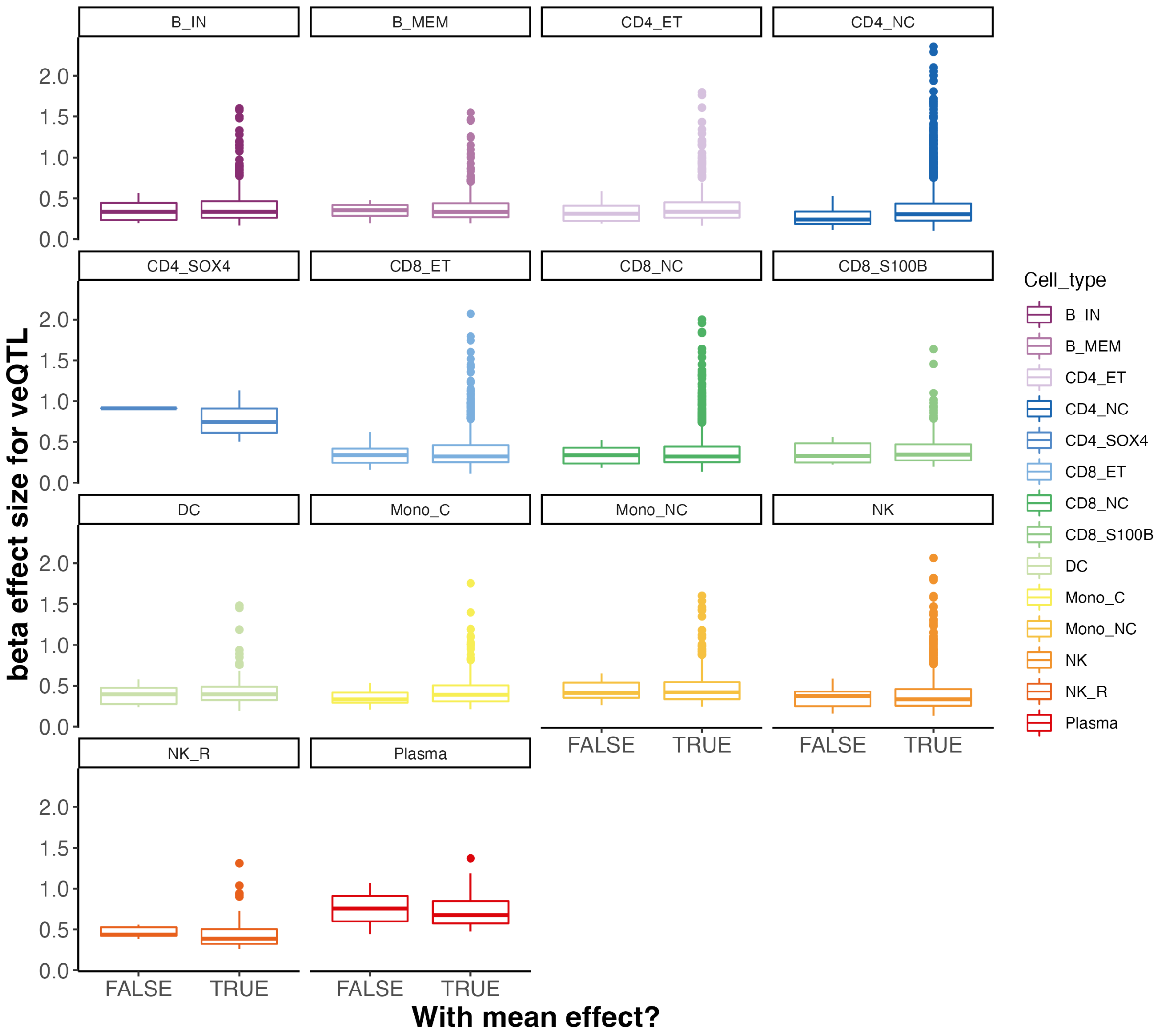


**Supplementary Figure 2. Effect size of top veQTL with or without mean effects.**

The boxplot shows the distribution of beta effect size of the top veQTL for each vGene, grouped by whether this vGene is also an eGene.

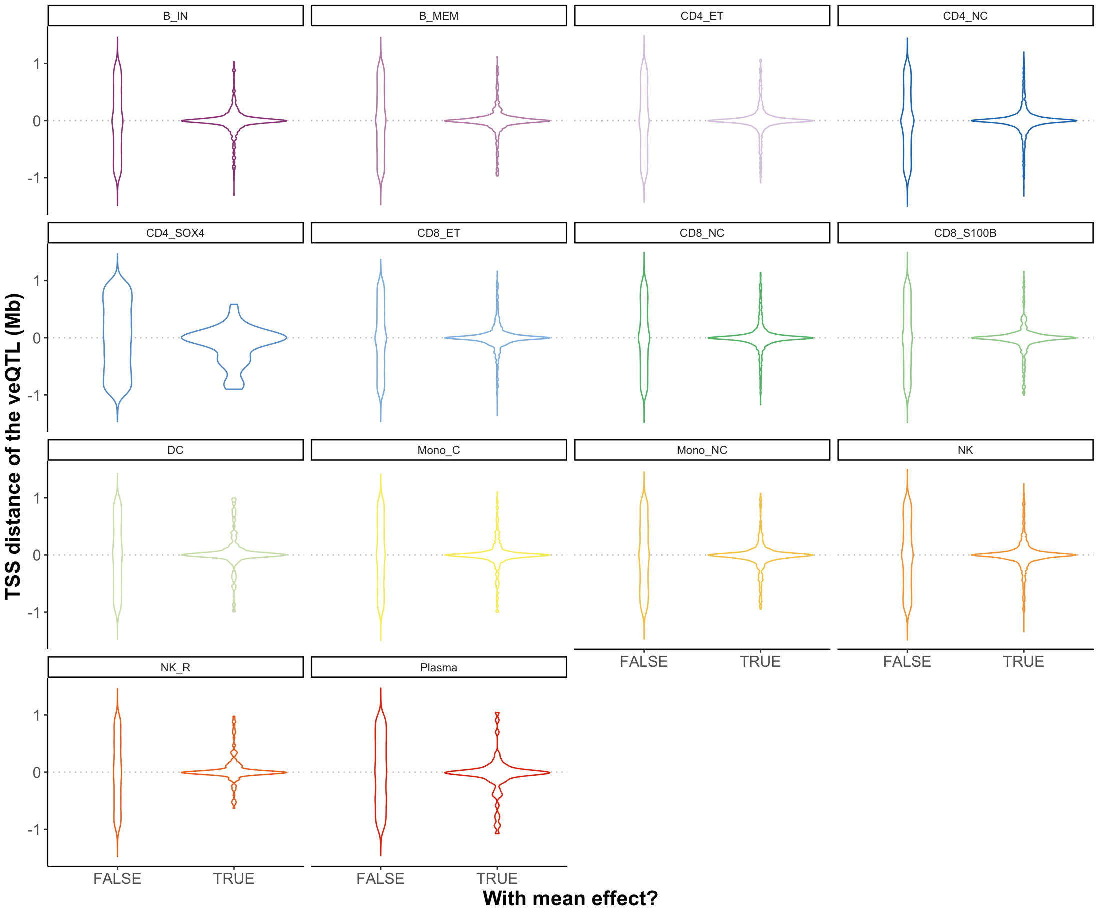


**Supplementary Figure 3. TSS distance top veQTL with or without mean effects.**
This violin plot compares the distance between top veQTL to the transcription start site for those vGenes with or without mean effects. The x-axis indicates the two groups of veQTLs. The y-axis indicates the distance by Mb.

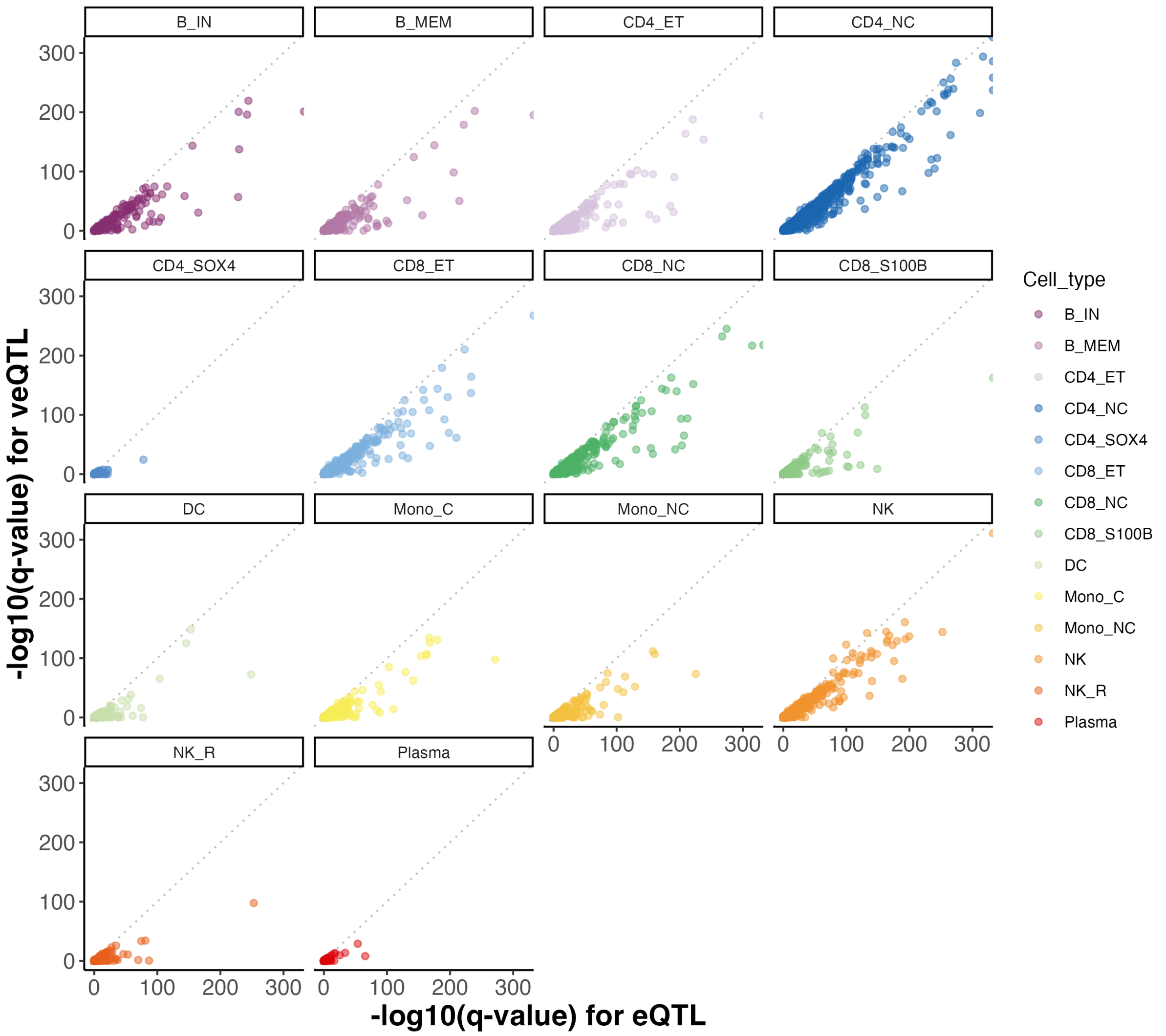


**Supplementary Figure 4. Comparison of association significant level between eQTL and veQTL.**

The x-axis and y-axis denote the -log10($q$-value) for eQTL and veQTL, respectively. Each dot indicates a SNP-gene pair test, and the colour of the dot indicates the cell type. The grey dashed line denotes the diagonal line of the coordinate panel.

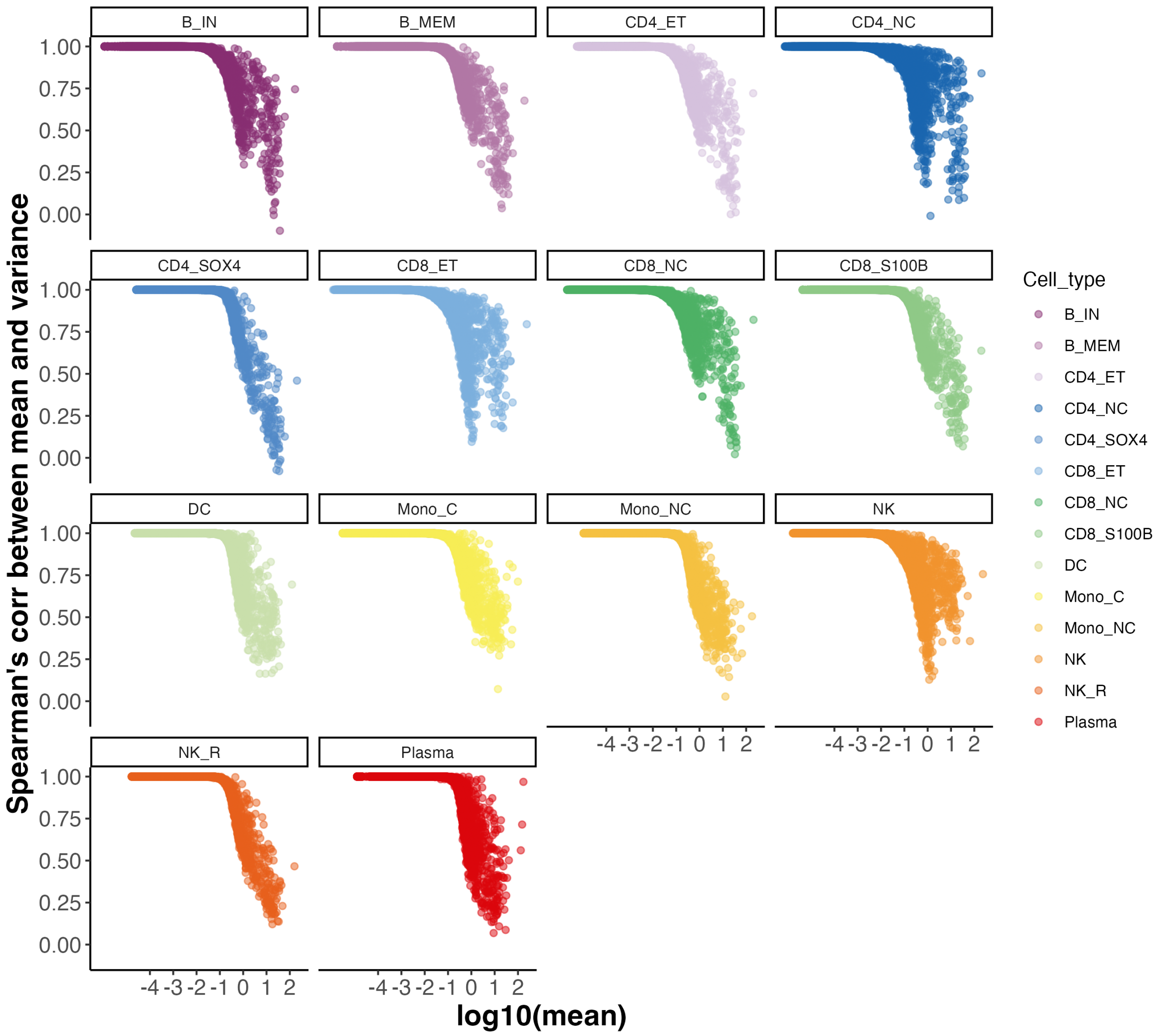


**Supplementary Figure 5. The relationship between the intra-individual mean and the mean-variance correlation.**

The scatter plot shows the relationship between the intra-individual mean and the Spearman's correlation efficient between intra-individual mean and variance across individuals per gene in 14 cell types. Each dot indicates one gene and there are 24,903 genes in total.

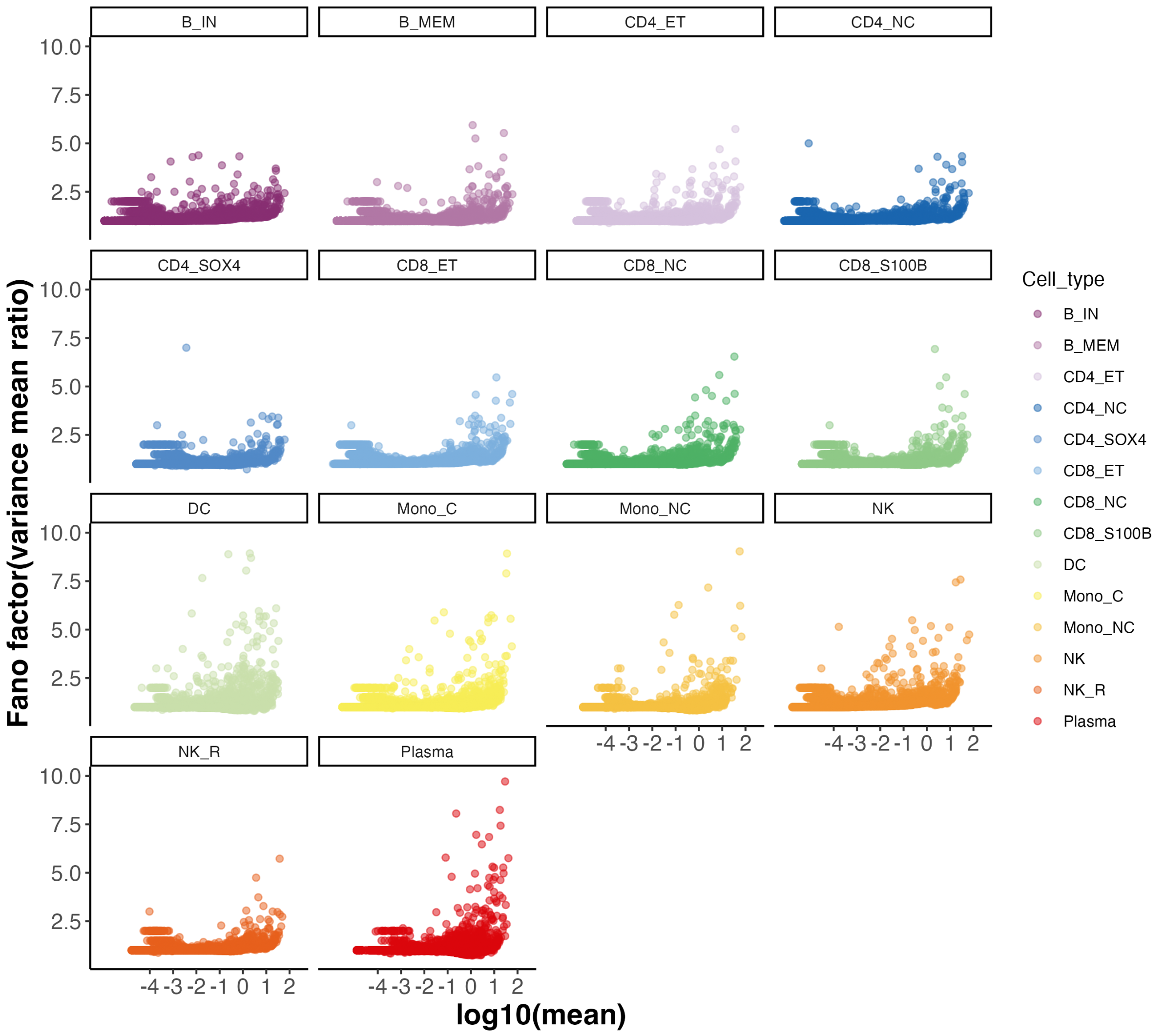


**Supplementary Figure 6. The relationship between the mean and the Fano factor (variance mean ratio).**

The scatter plot shows the relationship between the intra-individual mean and the proportion of non-expressed individuals per gene in 14 cell types. The x-axis denotes the log10 transformed mean per gene. The y-axis denotes the Fano factor, which is calculated as variance divided by mean. Each dot indicates one gene's mean Fano factor across individual. 46 outliers that with Fano factor > 10 were omitted. Genes with all zero expression do not have a Fano factor.

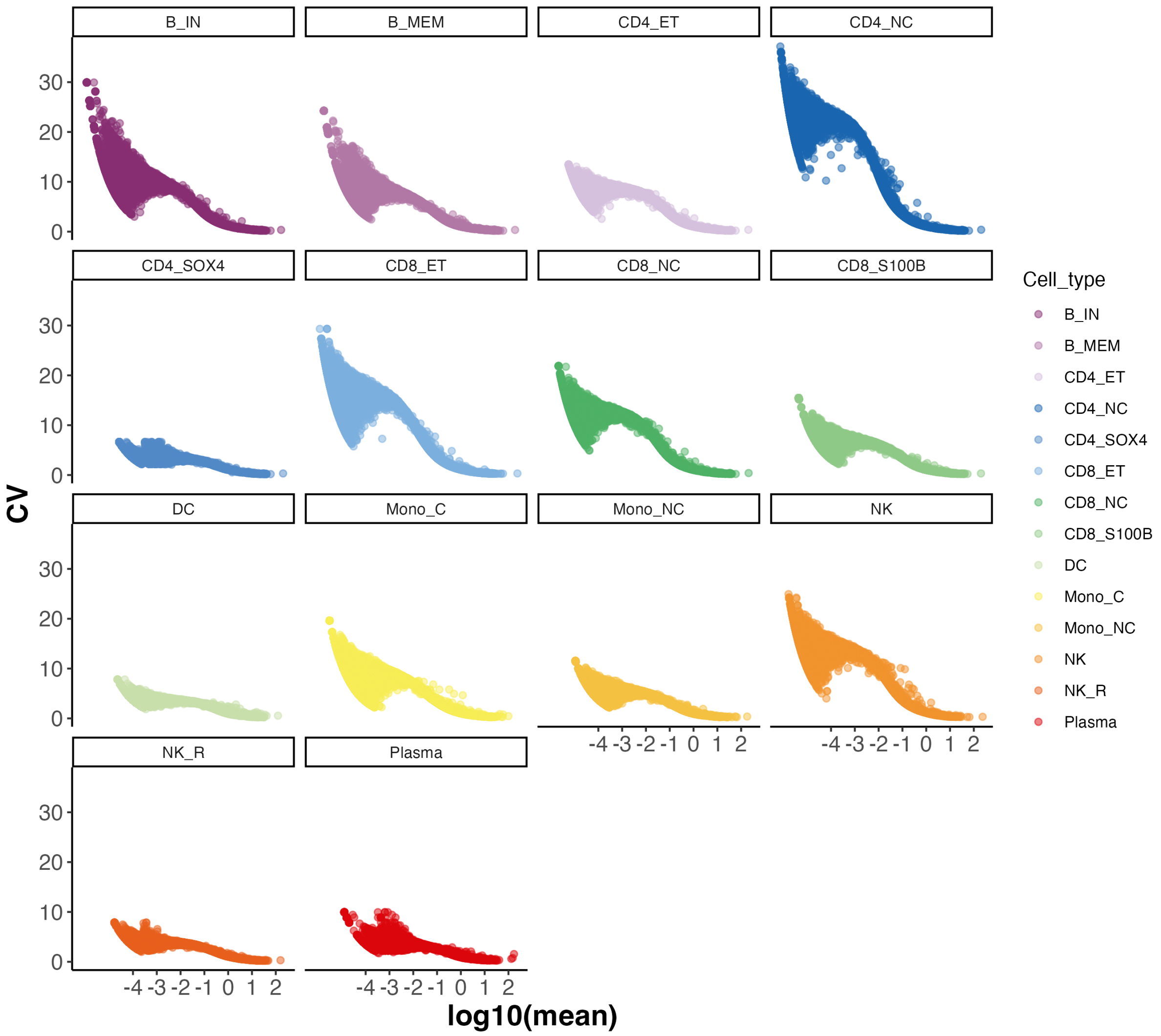


**Supplementary Figure 7. The relationship between the mean and the coefficient of variation (CV).**

The scatter plot shows the relationship between the intra-individual mean and the proportion of non-expressed individuals per gene in 14 cell types. The x-axis denotes the log10 transformed mean per gene. The y-axis denotes the CV (coefficient of variation), which is calculated as standard deviation divided by the mean per gene. Genes with all zero expression do not have a CV estimate.

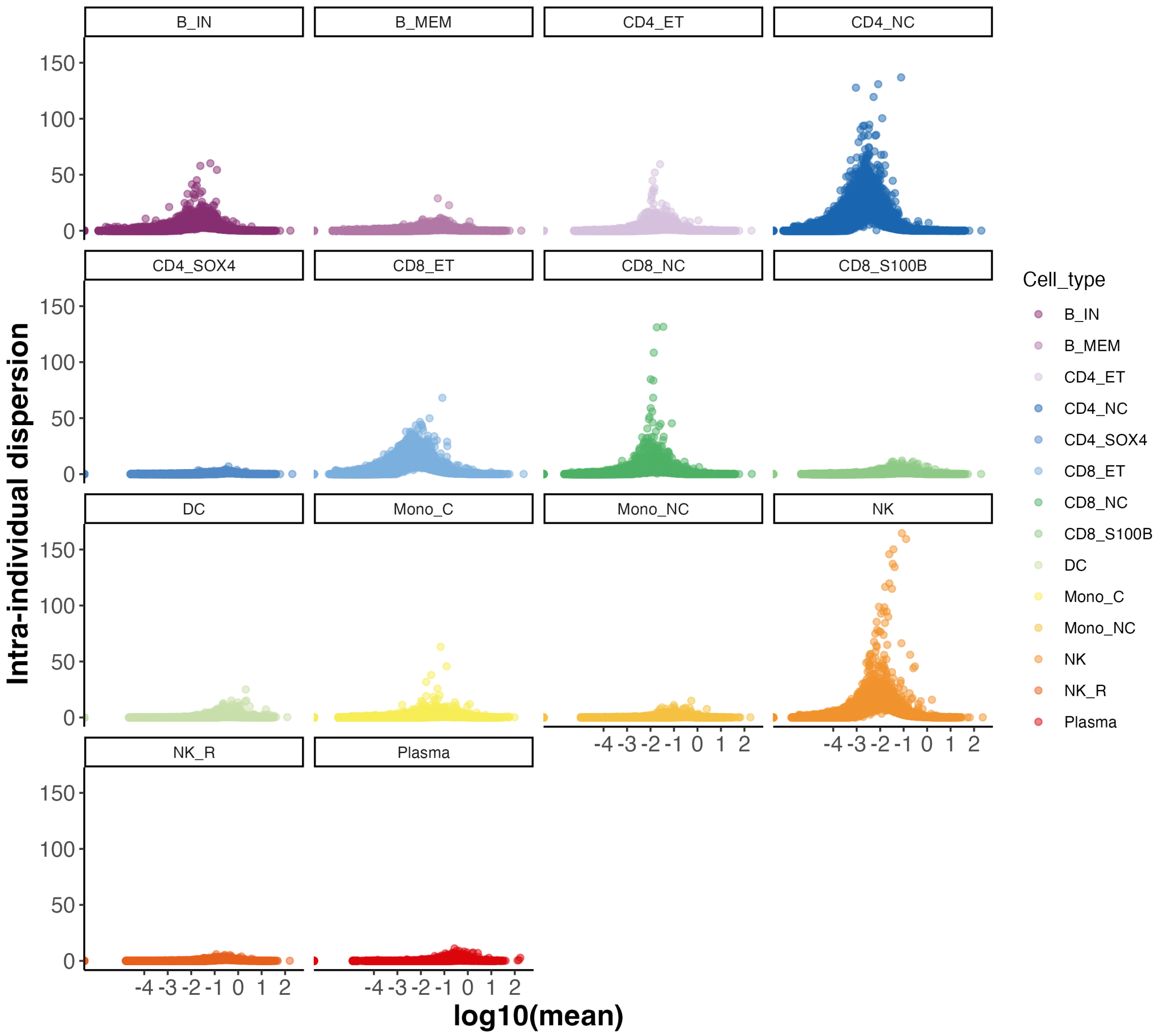


**Supplementary Figure 8. The relationship between the mean and the dispersion.**

The scatter plot shows the relationship between the intra-individual mean and intra-individual dispersion per gene in 14 cell types. The x-axis denotes the log10 transformed mean per gene. The y-axis denotes the intra-individual estimates for dispersion per gene, which is calculated by the Cox-Reid adjusted MLE method.

**Supplementary Figure 9. Genetic association plot for 55 deQTLs.**

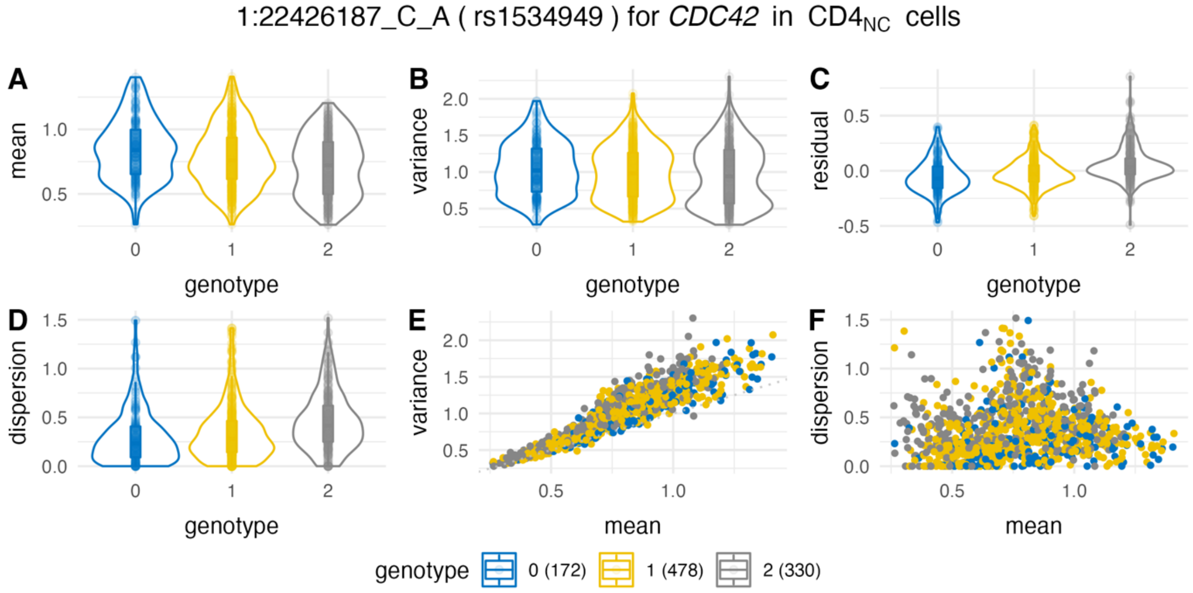


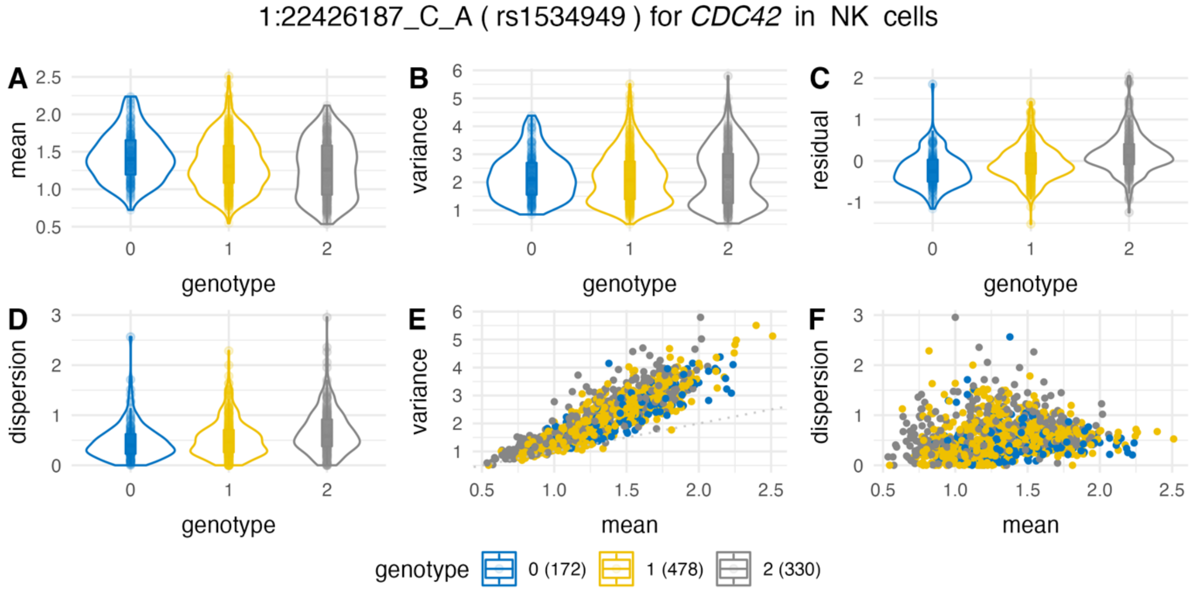

1:26616280_A_G ( rs3924324 ) for *CD52* in $CD4_{NC}$ cells

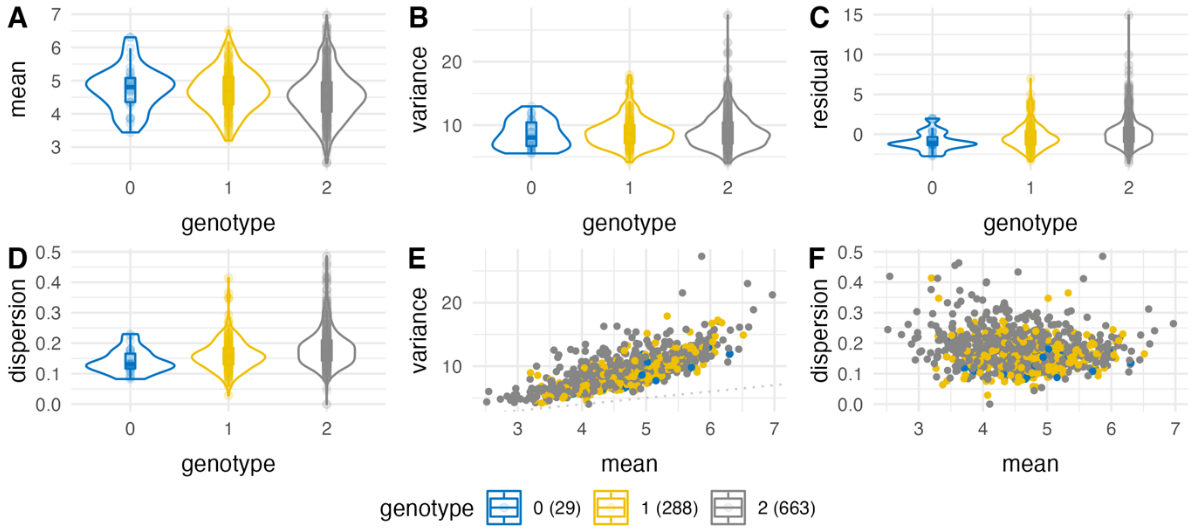


1:26645806_C_T ( rs11589222 ) for *CD52* in $CD8_{ET}$ cells

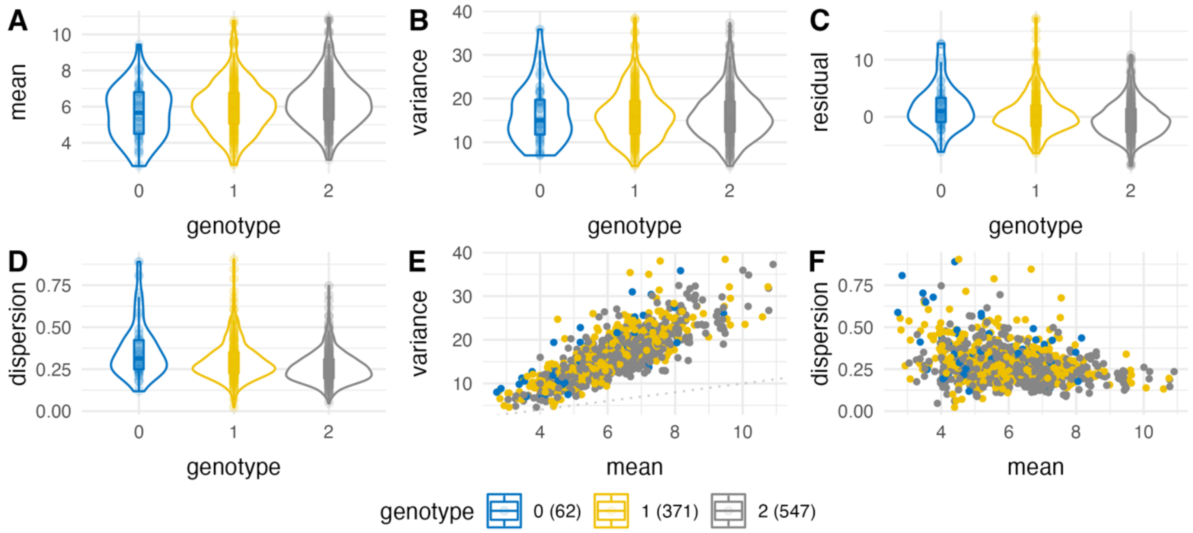


1:153514241_C_A ( rs7535476 ) for *S100A4* in NK cells

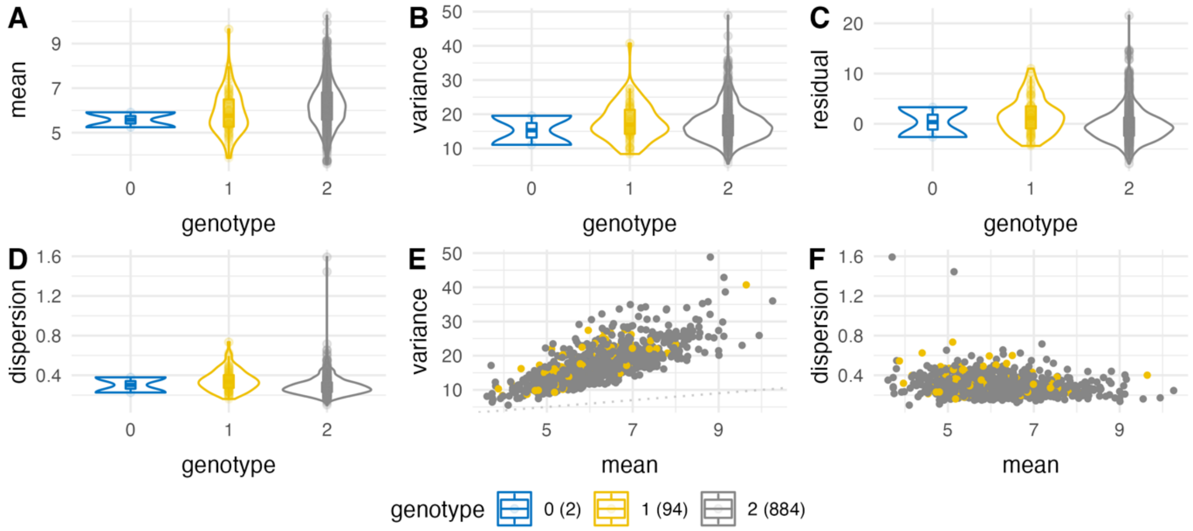

2:85916005_G_A ( rs4832181 ) for *GNLY* in $NK_R$ cells

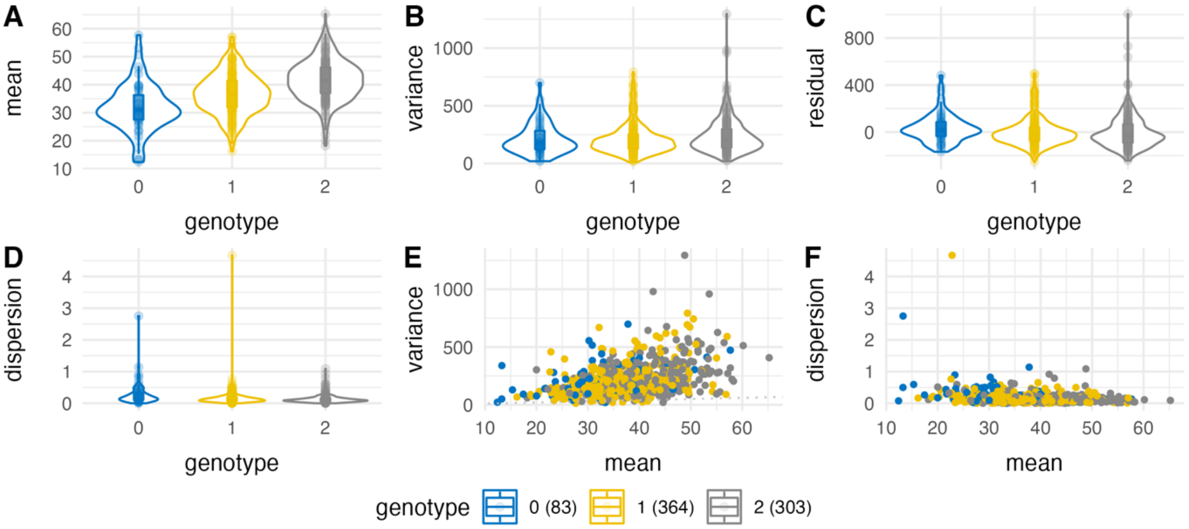


2:85920249_G_T ( rs3755007 ) for *GNLY* in NK cells

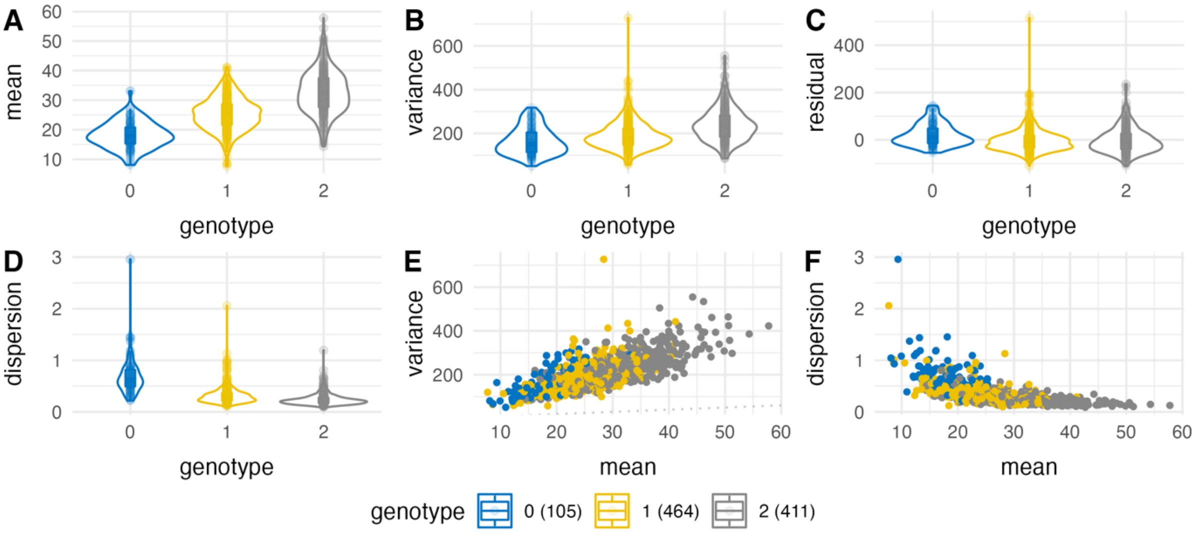


2:85934499_A_C ( rs12151621 ) for *GNLY* in $CD8_{ET}$ cells

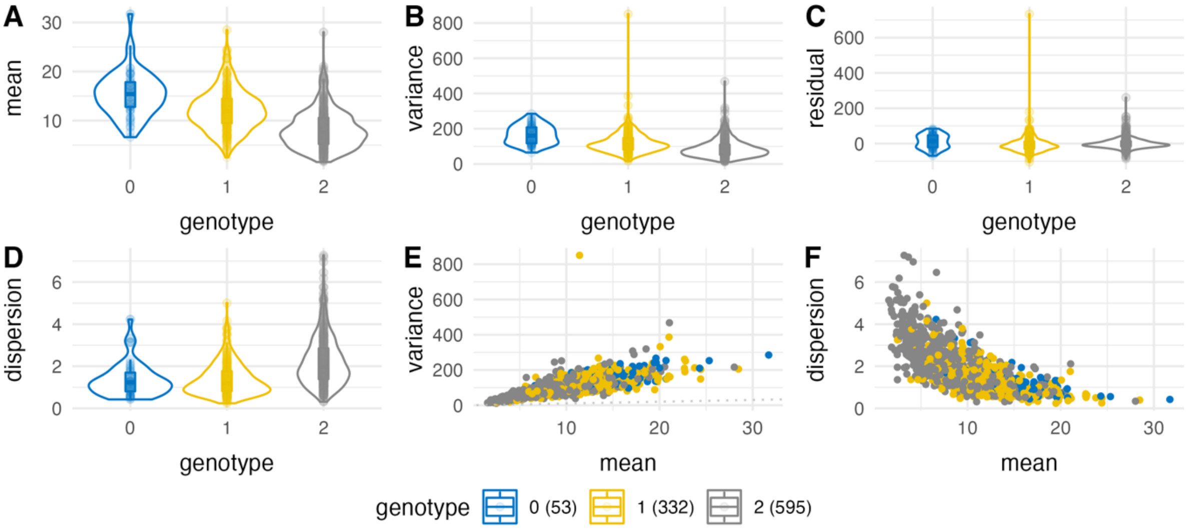

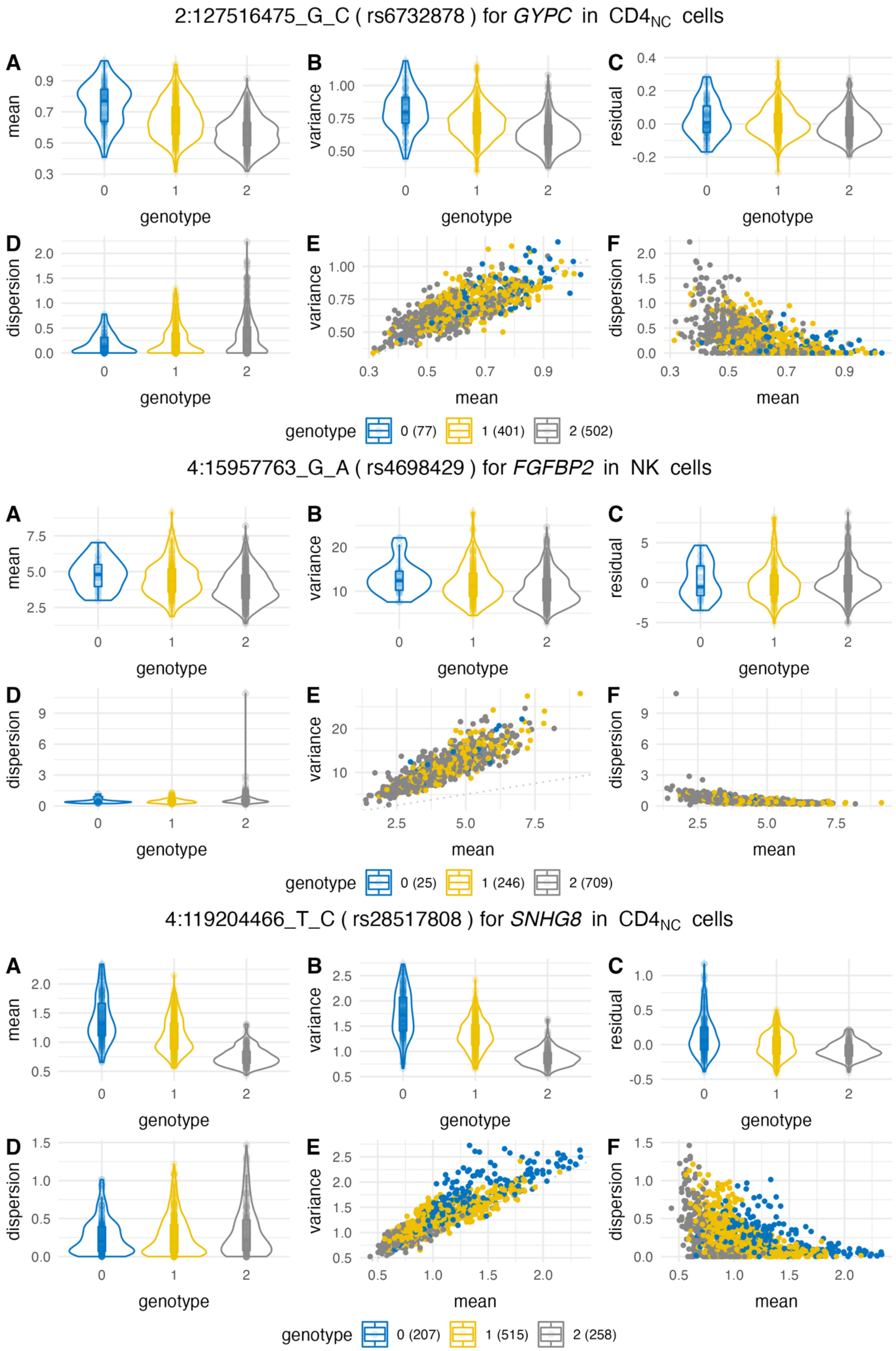
2:127516475_G_C ( rs6732878 ) for GYPC in CD4NC cells
A
B
C
D
E
F
mean
variance
residual
dispersion
genotype
genotype 0 (77) 1 (401) 2 (502)
4:15957763_G_A ( rs4698429 ) for FGFBP2 in NK cells
A
B
C
D
E
F
mean
variance
residual
dispersion
genotype
genotype 0 (25) 1 (246) 2 (709)
4:119204466_T_C ( rs28517808 ) for SNHG8 in CD4NC cells
A
B
C
D
E
F
mean
variance
residual
dispersion
genotype
genotype 0 (207) 1 (515) 2 (258)

5:178986632_C_A ( rs7703730 ) for *HNRNPH1* in $CD4_{NC}$ cells

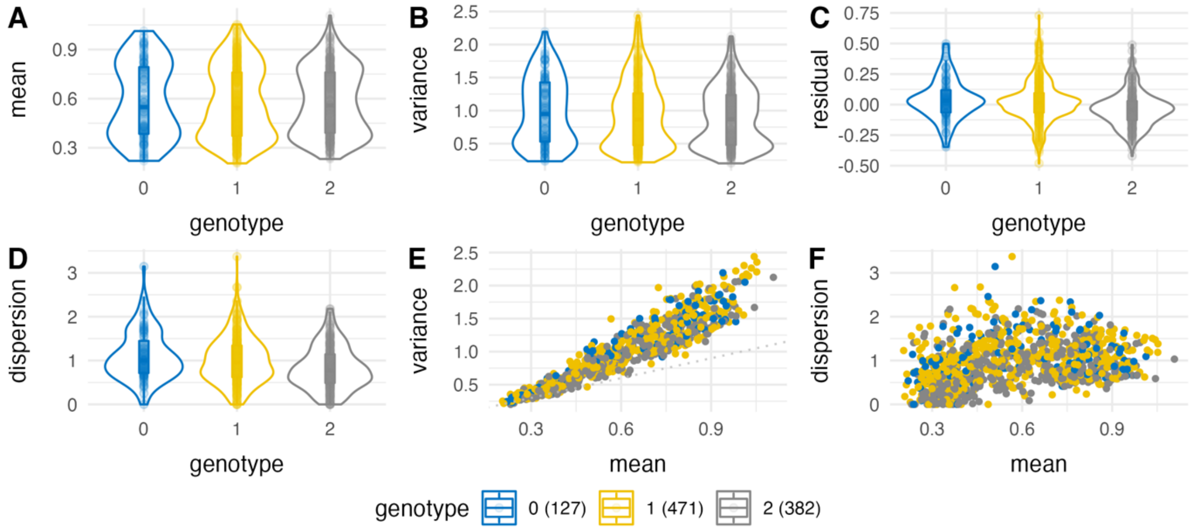


6:29913266_T_G ( rs1061156 ) for *HLA-A* in $CD4_{NC}$ cells

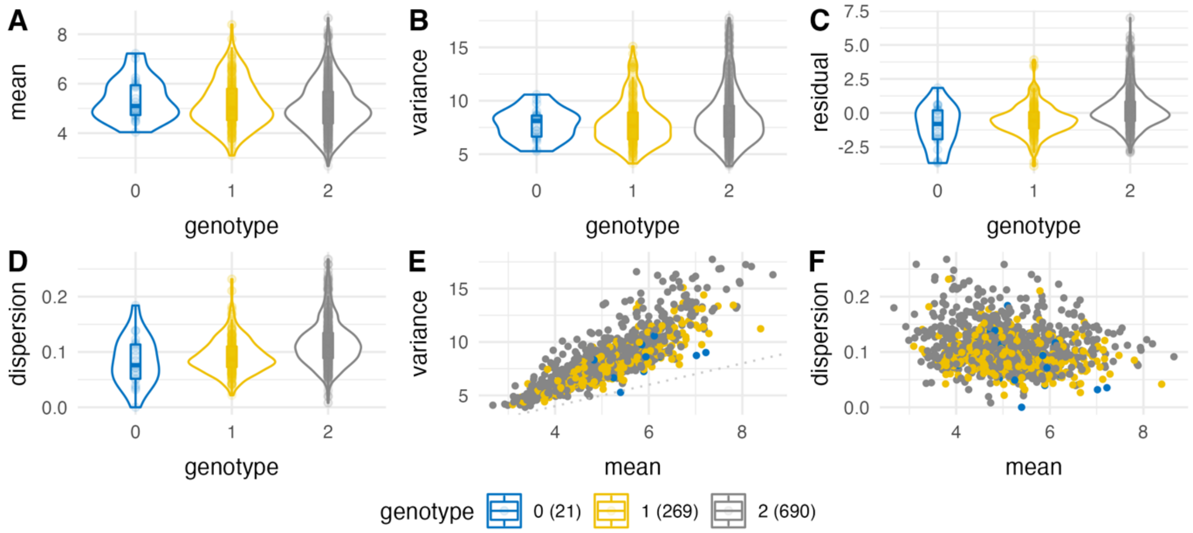


6:31221914_C_T ( rs9264219 ) for *HLA-C* in $CD4_{NC}$ cells

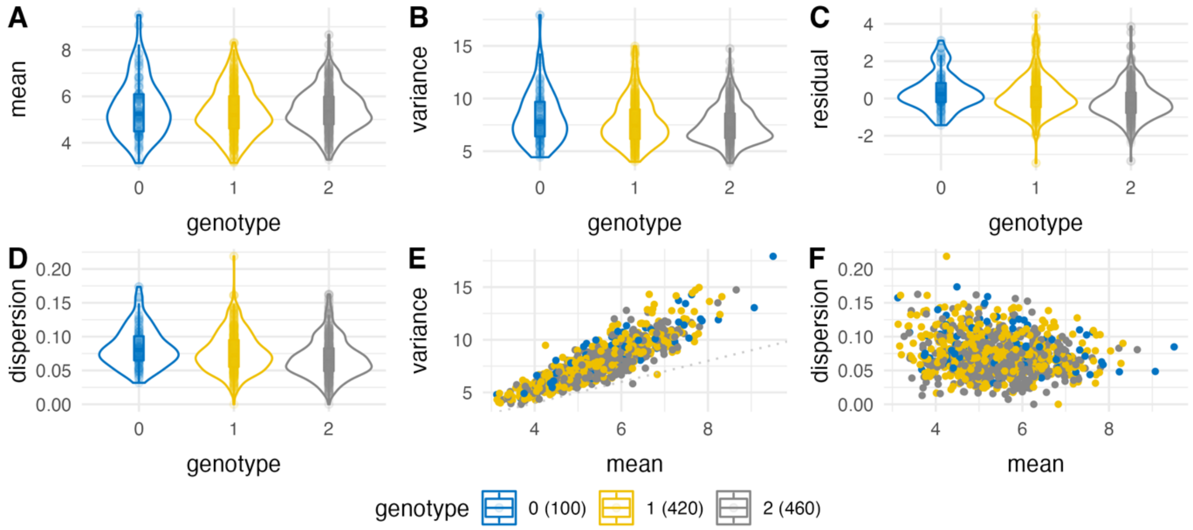

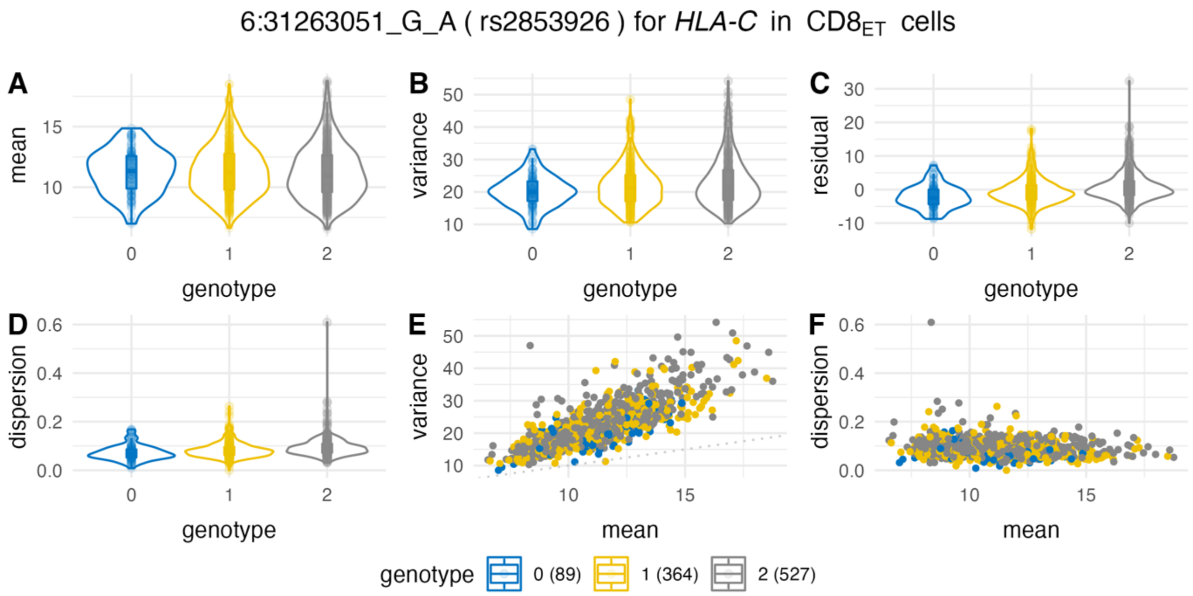
6:31263051_G_A ( rs2853926 ) for HLA-C in CD8ET cells
A
mean
genotype
B
variance
genotype
C
residual
genotype
D
dispersion
genotype
E
variance
mean
F
dispersion
mean
genotype 0 (89) 1 (364) 2 (527)

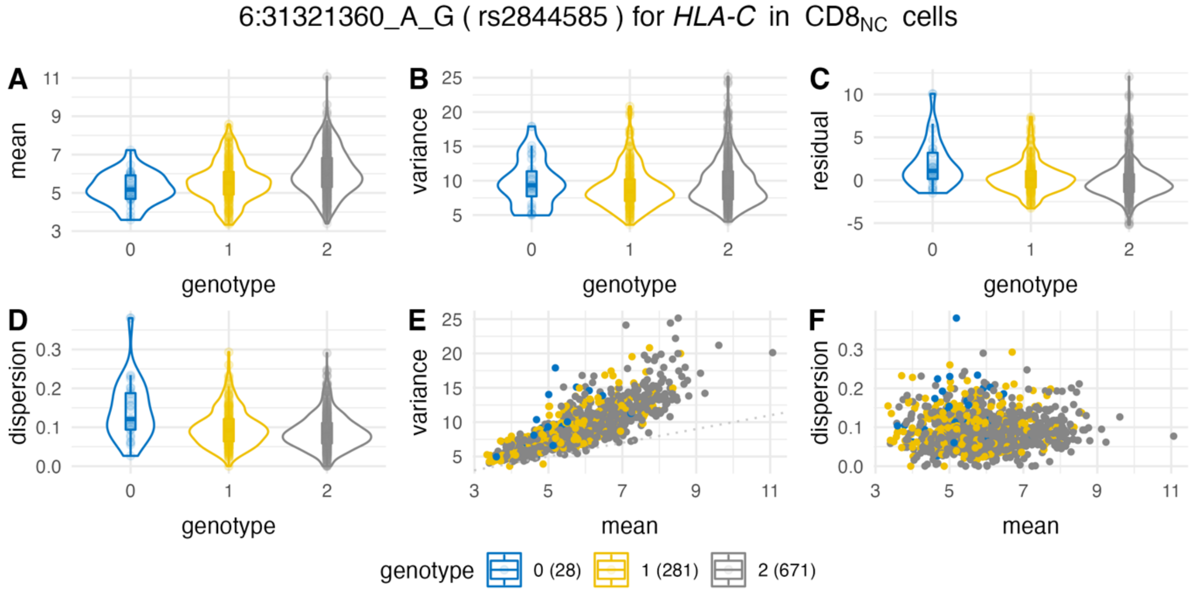
6:31321360_A_G ( rs2844585 ) for HLA-C in CD8NC cells
A
mean
genotype
B
variance
genotype
C
residual
genotype
D
dispersion
genotype
E
variance
mean
F
dispersion
mean
genotype 0 (28) 1 (281) 2 (671)

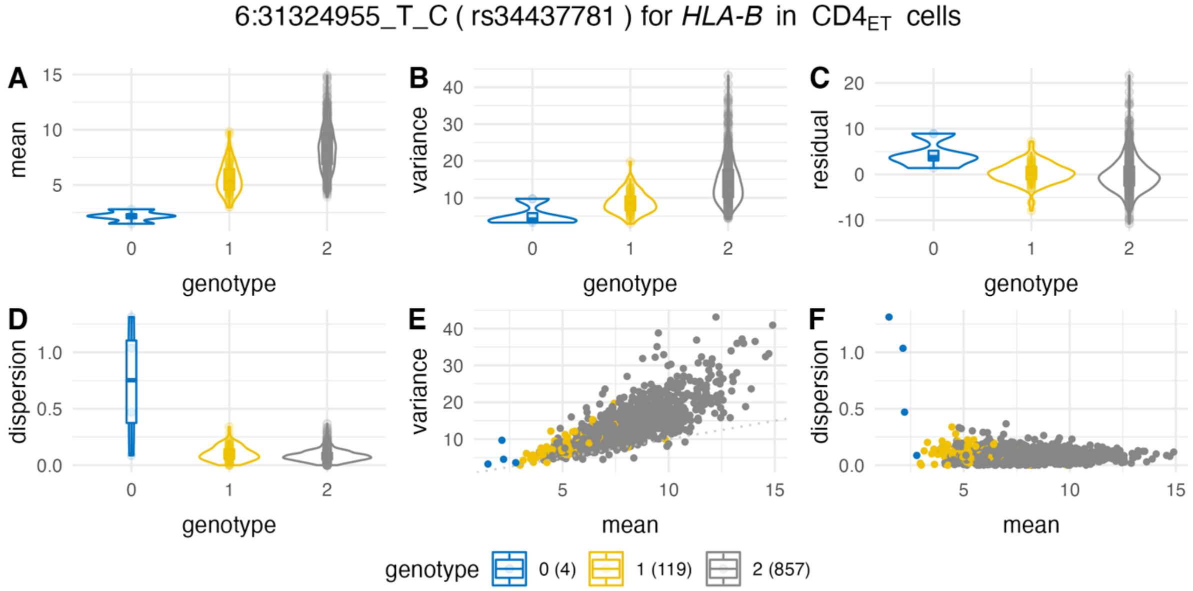
6:31324955_T_C ( rs34437781 ) for HLA-B in CD4ET cells
A
mean
genotype
B
variance
genotype
C
residual
genotype
D
dispersion
genotype
E
variance
mean
F
dispersion
mean
genotype 0 (4) 1 (119) 2 (857)

6:31327701_G_T ( rs9378249 ) for *HLA-B* in CD8$_{NC}$ cells

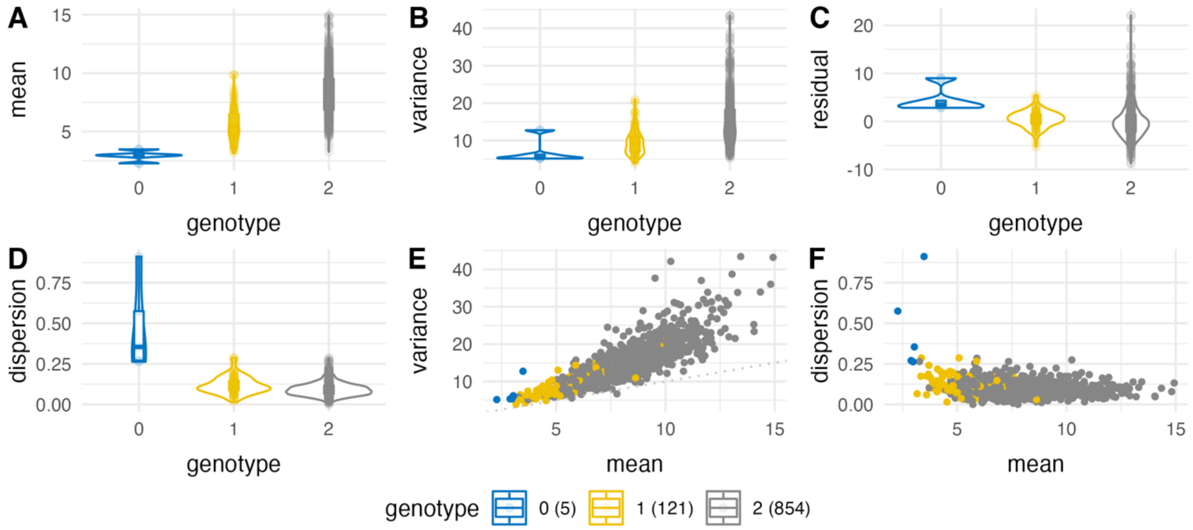


6:31366295_A_C ( rs9394070 ) for *HLA-B* in CD4$_{NC}$ cells

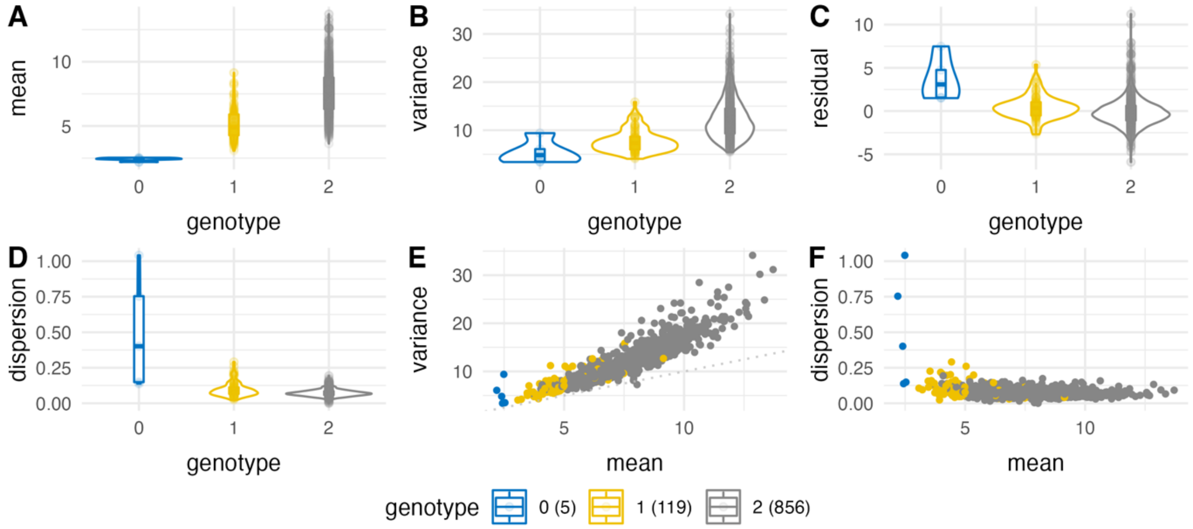


6:32589326_A_C ( rs9271503 ) for *HLA-DQA1* in B$_{MEM}$ cells

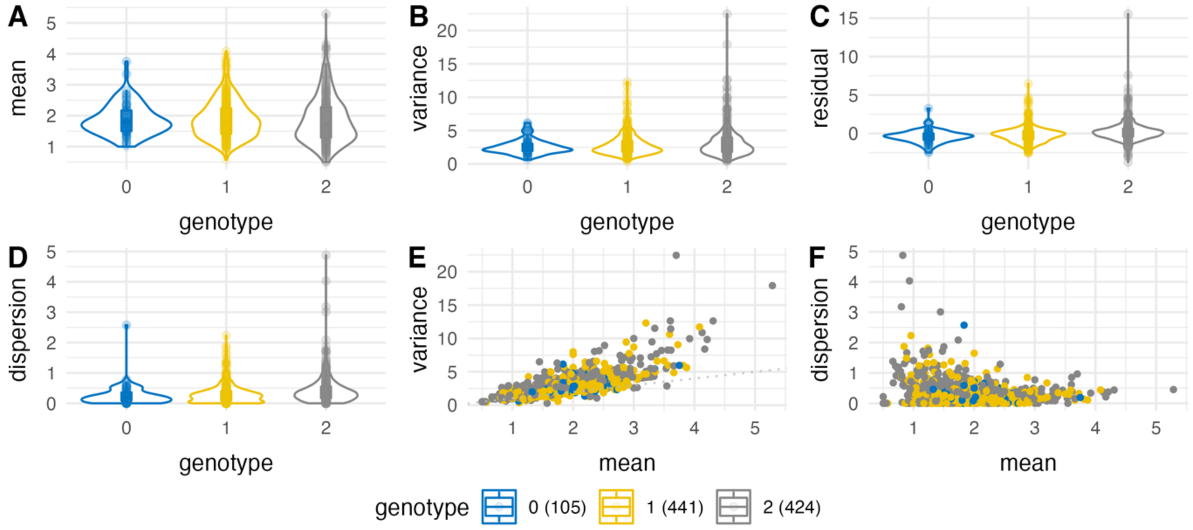

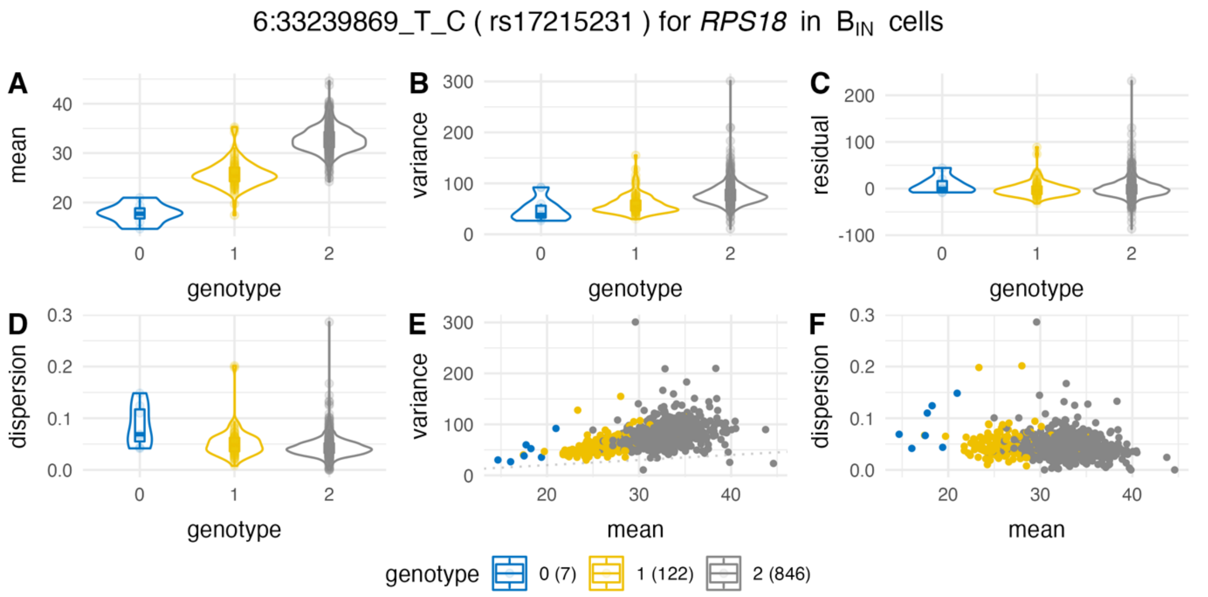

6:33239869_T_C ( rs17215231 ) for *RPS18* in $B_{IN}$ cells
A
B
C
D
E
F
mean
variance
residual
dispersion
genotype
mean
0
1
2
40
30
20
300
200
100
0
200
100
0
-100
0.3
0.2
0.1
0.0
genotype
0 (7)
1 (122)
2 (846)


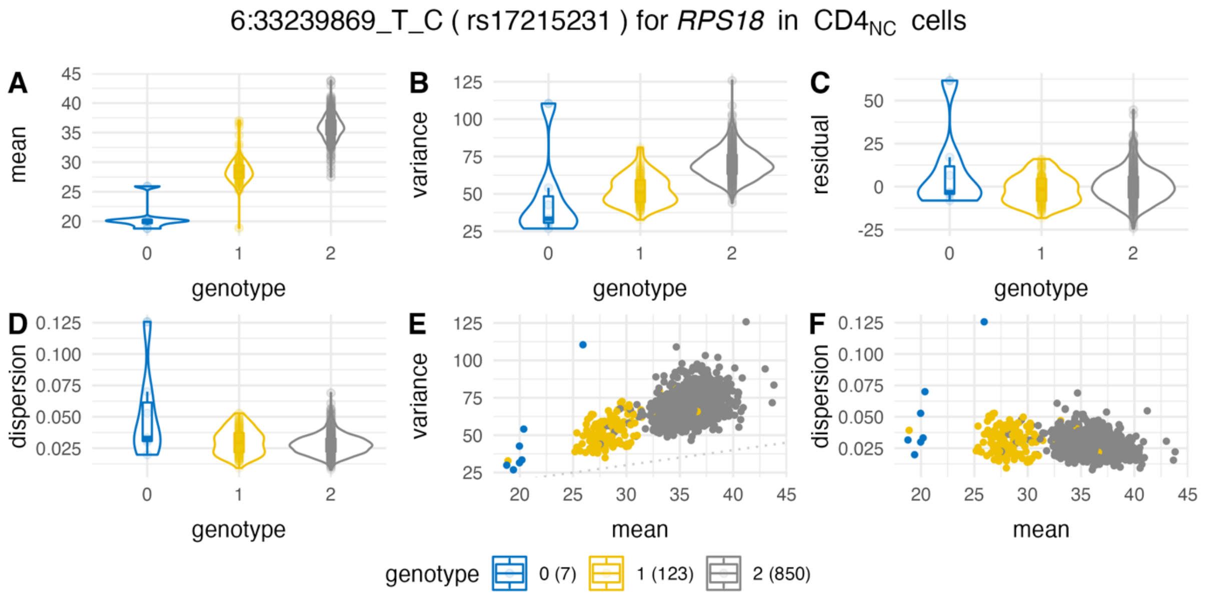

6:33239869_T_C ( rs17215231 ) for *RPS18* in $CD4_{NC}$ cells
A
B
C
D
E
F
mean
variance
residual
dispersion
genotype
mean
0
1
2
45
40
35
30
25
20
125
100
75
50
25
-25
0.125
0.100
0.075
0.050
0.025
genotype
0 (7)
1 (123)
2 (850)


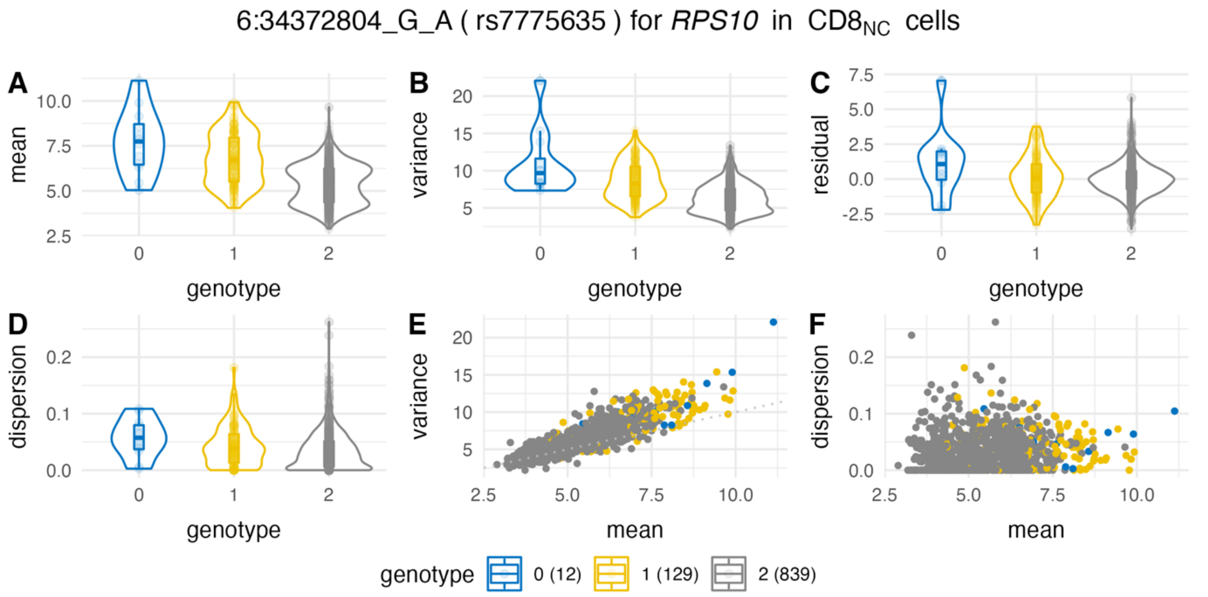

6:34372804_G_A ( rs7775635 ) for *RPS10* in $CD8_{NC}$ cells
A
B
C
D
E
F
mean
variance
residual
dispersion
genotype
mean
0
1
2
10.0
7.5
5.0
2.5
20
15
10
5
0.0
-2.5
0.2
0.1
genotype
0 (12)
1 (129)
2 (839)

6:167370999_G_A ( rs2769346 ) for *RNASET2* in $CD4_{NC}$ cells

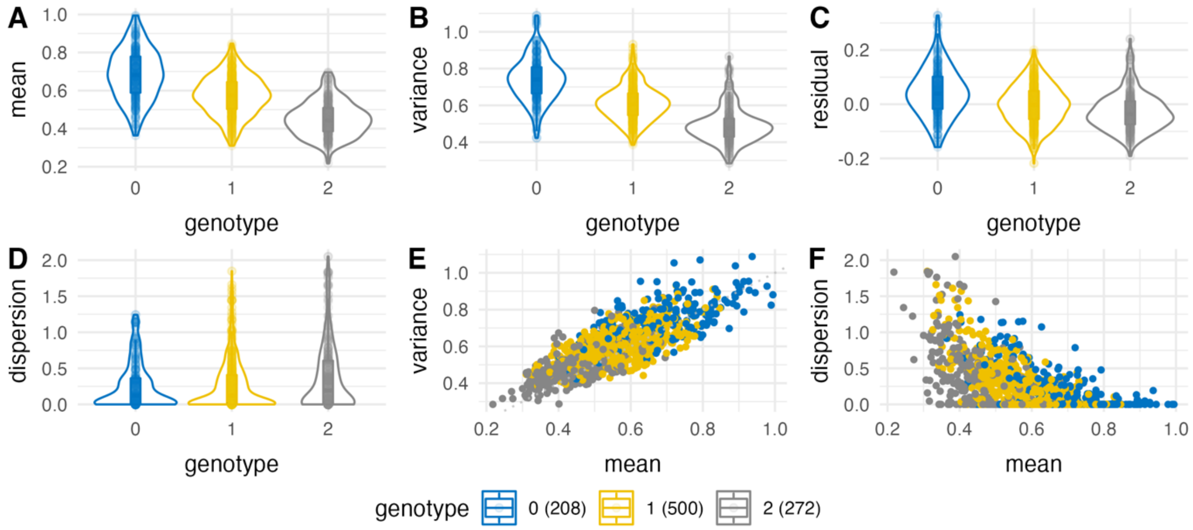


8:144075281_T_C ( rs4424237 ) for *LY6E* in NK cells

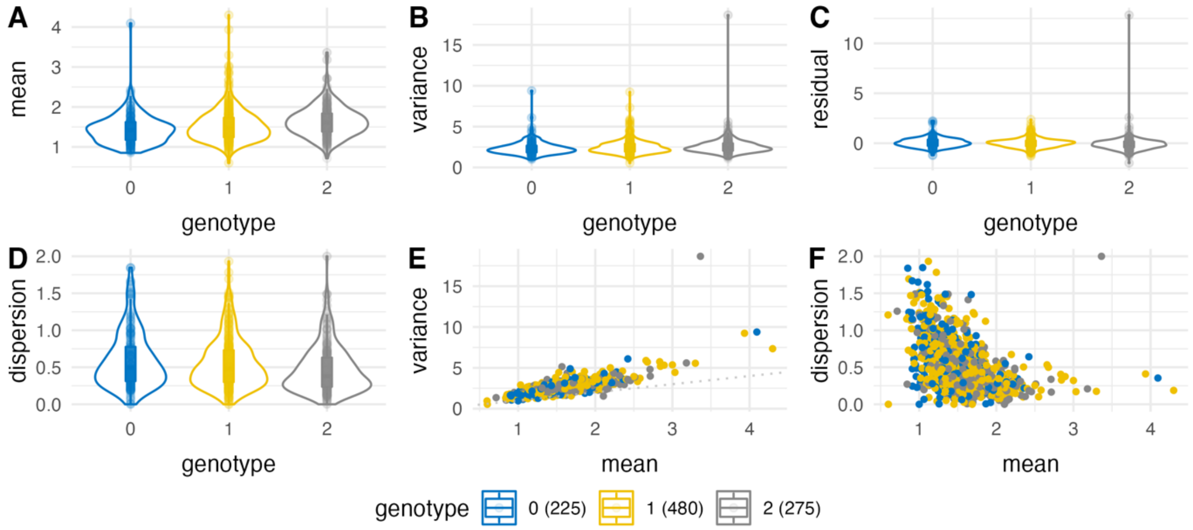


9:75769950_G_C ( rs2795112 ) for *ANXA1* in $CD4_{NC}$ cells

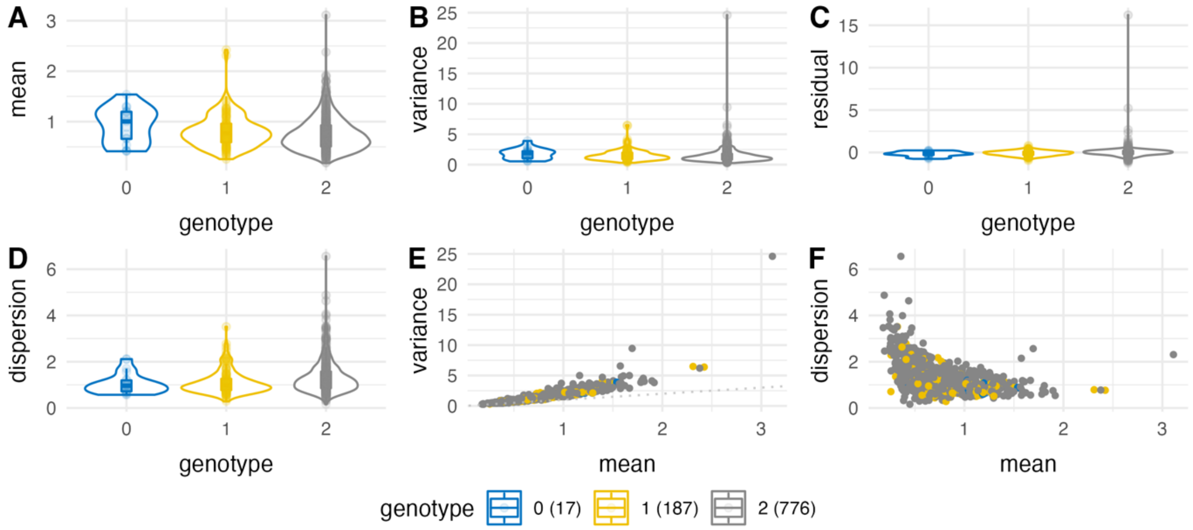

9:139848273_G_A ( rs2271869 ) for *PTGDS* in NK cells

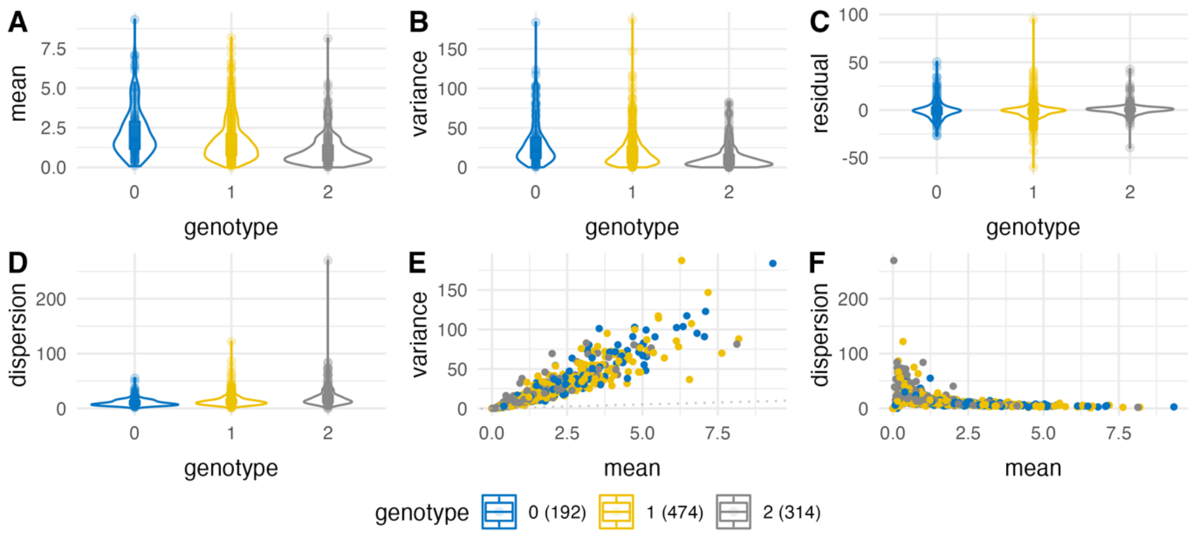


11:303271_A_T ( rs6598046 ) for *IFITM2* in $CD4_{NC}$ cells

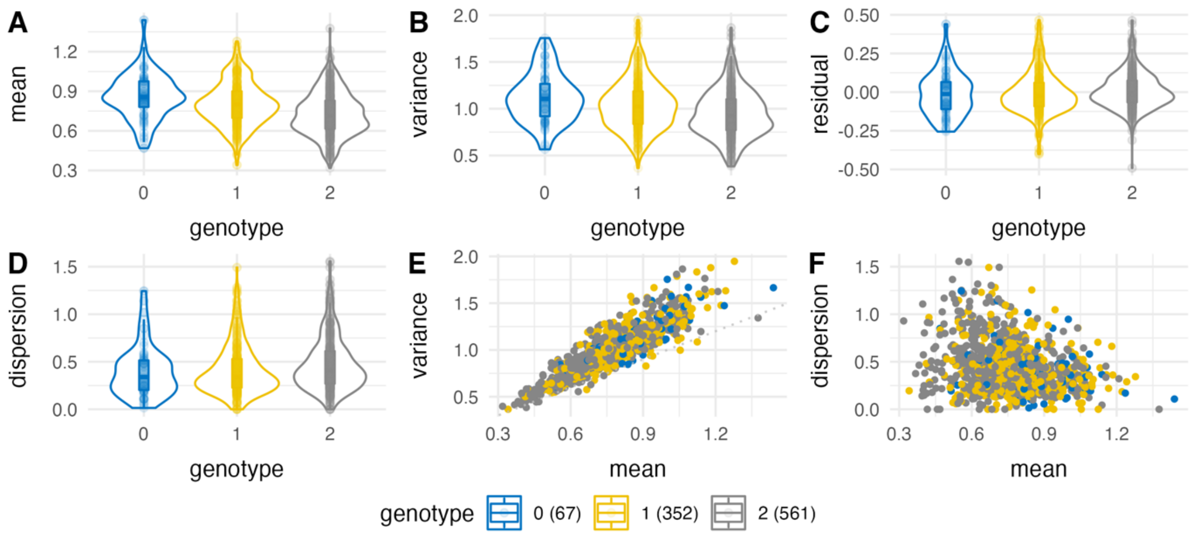


11:307539_A_G ( rs111412325 ) for *IFITM2* in $CD8_{ET}$ cells

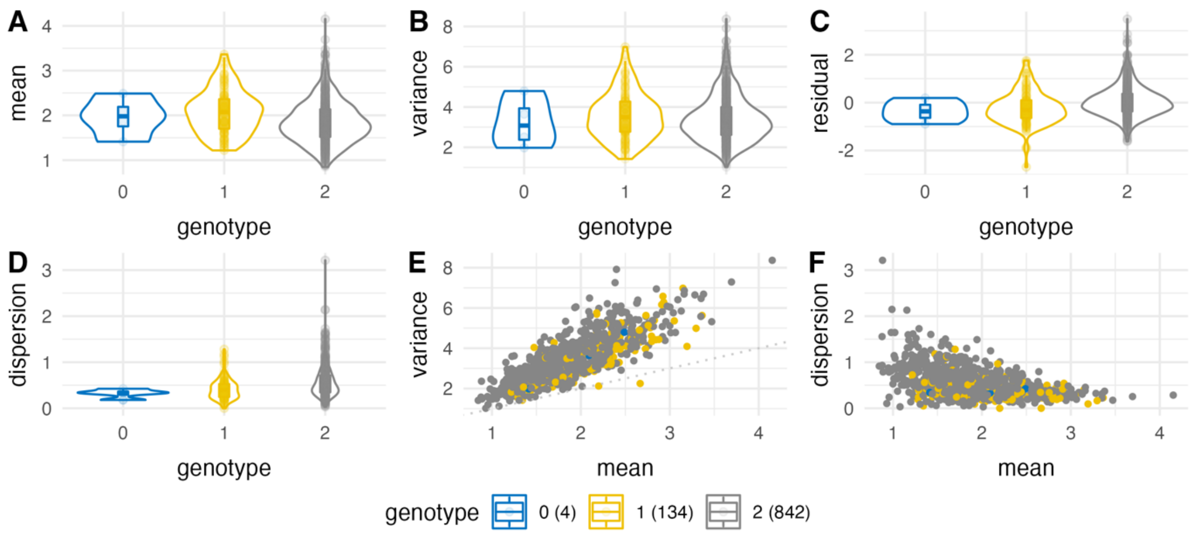

11:349122_C_T ( rs7117996 ) for *IFITM2* in NK cells

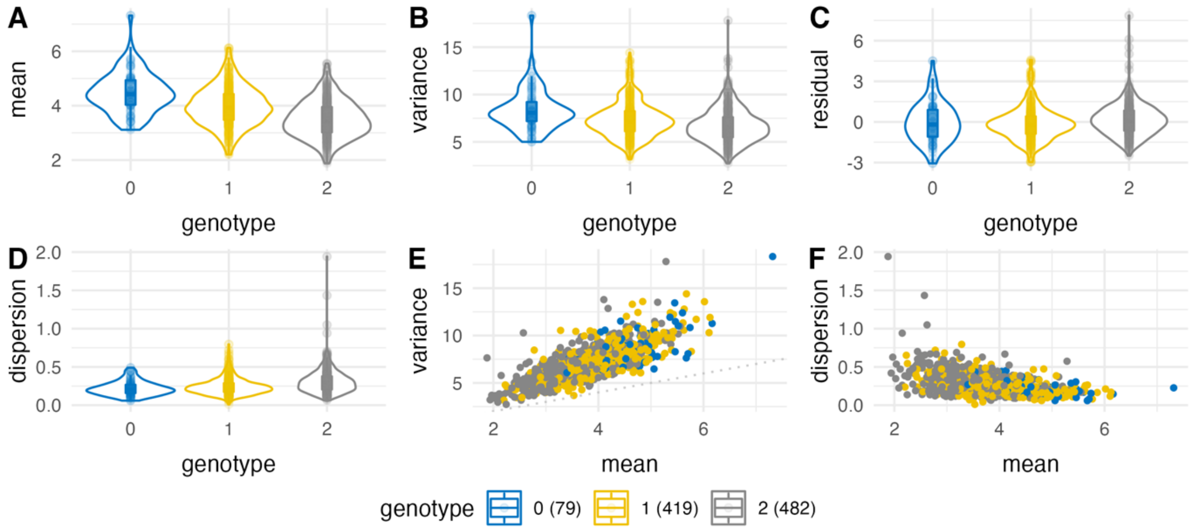


11:802902_T_G ( rs28360884 ) for *RPLP2* in $CD4_{NC}$ cells

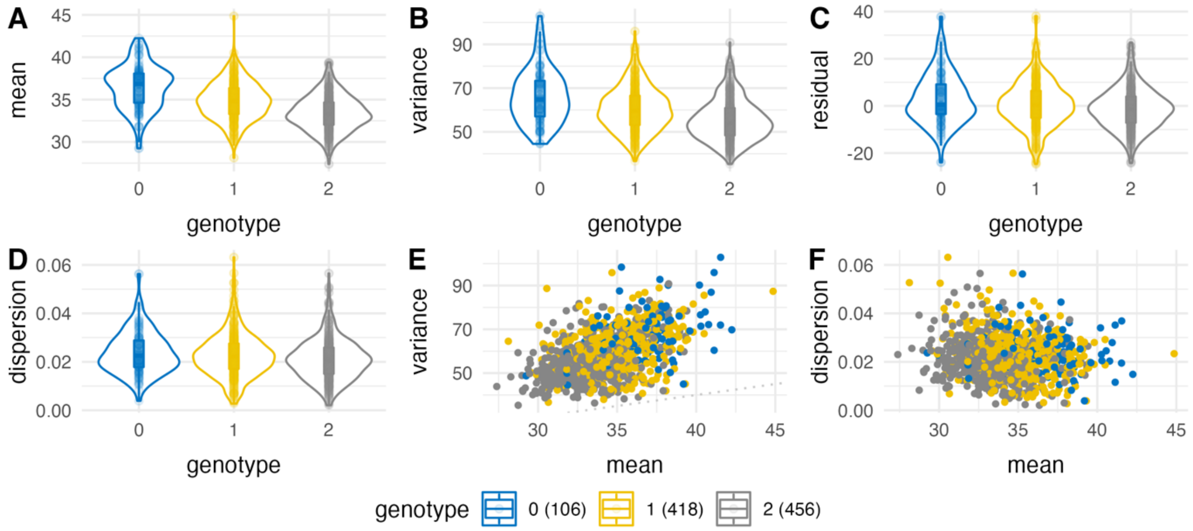


11:65644027_T_C ( rs583887 ) for *CTSW* in NK cells

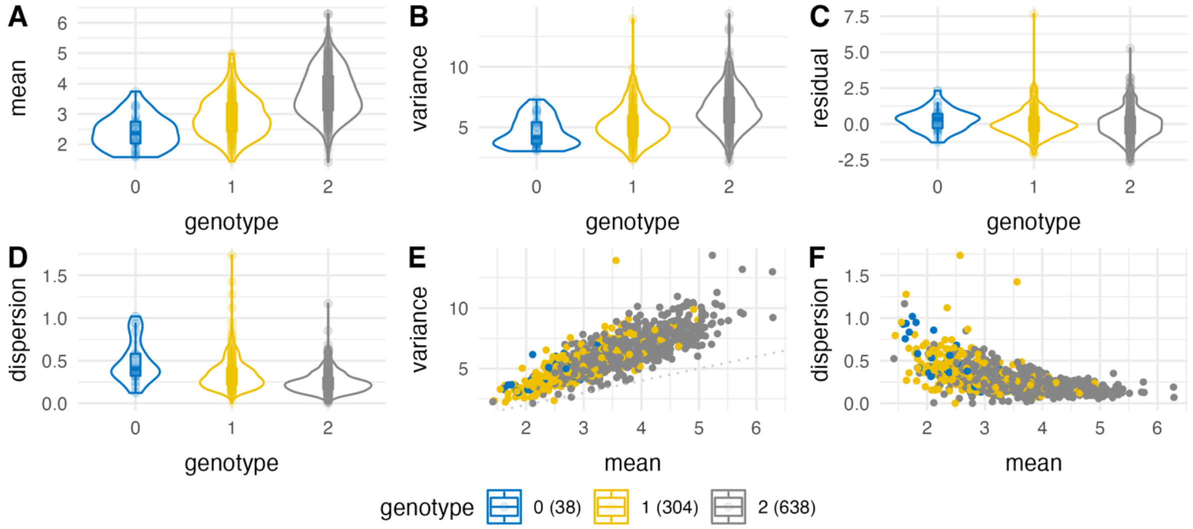

11:65645354_A_G ( rs596002 ) for *CTSW* in $CD8_{ET}$ cells

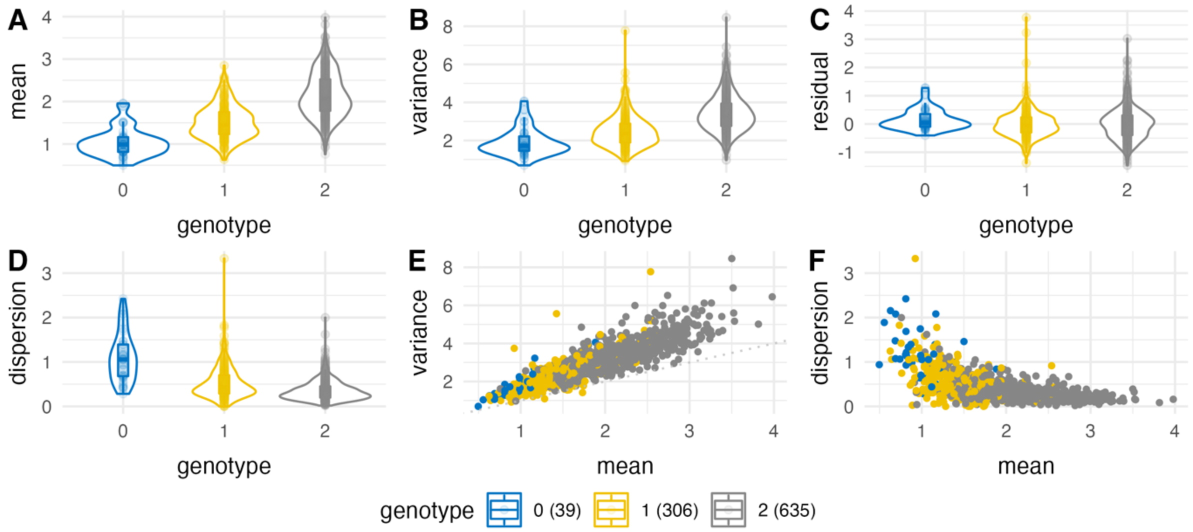


12:9623841_T_G ( rs10743738 ) for *KLRB1* in $CD4_{ET}$ cells

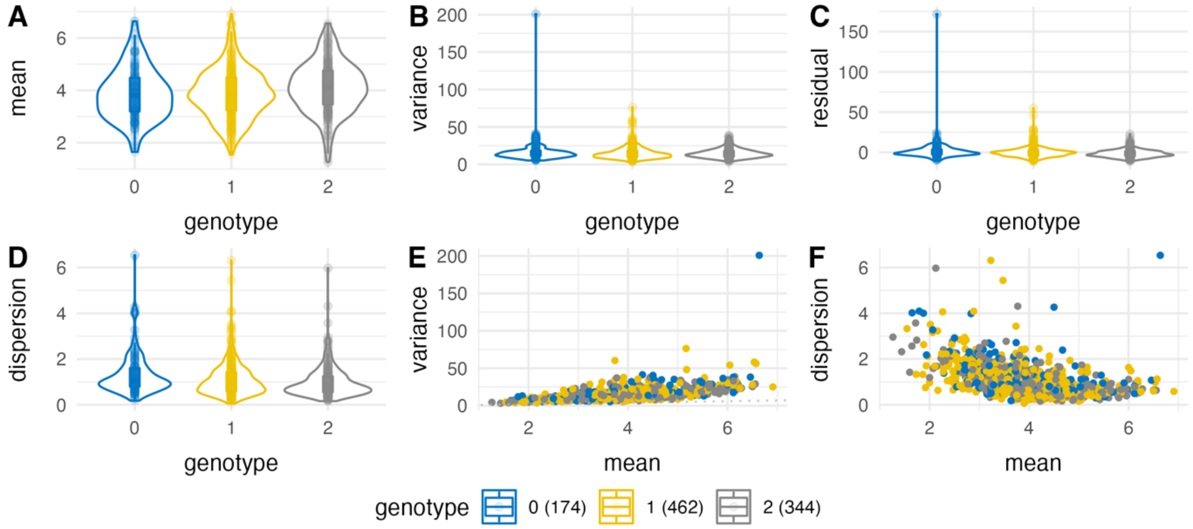


12:56435929_C_G ( rs1131017 ) for *RPS26* in $B_{IN}$ cells

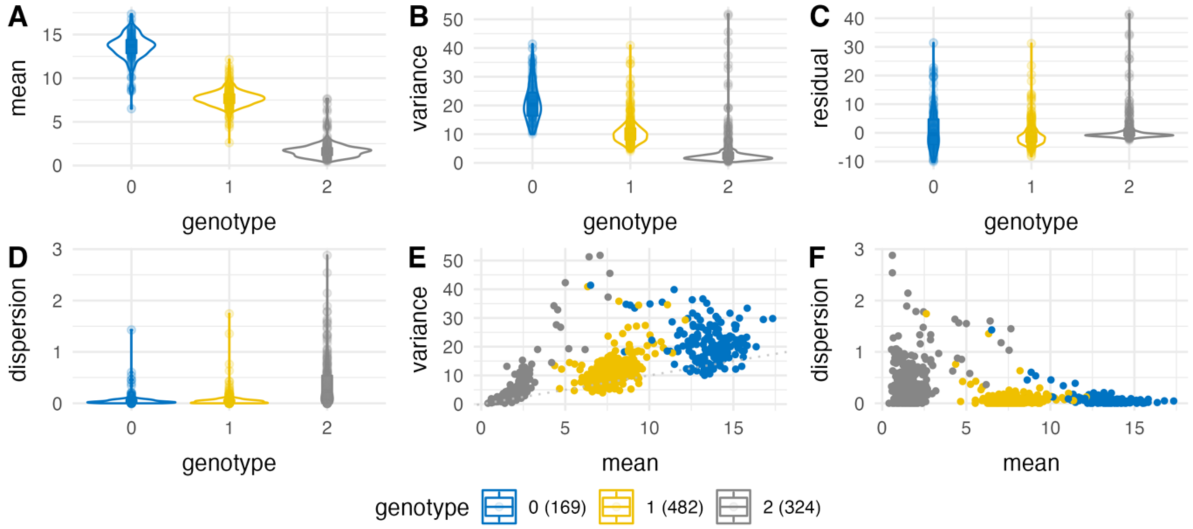

12:56435929_C_G ( rs1131017 ) for *RPS26* in $CD4_{ET}$ cells

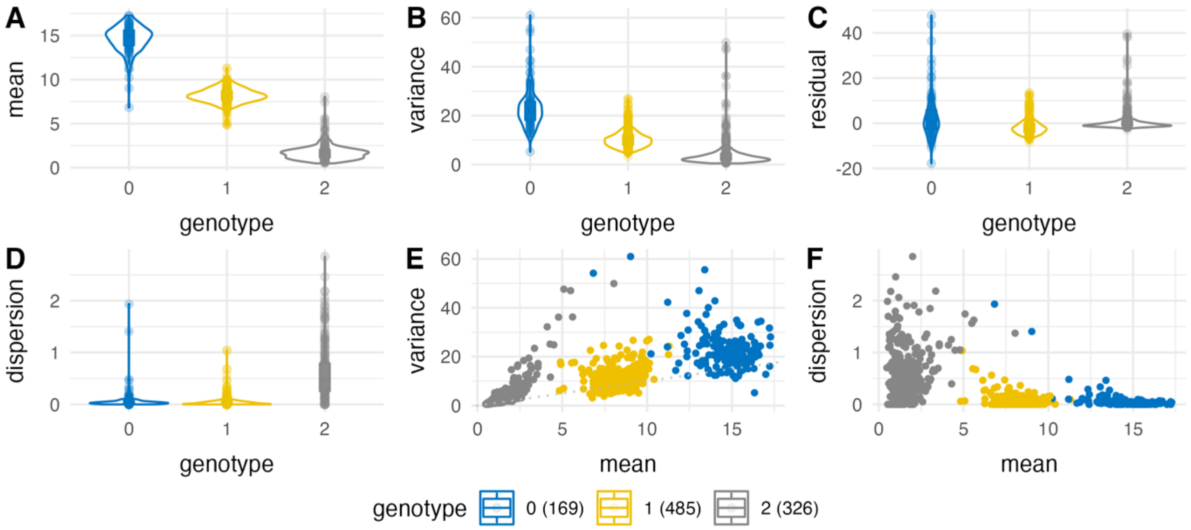


12:56435929_C_G ( rs1131017 ) for *RPS26* in $CD4_{NC}$ cells

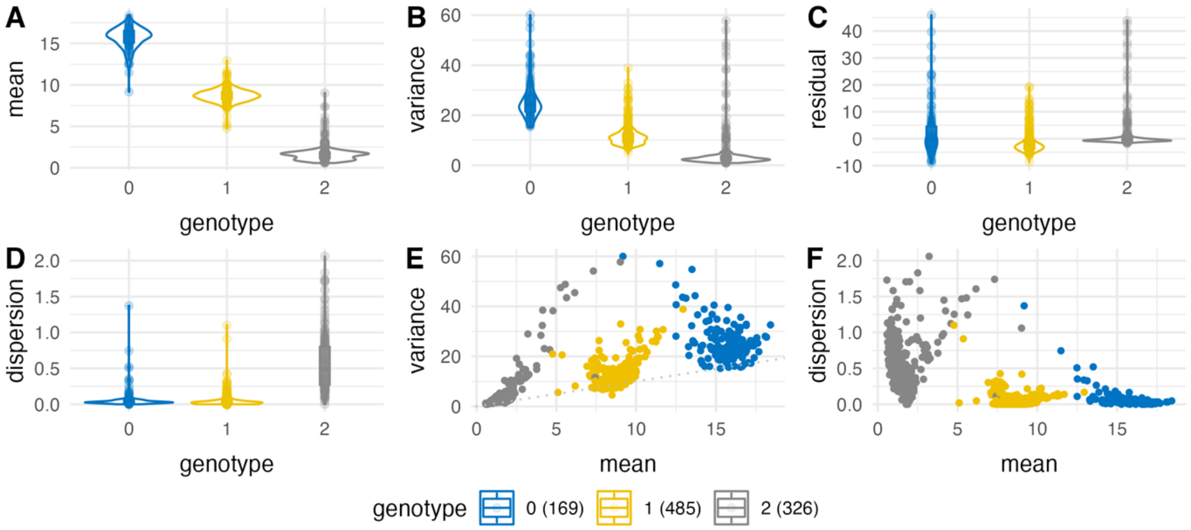


12:56435929_C_G ( rs1131017 ) for *RPS26* in $CD8_{ET}$ cells

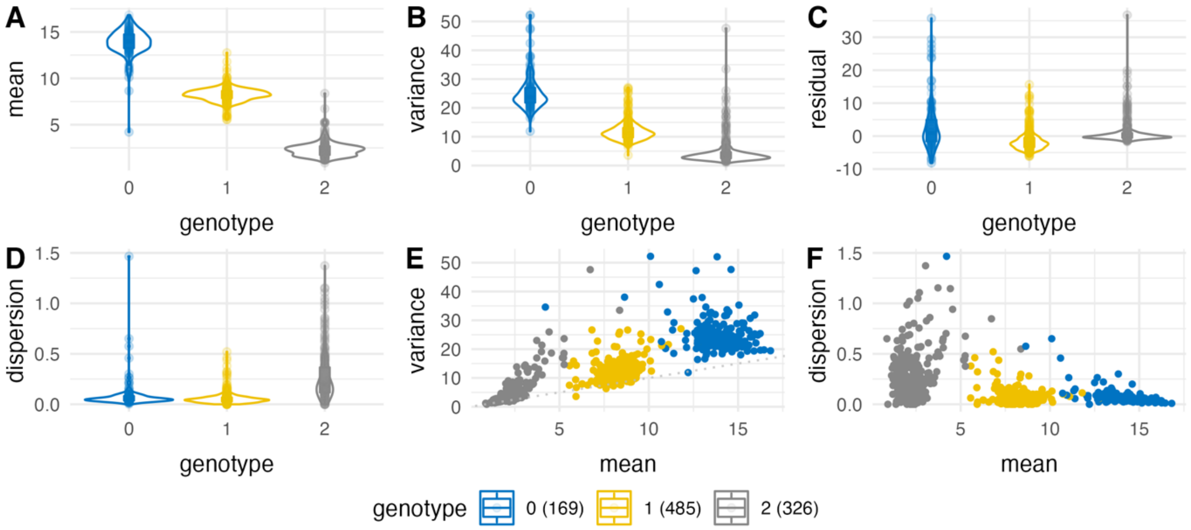

12:56435929_C_G ( rs1131017 ) for *RPS26* in $CD8_{NC}$ cells

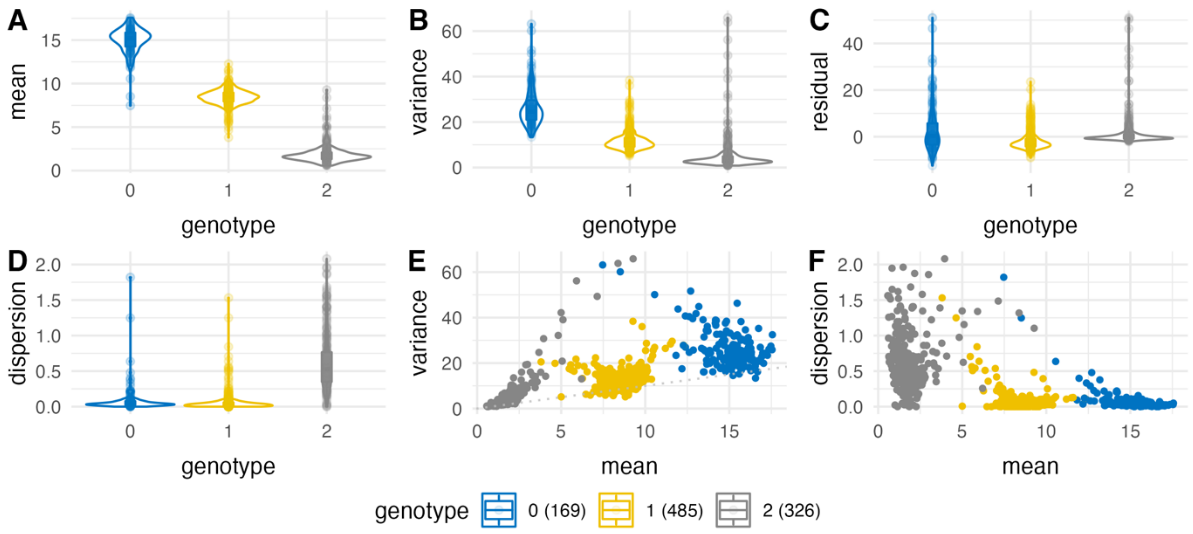


12:69747834_T_C ( rs1384 ) for *LYZ* in $Mono_{C}$ cells

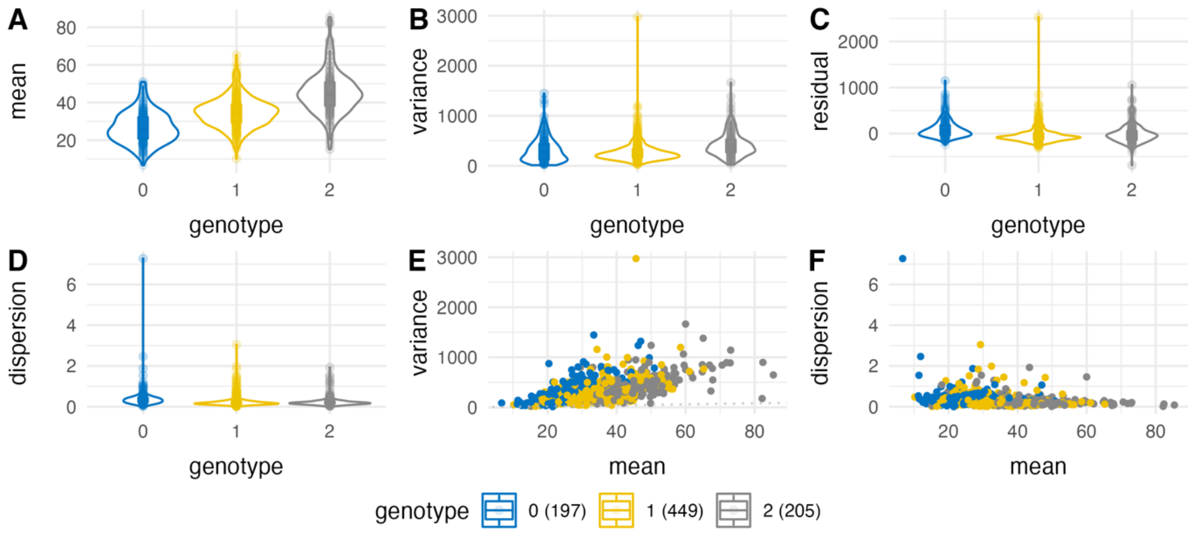


14:25083383_G_A ( rs11158812 ) for *GZMH* in NK cells

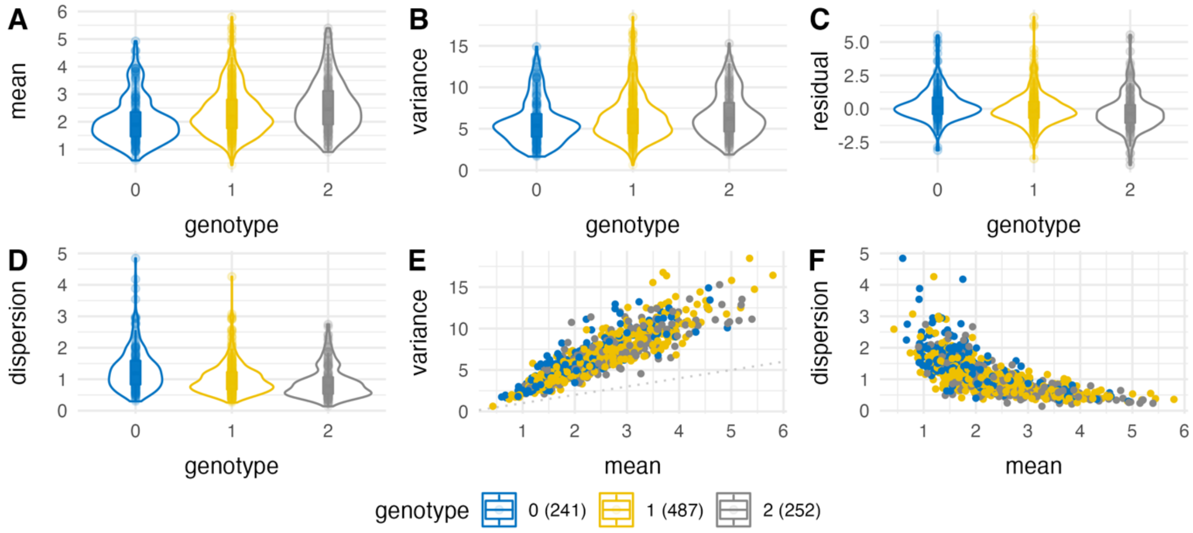

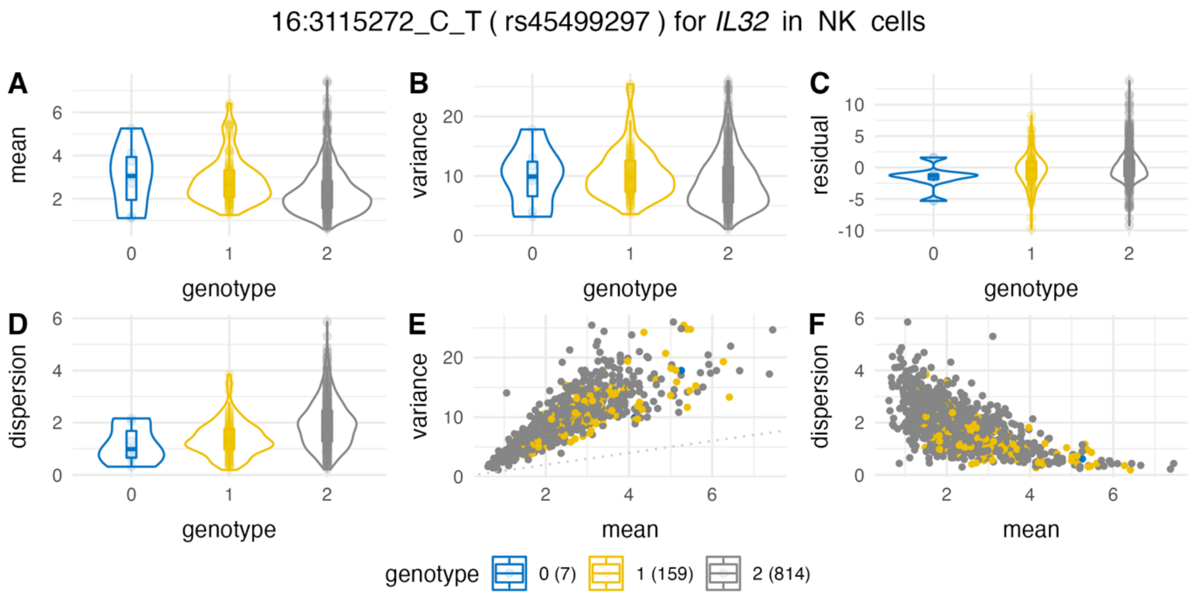
16:3115272_C_T ( rs45499297 ) for *IL32* in NK cells
A
B
C
D
E
F
mean
variance
residual
dispersion
genotype
mean
genotype 0 (7) 1 (159) 2 (814)

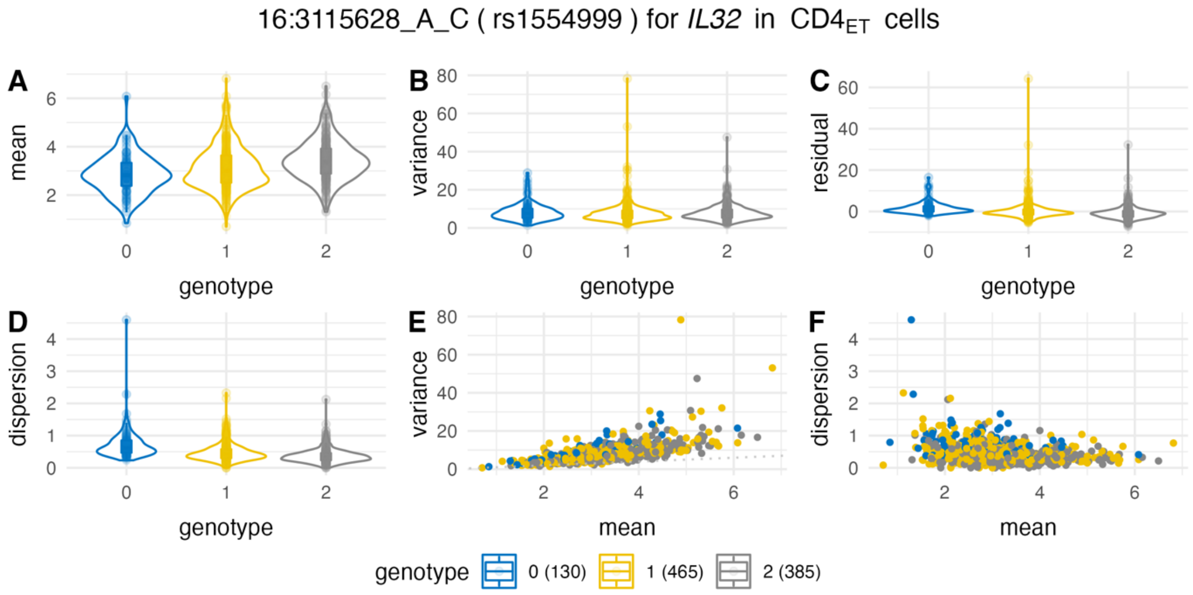
16:3115628_A_C ( rs1554999 ) for *IL32* in $CD4_{ET}$ cells
A
B
C
D
E
F
mean
variance
residual
dispersion
genotype
mean
genotype 0 (130) 1 (465) 2 (385)

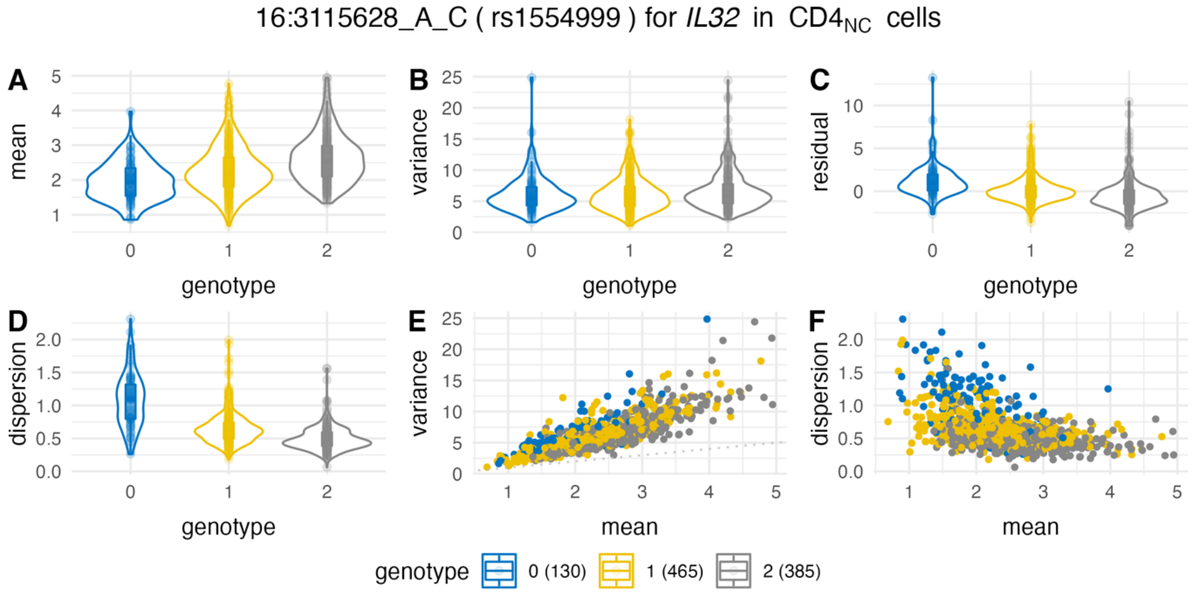
16:3115628_A_C ( rs1554999 ) for *IL32* in $CD4_{NC}$ cells
A
B
C
D
E
F
mean
variance
residual
dispersion
genotype
mean
genotype 0 (130) 1 (465) 2 (385)

16:3115628_A_C ( rs1554999 ) for *IL32* in $CD8_{ET}$ cells

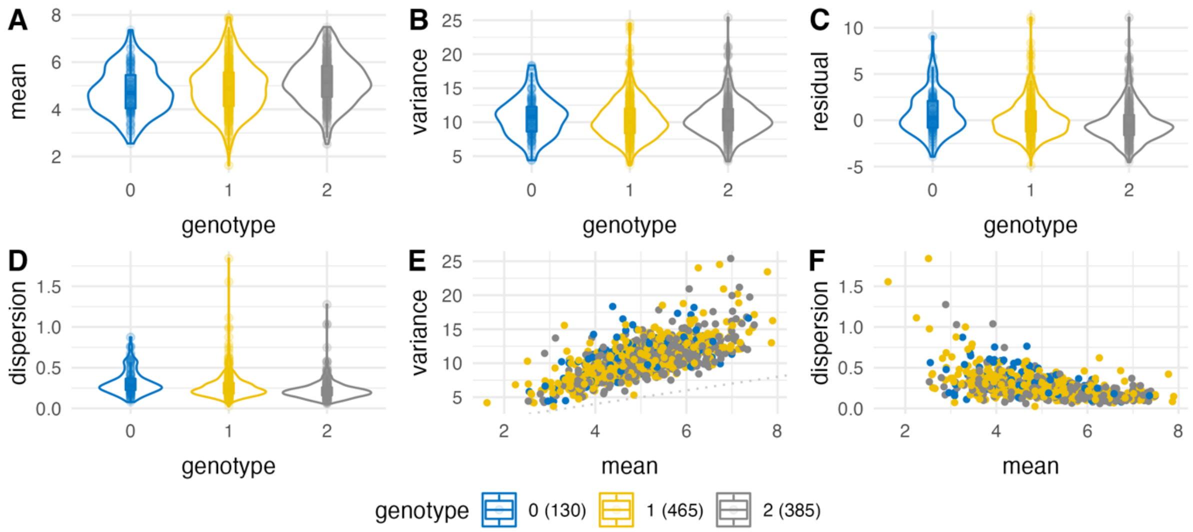


16:3115628_A_C ( rs1554999 ) for *IL32* in $CD8_{NC}$ cells

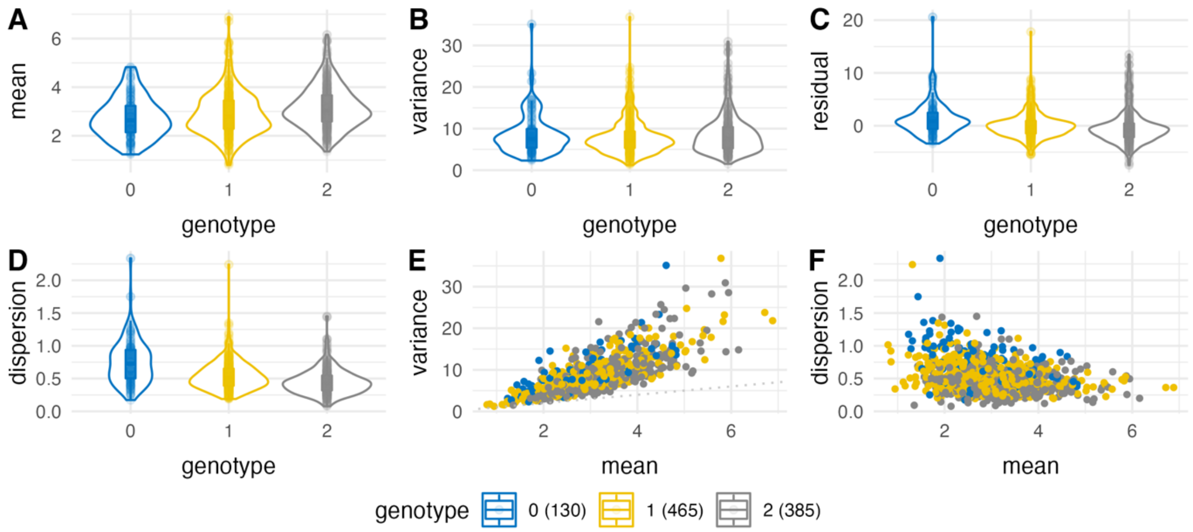


17:7207964_A_C ( rs7503161 ) for *EIF5A* in $CD4_{NC}$ cells

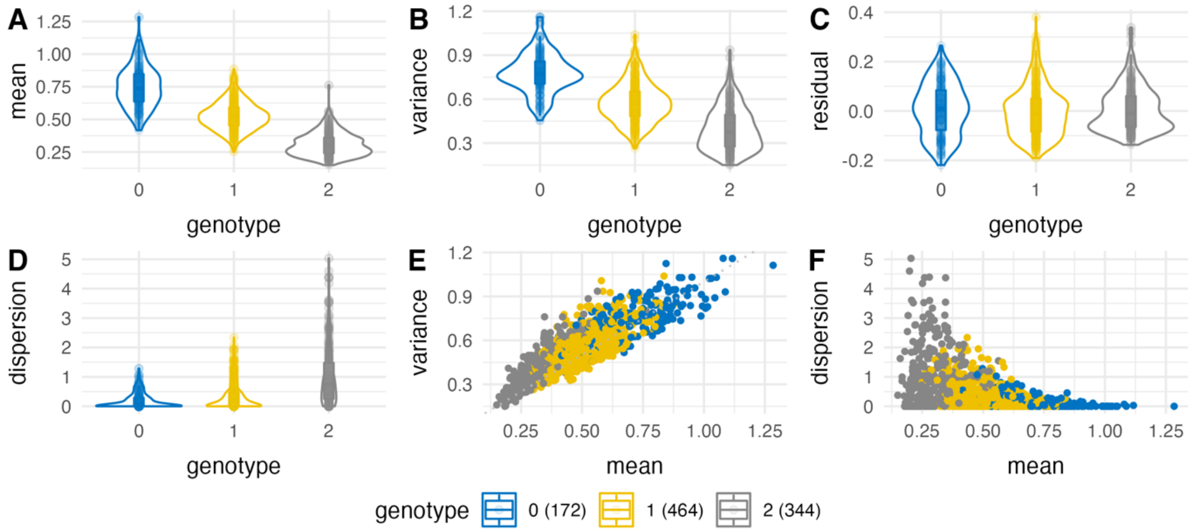

17:34397258_G_A ( rs854471 ) for *CCL3* in NK cells

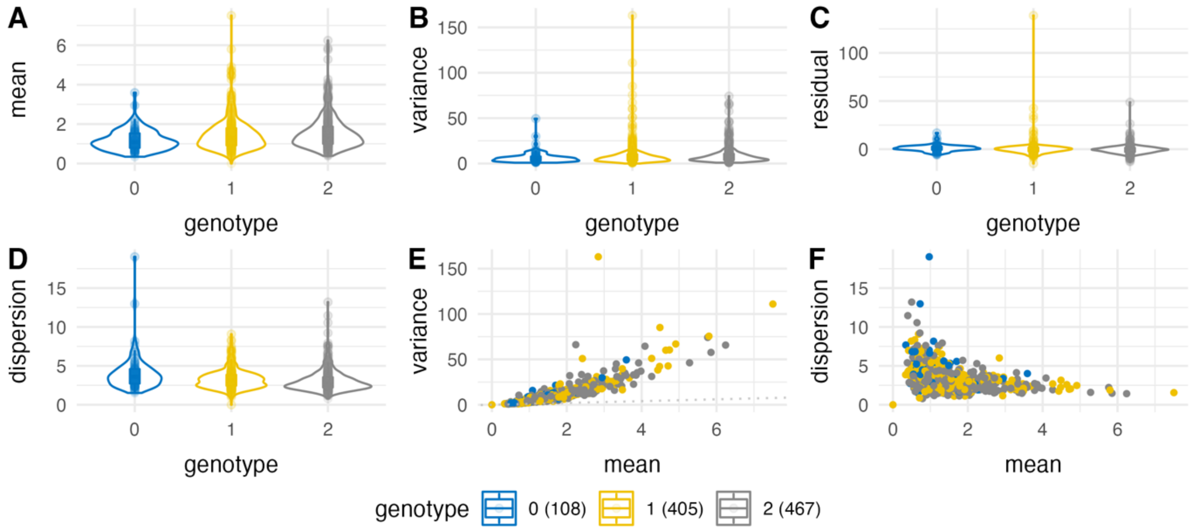


17:34411105_A_G ( rs1634490 ) for *CCL4* in $CD8_{ET}$ cells

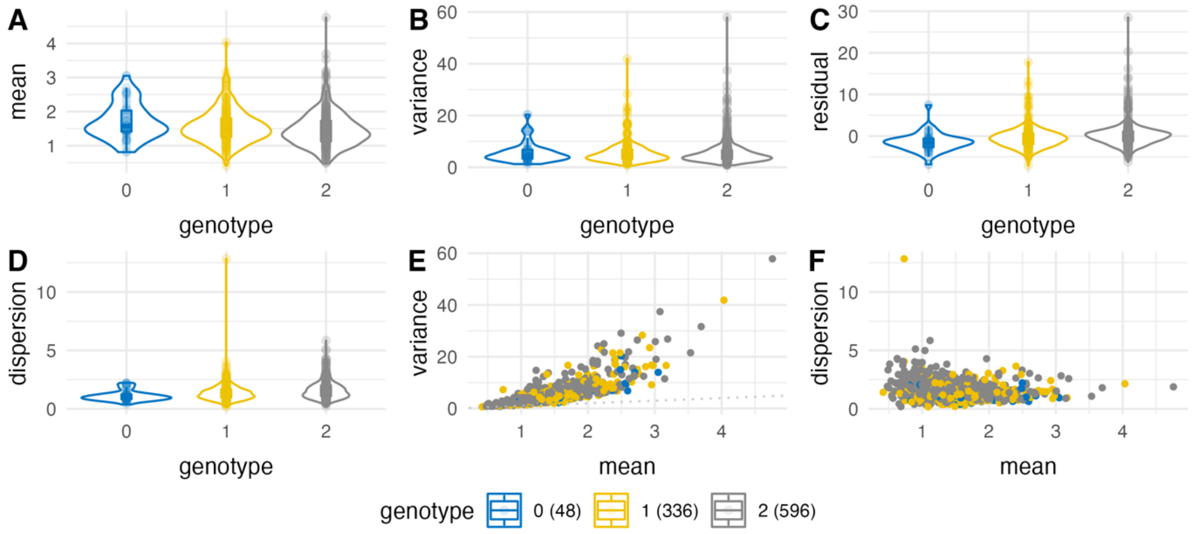


19:35658380_G_T ( rs12461097 ) for *FXYD5* in $CD4_{NC}$ cells

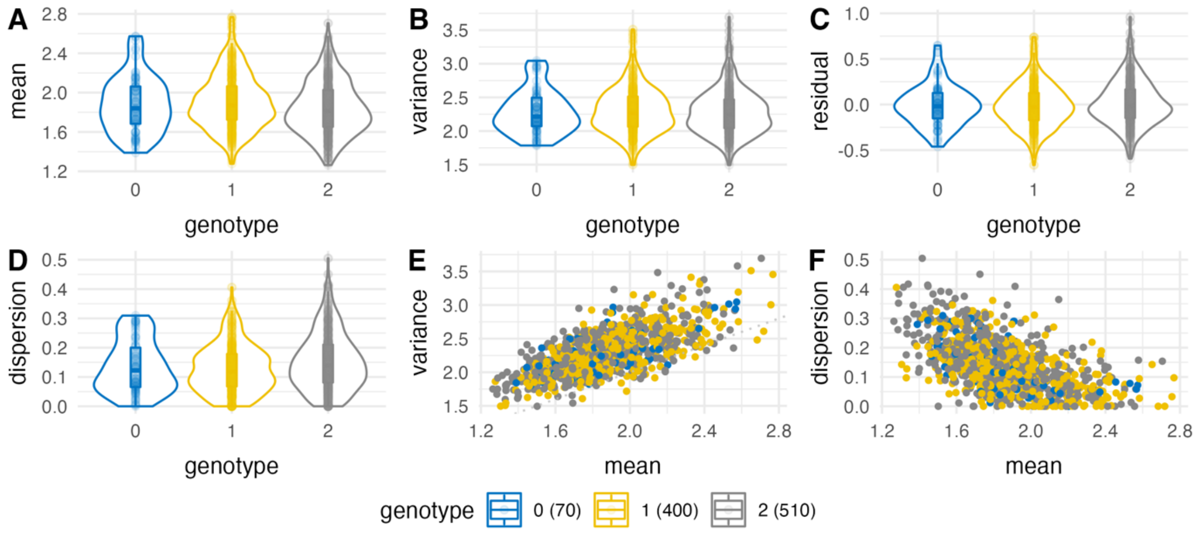

19:54700668_C_T ( rs34172242 ) for *RPS9* in CD4$_{NC}$ cells

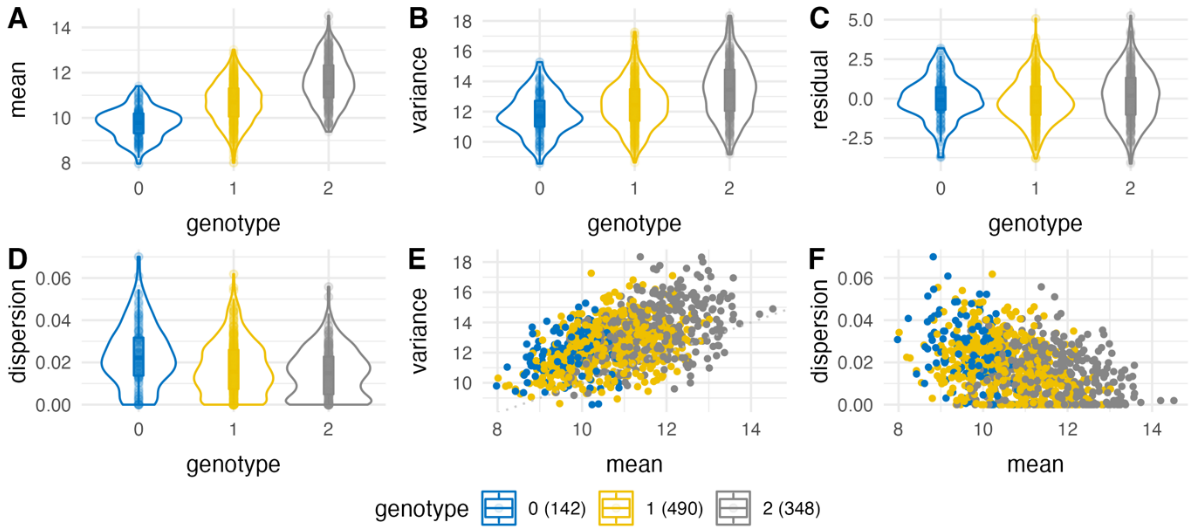


20:62152519_G_C ( rs72629024 ) for *PPDPF* in NK cells

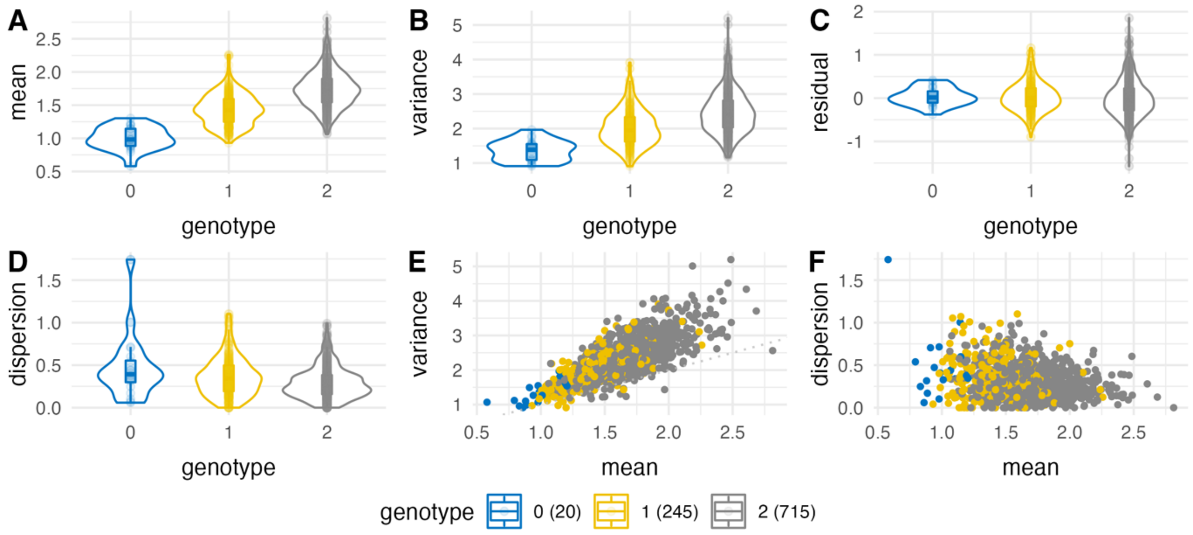


21:46328099_T_C ( rs760462 ) for *ITGB2* in CD8$_{ET}$ cells

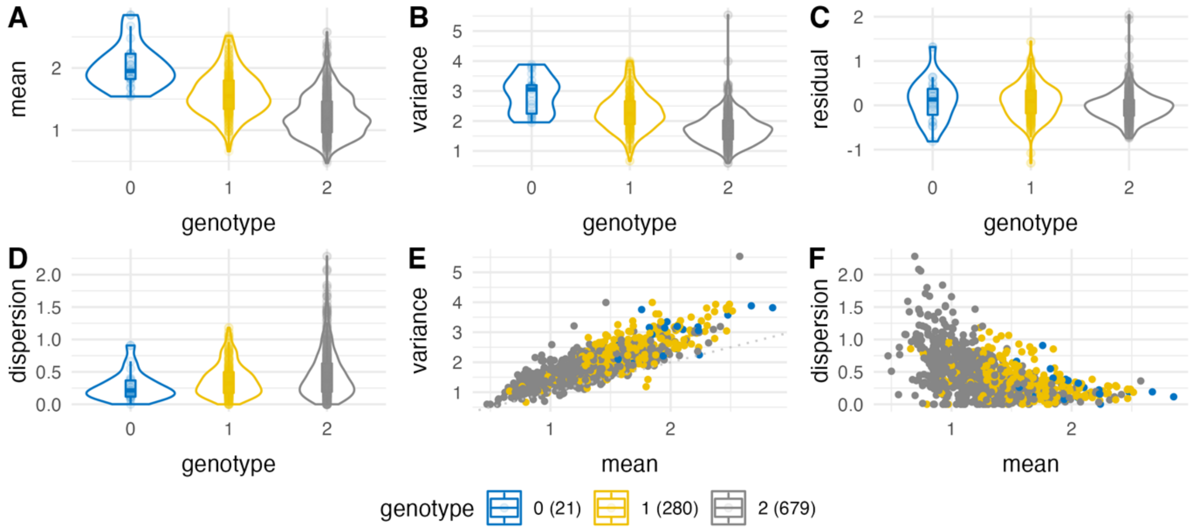

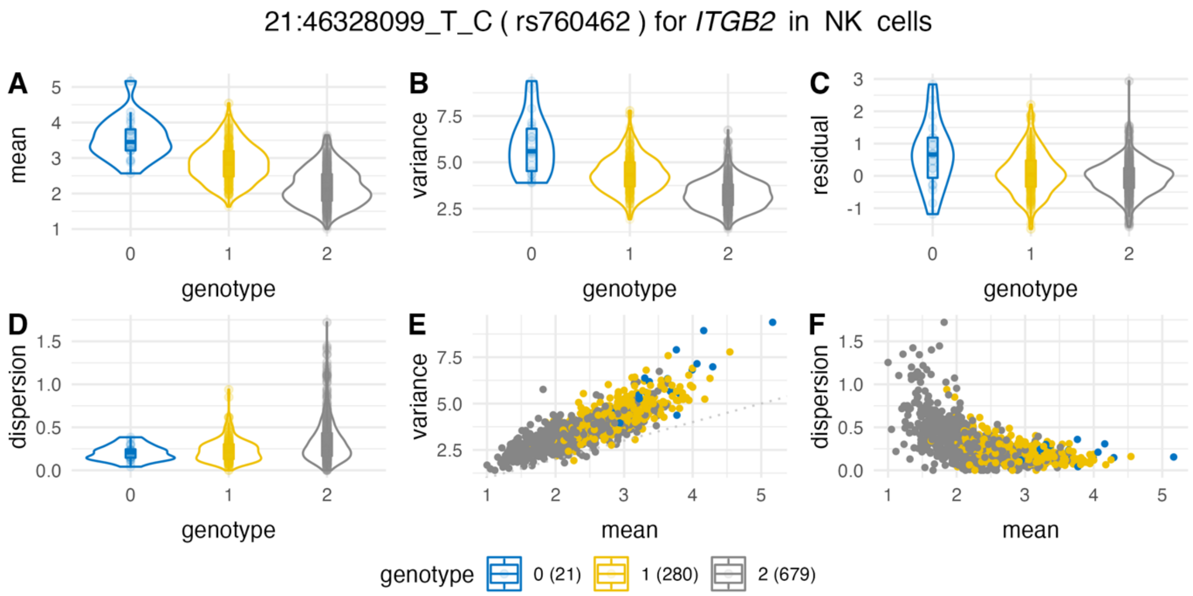


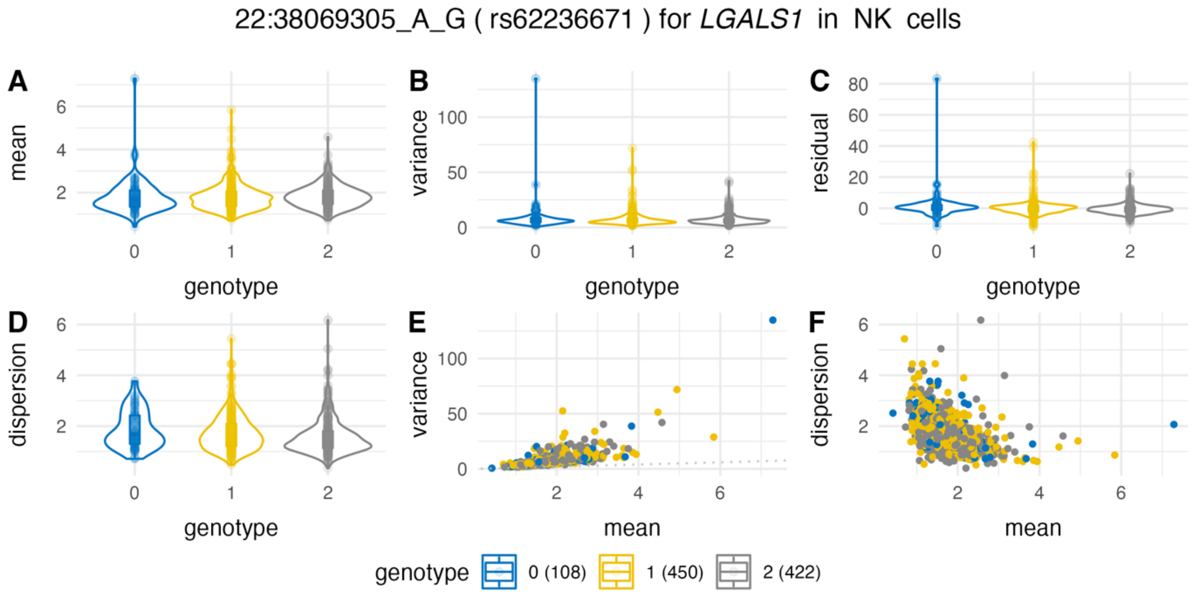


**Supplementary Figure 9. Genetic association plot for 55 deQTL-dGene pairs.**
**(A-D)** The violin plots of individual genotype of each deQTL corresponding to the intra-individual mean, variance, residual, dispersion of the dGene expression in certain cell types. The x-axis indicates the genotype (coded as 0, 1, 2 indicating number of alternative allele carried). **(E)** Scatter plot of intra-individual mean against intra-individual variance of expression. **(F)** Scatter plot of intra-individual mean against intra-individual dispersion of expression. The dispersion was estimated by CR-MLE method.

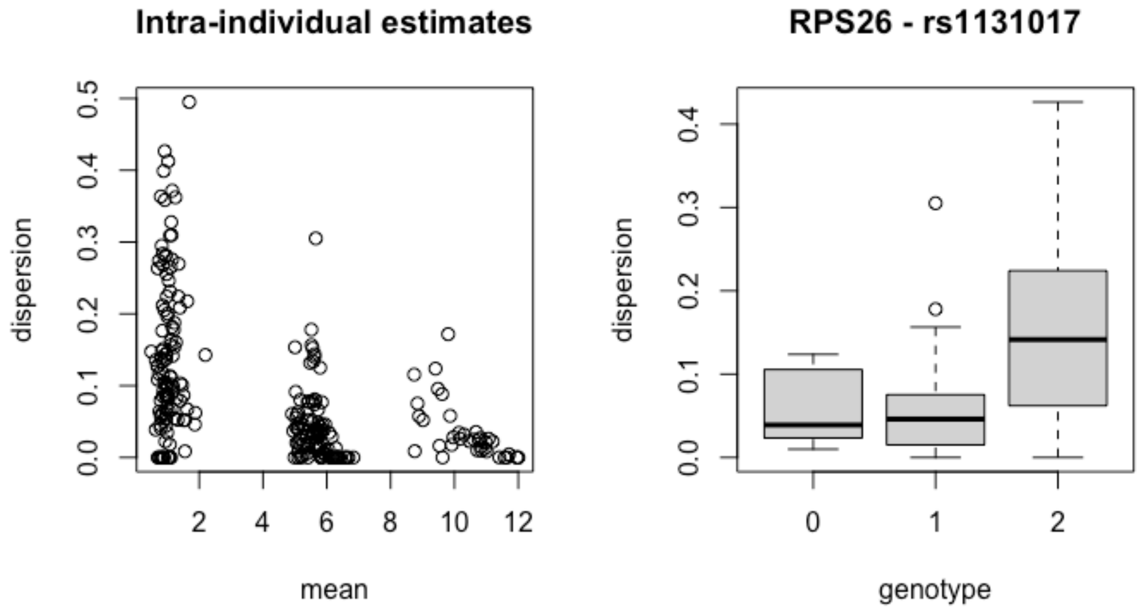


**Supplementary Figure 10. Mean-dispersion and genetic association plot for rs1131017-*RPS26* locus in CD4 T cells in Perez et al.** The left panel demonstrated the intra-individual estimates for mean and dispersion for *RPS26* in CD4 cells in the replication cohort. The right panel indicates the association pattern between genotype and intra-individual dispersion estimates.

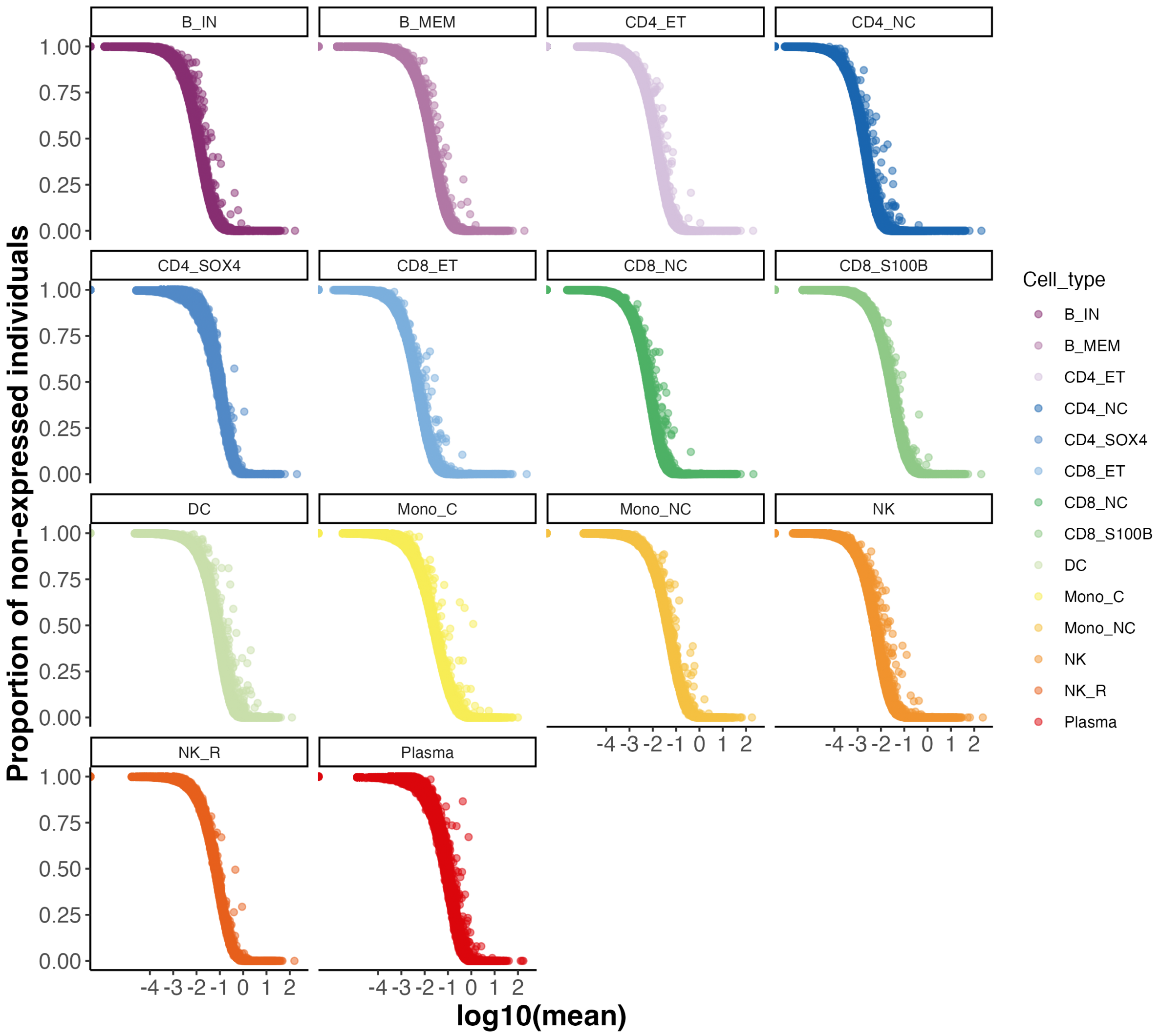


**Supplementary Figure 11. The relationship between the mean and the proportion of non-expression individuals.**

The scatter plot shows the relationship between the intra-individual mean and the proportion of non-expressed individuals per gene in 14 cell types. Each dot denotes one gene. The x-axis denotes the log10 transformed mean per gene. The y-axis denotes the proportion of individuals that do not express that gene.

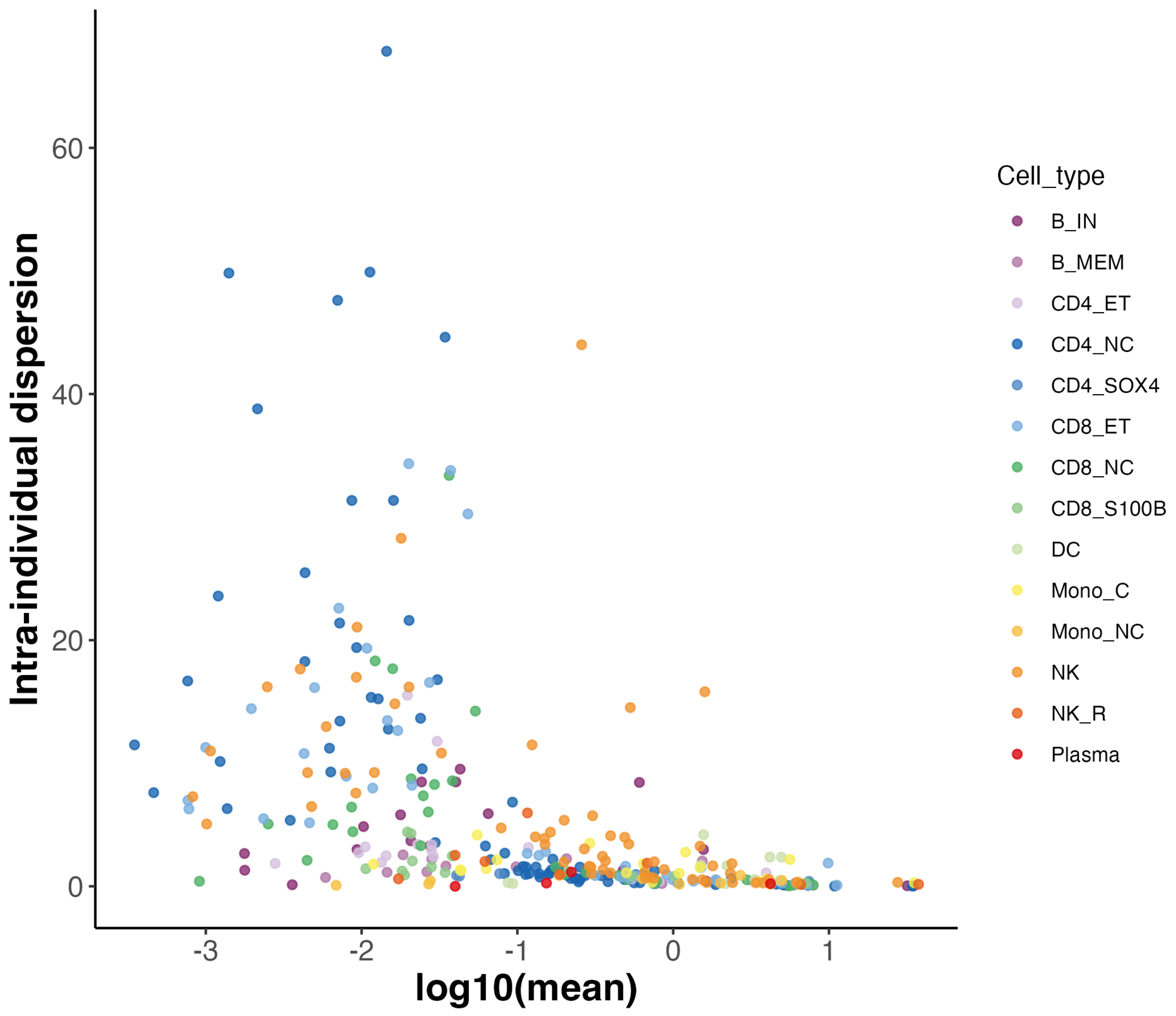


**Supplementary Figure 12. Distribution of dispersion estimates for 199 genes with 326 deQTLs in the preliminary test.**

This plot indicates the intra-individual estimates of mean and dispersion for 199 genes with 326 deQTL in 14 cell types in the preliminary test before filtering out the inflated genes. The x-axis indicates the log10 transformed intra-individual mean estimates and y-axis for the intra-individual dispersion estimates using Cox-Reid MLE method. The colour of the dot indicates the corresponding cell type.

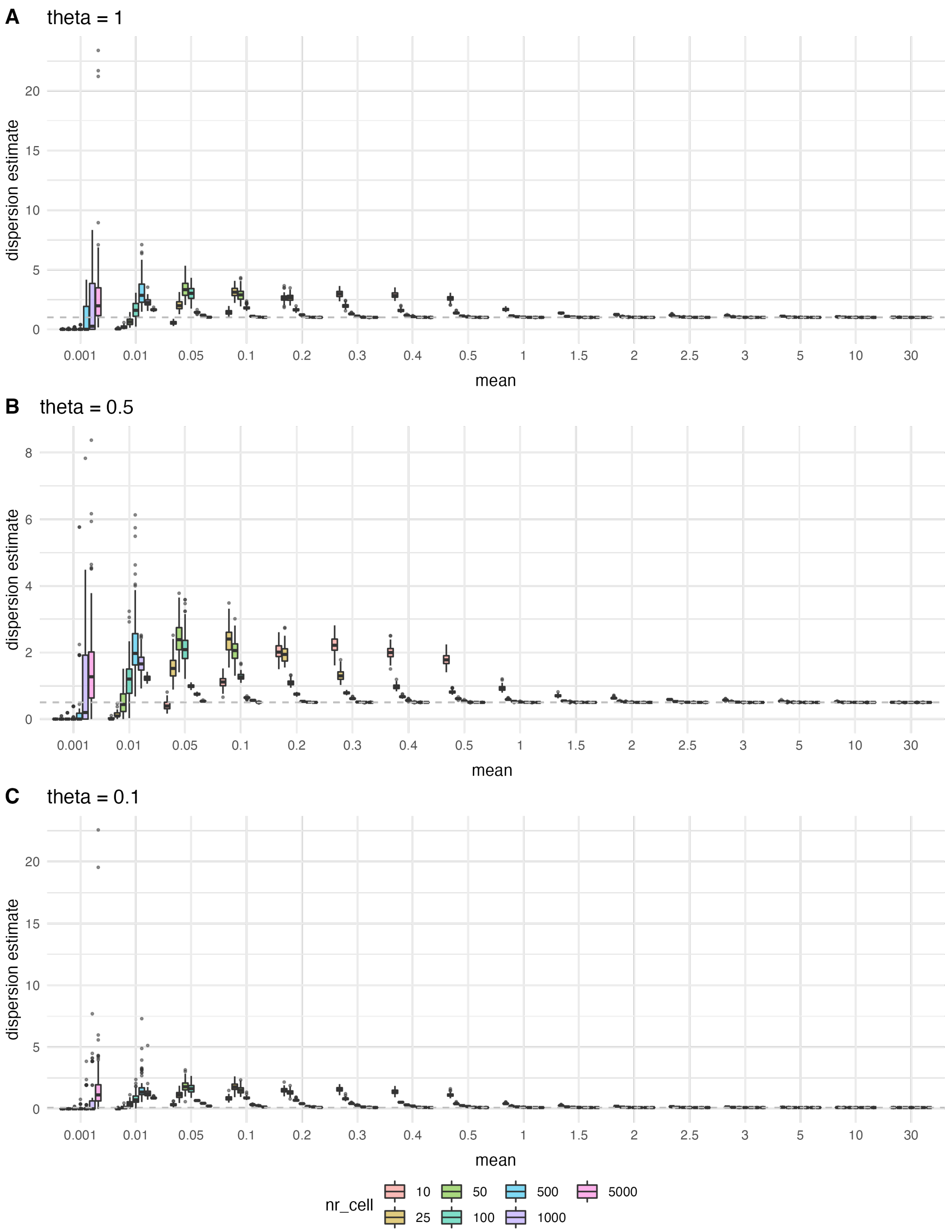


**Supplementary Figure 13. The CR-MLE estimates for intra-individual expression in simulation.**

The x-axis denotes the mean of the intra-individual expression, and the y-axis indicates the CR-MLE estimates. The colour group denotes five different settings for number of cells per individual. The horizontal grey dashed indicates the true dispersion parameter.

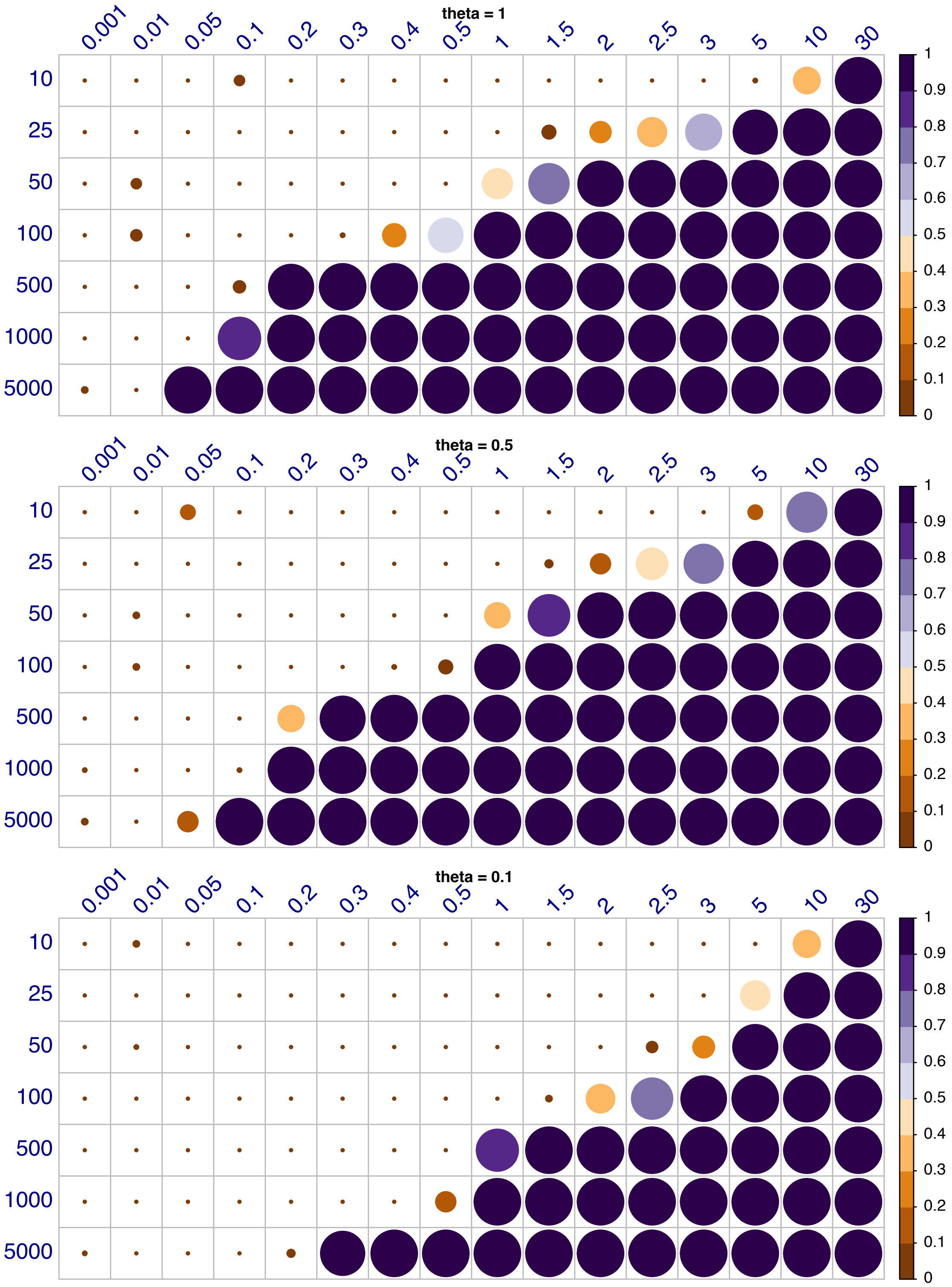


**Supplementary Figure 14. The accuracy of dispersion estimates in simulation.**
The heatmap plot demonstrates the accuracy of the dispersion estimates in the simulation under different scenarios. The row indicates the number of cells per individual, and the column indicates the mean of intra-individual expression. Each circle indicates the proportion of 100 replicates that the estimate is within $\pm 5\%$ of the true dispersion parameter.

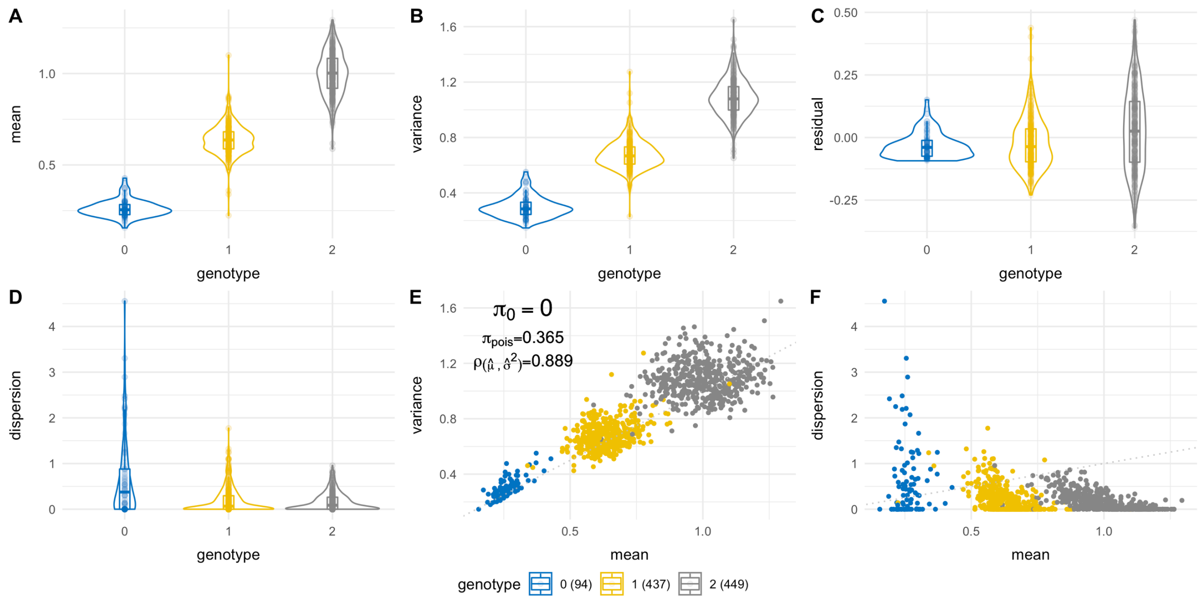


**Supplementary Figure 15. Example of inflated estimation of dispersion for *SMDT1* in $CD4_{NC}$ cells.**

The mean threshold for unbiased estimation of dispersion in $CD4_{NC}$ cells is 0.3. Although the mean of intra-individual mean for all individuals passed the threshold (0.3), the mean in the genotype 0 group (G/G of rs133379) did not, thus might suffer from inflated estimates.
